# The Existence and Uniqueness of Solutions in the Leaf Photosynthesis-Transpiration-Stomatal Conductance Model

**DOI:** 10.1101/2025.01.08.632031

**Authors:** Yuji Masutomi, Kazuhiko Kobayashi

## Abstract

The ***A***^***n***^-***E***-***g***^***s***^ model, which consistently describes leaf photosynthesis (***A***^***n***^), transpiration (***E***), and stomatal conductance (***g***^***s***^), is widely recognized and utilized as a “standard model” for quantifying these processes in terrestrial plants. However, since its proposal over 30 years ago, the model has faced a longstanding challenge: the “solution selection” problem, arising from the existence of multiple solutions with no guarantee that the obtained solution is correct.

In this study, we mathematically proved that the ***A***^***n***^-***E***-***g***^***s***^ model always has a unique solution satisfying the criteria ***g***^***s***^ ***>* 0** and ***C***^***i***^ ***>* 0**, where ***C***^***i***^ represents the CO^***2***^ concentration inside the leaf. This result establishes a rigorous mathematical theorem on the existence and uniqueness of solutions in the model. The theorem resolves the longstanding “solution selection” problem by enabling the unambiguous identification of the correct solution through selecting the unique solution that satisfies these criteria. Furthermore, the theorem ensures the validity of past estimations that satisfy these criteria and guarantees that future studies applying these criteria will yield correct estimations. These findings provide a robust mathematical foundation for the ***A***^***n***^-***E***-***g***^***s***^ model, reinforcing its role as the standard model for estimating leaf photosynthesis, transpiration, and stomatal conductance across diverse disciplines, from plant biology to climate science and beyond.

## 1 Introduction

Photosynthesis, transpiration, and stomata in terrestrial plants facilitate the exchange of carbon and water between the land surface and the atmosphere, playing crucial roles in global carbon and water cycles. According to Friedlingstein et al. (2022) [1], terrestrial plants absorb 130 PgC yr^−1^ of carbon through photosynthesis, which is 12.3 times higher than the 10.9 PgC yr^−1^ of carbon emissions caused by human activities, a major contributor to climate change. However, a nearly equivalent amount of carbon is released back into the atmosphere through respiration by plants, soil microbes, and other organisms. As a result, the net carbon uptake by land is only 2.4% (3.1 PgC yr^−1^) of the gross carbon absorption. Nevertheless, even this small net uptake mitigates atmospheric CO_2_ increases, slowing climate change and its impacts [2, 3]. In crops, photosynthesis directly correlates with food production. Even slight changes can impact food security in vulnerable countries, and climate change is expected to significantly affect crop production [4–7]. Additionally, most terrestrial water transport to the atmosphere occurs through transpiration facilitated by stomata [8, 9]. With the increasing frequency of water-related disasters such as floods and droughts driven by climate change [10], accurate comprehension of transpiration and stomatal conductance, key factors in the water cycle, is critical for addressing these challenges. Thus, precise estimation of photosynthesis, transpiration, and stomatal conductance is fundamental for understanding global carbon and water cycles, climate change, and its impacts.

The Leaf Photosynthesis (*A*_*n*_)-Transpiration (*E*)-Stomatal Conductance (*g*_*s*_) model, first proposed by Collatz et al. (1991: [11]), is widely used as a standard model for quantitatively estimating plant photosynthesis, transpiration, and stomatal conductance. It has been applied not only in ecosystem and vegetation models ([12–15]) but also in crop, hydrological, land surface, climate, and earth system models ([16– 23]). For instance, out of the 16 ecosystem models employed in the “Global Carbon Project” by researchers worldwide to estimate Earth’s carbon balance ([1]), 10 models adopt the *A*_*n*_-*E*-*g*_*s*_ model and its variant forms. Among the remaining six models, four use simplified versions of the *A*_*n*_-*E*-*g*_*s*_ model. However, a significant issue of “solution selection” has persisted with this *A*_*n*_-*E*-*g*_*s*_ model for over 30 years since its proposal. This problem stems from the fact that the *A*_*n*_-*E*-*g*_*s*_ model reduces to a fifth-degree equation concerning photosynthesis or leaf internal CO_2_ concentration, resulting that there exist multiple solutions to the model (Longo et al., 2019: [14]; Masutomi, 2023: [24]). Due to the existence of multiple solutions, it is uncertain whether a single solution obtained is correct. In other words, the obtained solution might be incorrect, considering the existence of other potential solutions. Despite the extensive use of this model in numerous studies, including the “Global Carbon Project,” the values obtained in these studies might not be correct, leading to the possibility that the conclusions and implications drawn from them could also be incorrect.

Several studies have explored the issue of “solution selection” in the *A*_*n*_-*E*-*g*_*s*_ model. Baldocchi (1994) ([25]) investigated a simplified version of the model in which the equations for transpiration were omitted, and leaf humidity was provided externally. The study demonstrated that this simplified model could be reduced to a cubic equation for the photosynthesis rate, allowing the solutions to be expressed as three analytical solutions of the cubic equation. Additionally, it concluded that one specific analytical solution among these three was always appropriate, suggesting that the problem of “solution selection” for the simplified model appeared to be resolved. However, Masutomi (2023) ([24]) extended the analysis of this simplified model to a broader range of environmental conditions and showed numerically that the biologically and physically appropriate analytical solution, defined as the solution satisfying the criteria *g*_*s*_ *>* 0 and *C*_*i*_ *>* 0 (internal CO_2_ concentration), varied among the three solutions depending on environmental conditions. Furthermore, it was numerically proven that only one biologically and physically appropriate solution always exists within the investigated range of environmental conditions. This finding provided numerical evidence for the existence and uniqueness of biologically and physically appropriate solutions in the simplified model. Since there is always only one such solution, the problem of “solution selection” is effectively resolved for the simplified model. In contrast, Longo et al. (2019) ([14]) proposed an algorithm to obtain a single solution from the full *A*_*n*_-*E*-*g*_*s*_ model by analyzing the singularities of functions derived from the model. However, the correctness of the solution obtained using their algorithm has not been rigorously proven, leaving open the possibility that other solutions might be correct. Consequently, the problem of “solution selection” remains unresolved for the *A*_*n*_-*E*-*g*_*s*_ model.

The objective of this study is to resolve the longstanding problem of “solution selection” in the *A*_*n*_-*E*-*g*_*s*_ model for C3 plants. This is achieved by analytically demonstrating the existence and uniqueness of biologically and physically appropriate solutions in the *A*_*n*_-*E*-*g*_*s*_ model. The proof employed in this study is inspired by singularity analysis used in Longo et al. (2019) ([14]). Singularity of functions strongly characterizes the geometry of the functions and, when combined with other geometric features such as zeros and derivatives, can fully define the geometry of these functions. In this study, by analytically examining such geometric features of the functions derived from the *A*_*n*_-*E*-*g*_*s*_ model, the existence and uniqueness of biologically and physically appropriate solutions are ultimately proven.

## 2 Theorem

In this paper, we establish the following theorem on the existence and uniqueness of biologically and physically appropriate solutions to the *A*_*n*_-*E*-*g*_*s*_ model. The description of the model is given in Section 3.

**Theorem** (Existence and Uniqueness of Biologically and Physically Appropriate Solutions for the *A*_*n*_-*E*-*g*_*s*_ Model). *For the A*_*n*_*-E-g*_*s*_ *model, there always exists a unique biologically and physically appropriate solution*.

A solution is considered *biologically and physically appropriate* if it satisfies the following criteria:

**Criteria** (Criteria for Biologically and Physically Appropriate Solutions). *A solution must satisfy g*_*s*_ *>* 0 *(stomatal conductance) and C*_*i*_ *>* 0 *(CO*_2_ *concentration inside the leaf)*.

The *A*_*n*_-*E*-*g*_*s*_ model may produce multiple solutions for variables such as leaf photosynthesis (*A*_*n*_), stomatal conductance (*g*_*s*_), and internal CO_2_ concentration (*C*_*i*_) [14, 24]. However, some of these solutions result in negative values for *g*_*s*_ and/or *C*_*i*_. Since *g*_*s*_ and *C*_*i*_ represent biological and physical quantities, they must remain positive to be meaningful [24]. Therefore, only solutions satisfying these criteria are classified as *biologically and physically appropriate*.

In the following sections, the term “appropriate” exclusively refers to solutions that satisfy the criteria of “biologically and physically appropriate.”

## 3 Methods

### 3.1 The *A*_*n*_-*E*-*g*_*s*_ Model

The *A*_*n*_-*E*-*g*_*s*_ model explored in this study is based on the formulation proposed by Collatz et al. (1991) [11]. In this model, the net photosynthesis rate, *A*_*n*_(*C*_*i*_) [mol(CO_2_) m^−2^ s^−1^], follows the framework of Farquhar et al. (1980) [26] and is given by:

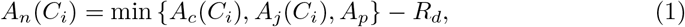

where

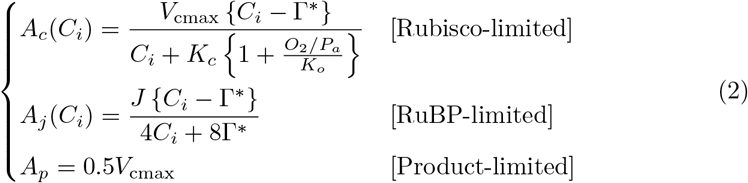

Here, *A*_*c*_(*C*_*i*_), *A*_*j*_(*C*_*i*_), and *A*_*p*_ [mol(CO_2_) m^−2^ s^−1^] represent Rubisco-limited, RuBP-limited, and Product-limited photosynthesis rate, respectively. *R*_*d*_ [mol(CO_2_) m^−2^ s^−1^] is the rate of dark respiration; *V*_cmax_ [mol m^−2^ s^−1^] is the maximum rate of carboxylation; *J* [mol m^−2^ s^−1^] is the electron transport rate; *C*_*i*_ [mol mol^−1^] is the internal leaf CO_2_ concentration; *O*_2_ [Pa] is the atmospheric partial pressure of oxygen; *P*_*a*_ [Pa] is the surface air pressure; Γ^*^ [mol mol^−1^] is the CO_2_ compensation point where the rate of CO_2_ uptake by carboxylation is balanced by the rate of photorespiratory CO_2_ release; and *K*_*c*_ and *K*_*o*_ [mol mol^−1^] are the Michaelis-Menten constants for CO_2_ and O_2_, respectively. The mathematical notations used in this study is detailed in the Appendix.

A variety of models have been proposed for stomatal conductance during daytime for water vapor [27]. In this study, we adopt Medlyn’s model for daytime conditions (*A*_*n*_(*C*_*i*_) *>* 0)[28], which is used in recent land surface models [15, 29]. For nighttime conditions (*A*_*n*_(*C*_*i*_) ≤ 0), *g*_*s*_ is assumed to be at its minimum. Therefore, stomatal conductance *g*_*s*_ [mol m^−2^ s^−1^] is given by:

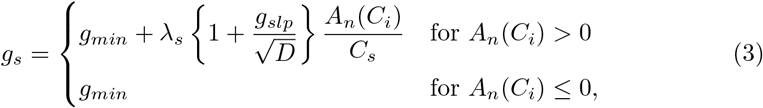

where *C*_*s*_ [mol mol^−1^] is the CO_2_ concentration at the leaf surface; *λ*_*s*_ [-] is the ratio of the diffusivities of CO_2_ and water vapor through stomata, fixed at 1.6; *g*_*min*_ [mol m^−2^ s^−1^] is the minimum stomatal conductance, and *g*_*slp*_ [Pa^1*/*2^] represents the sensitivity of *g*_*s*_ to variables such as *A*_*n*_(*C*_*i*_), *C*_*s*_, and *D*. Both *g*_*min*_ and *g*_*slp*_ are plant-specific parameters. *D* [Pa] is the vapor pressure deficit at the leaf surface and is given by:

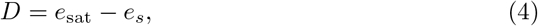

where *e*_*s*_ [Pa] represents the vapor pressure at the leaf surface, while *e*_sat_ [Pa] represents the saturated vapor pressure at the leaf surface and inside the leaf. Note that the equation for *g*_*s*_ (Eq. (3)) is mathematically undefined at *D* = 0 because *g*_*s*_ diverges at this point. Therefore, the case of *D* = 0 requires special treatment in the analysis.

The fluxes of CO_2_ and water vapor between the leaf and the environment are governed by standard mass transport relationships [30]. For CO_2_ fluxes, we have:

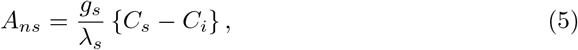

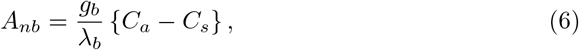

where *A*_*ns*_ and *A*_*nb*_ are the CO_2_ fluxes from the leaf surface to the leaf interior and from the atmosphere to the leaf surface, respectively; *C*_*a*_ [mol mol^−1^] is the atmospheric CO_2_ concentration; *g*_*b*_ [mol m^−2^ s^−1^] is the conductance of the leaf boundary layer for water vapor; and *λ*_*b*_ [-] is the ratio of the diffusivities of CO_2_ and water vapor at the leaf surface, fixed at 1.4. Under steady-state conditions for *C*_*i*_ and *C*_*s*_, the fluxes *A*_*ns*_ and *A*_*nb*_ become:

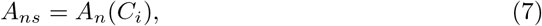

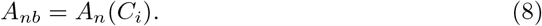

Similarly, the transpiration rate is related to *g*_*s*_ and *g*_*b*_ as follows:

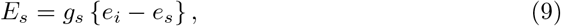

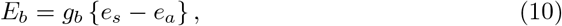

where *E*_*s*_ and *E*_*b*_ [mol(H_2_O) m^−2^ s^−1^] are the transpiration rates from the leaf interior to the surface, and from the surface to the surrounding air, respectively; and *e*_*a*_ and *e*_*i*_ [Pa] represent the vapor pressures in the air and inside the leaf, respectively. *e*_*i*_ is assumed to be saturated and is expressed as *e*_*i*_ = *e*_sat_. If *e*_*i*_ *< e*_*a*_ (and thus *e*_*s*_ *< e*_*a*_), vapor flux occurs from the atmosphere to the leaf surface, causing *e*_*s*_ to increase until it reaches saturation, i.e., *e*_*s*_ = *e*_sat_ = *e*_*i*_, resulting in *D* = 0 (Eq. (4)). Consequently, special treatment is required for this case because *g*_*s*_ diverges when *D* = 0. Conversely, when *e*_*i*_ *> e*_*a*_, vapor flux occurs outward from the leaf interior. Under steady-state conditions for *e*_*s*_, *E*_*b*_ = *E*_*s*_ is achieved. Thus, we have:

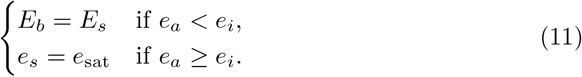

The *A*_*n*_-*E*-*g*_*s*_ model comprises 11 equations (Eqs. (1) to (11)) and 11 unknowns: *A*_*n*_, *A*_*c,j,p*_, *A*_*ns*_, *A*_*nb*_, *E*_*b*_, *E*_*s*_, *C*_*i*_, *C*_*s*_, *e*_*s*_, *D*, and *g*_*s*_. Consequently, these unknowns can be determined internally through the model equations and are referred to as *internal variables*. Other variables, including plant-specific parameters, are provided externally and are termed *external variables*. Table 1 presents all the internal and external variables in the *A*_*n*_-*E*-*g*_*s*_ model, categorizing them based on their possible ranges.

**Table 1:**
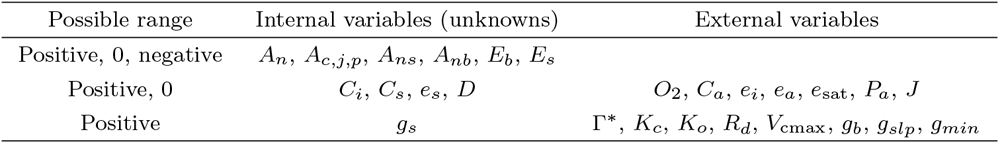
Internal and external variables.

Flux variables, such as *A*_*n*_, *A*_*c,j,p*_, *A*_*ns*_, *A*_*nb*_, *E*_*s*_, and *E*_*b*_, can take positive, negative, or zero values. The remaining variables must be non-negative, as they represent biological or physical quantities. Furthermore, the non-negative variables are divided into two categories: those that can be zero and those that cannot. We assume that physical variables can reach zero, whereas biological ones generally cannot, except for *J*, which can be zero under conditions such as darkness (e.g., at night).

### 3.2 Net Photosynthesis Rate *A*_*n*_(*C*_*i*_)

The expression for the net photosynthesis rate, *A*_*n*_(*C*_*i*_), in Eq. (1) can be reformulated by combining it with Eq. (2) as follows:

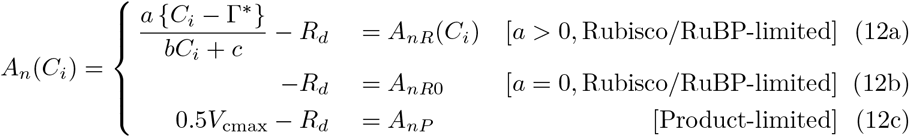

where the constants *a, b*, and *c* in Eq. (12a) are defined as:

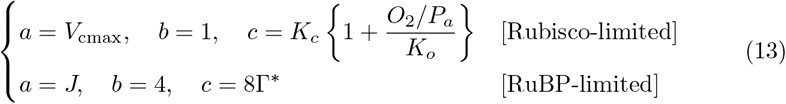

Here, we introduced three forms of *A*_*n*_(*C*_*i*_): *A*_*nR*_(*C*_*i*_), *A*_*nR*0_, and *A*_*nP*_. The subscripts “*R*” and “*P* “in Eq. (12) refer to the two limiting cases: Rubisco/RuBP-limited and Product-limited photosynthesis. “*R*0” in Eq. (12b) represents the special case where *a* = 0 under Rubisco/RuBP-limited condition. This case occurs, for example, at night when *J* = 0 due to the absence of light. Based on Eq (13) and possible range of variables shown in Table 1, we assume the following inequalities:

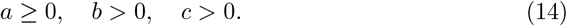

The function *A*_*nR*_(*C*_*i*_) depends on *C*_*i*_ and can take both positive and negative values. In contrast, *A*_*nR*0_ and *A*_*nP*_ are independent of *C*_*i*_. Specifically, *A*_*nR*0_ is always negative (Eq. (12b)), while *A*_*nP*_ is positive under a reasonable range of leaf temperatures (Eq. (12c)). Thus, we assume the following inequalities:

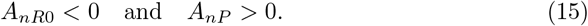

### 3.3 Two Functions for Stomatal Conductance: *F* (*C*_*i*_) and *G*(*C*_*i*_)

The *A*_*n*_-*E*-*g*_*s*_ model can be reduced to two independent equations for *g*_*s*_, both expressed as functions of *C*_*i*_. The first equation is derived from the CO_2_ flux equations (Eqs. (5) to (8)) and is denoted by *F* (*C*_*i*_). The second equation comes from the conductance model (Eq. (3)) and is represented by *G*(*C*_*i*_).

(a) *F* (*C*_*i*_)**: Stomatal conductance from CO**_2_ **flux** By eliminating *C*_*s*_ from the CO_2_ flux equations (Eqs. (5) to (8)), we can express *g*_*s*_ as:

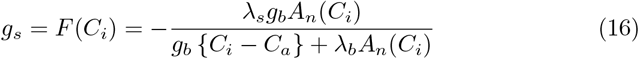 Note that *F* (*C*_*i*_) has three different forms, corresponding to the three different equations for *A*_*n*_(*C*_*i*_) (Eq. (12)), as follows:

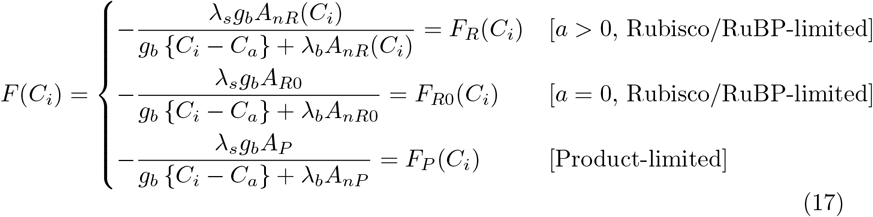 These forms are distinguished by the subscripts “*R*”, “*R*0”, and “*P* “, following the notation for *A*_*n*_(*C*_*i*_), resulting in *F*_*R*_(*C*_*i*_), *F*_*R*0_(*C*_*i*_), and *F*_*P*_ (*C*_*i*_).
(b) *G*(*C*_*i*_)**: Stomatal conductance from the conductance model** By eliminating the unknown variables *E*_*s*_, *E*_*b*_, *e*_*s*_, and *g*_*s*_ from Eqs. (4), (9), (10), (11), and (16), the vapor pressure deficit *D* can be written as:

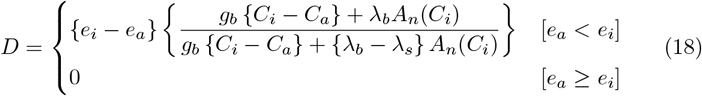 Substituting Eqs. (6), (8), and (18) into Eq. (3) to eliminate *D* and *C*_*s*_, we obtain:

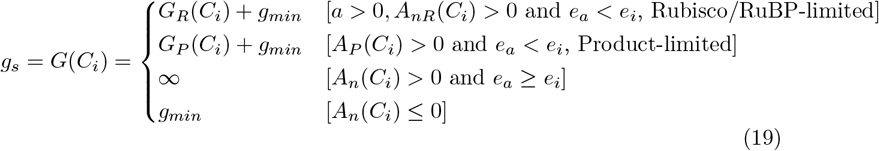 Here, *G*_*R*_(*C*_*i*_) and *G*_*P*_ (*C*_*i*_) are expressed as:

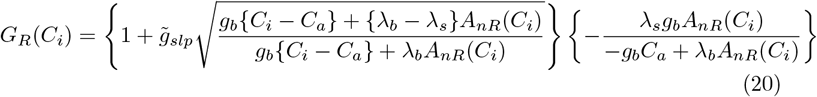

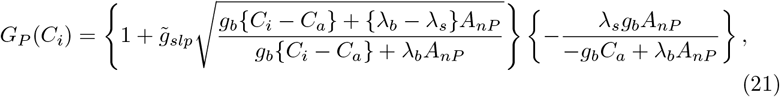

where 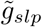 is given as:

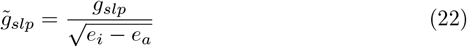

### 3.4 Steps for Proof

As shown in the previous section, the *A*_*n*_-*E*-*g*_*s*_ model (Section 3.1), which consists of 11 equations with 11 independent variables, can be reduced to two independent key functions: *F* (*C*_*i*_) and *G*(*C*_*i*_) (Eqs. (16) and (19)). These functions are expressed as:

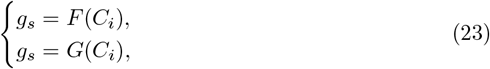

where both *F* (*C*_*i*_) and *G*(*C*_*i*_) express stomatal conductance, *g*_*s*_, as functions of the internal leaf CO_2_ concentration, *C*_*i*_. Thus, the value of *C*_*i*_ that satisfies all the equations in the *A*_*n*_-*E*-*g*_*s*_ model is the one for which *F* (*C*_*i*_) = *G*(*C*_*i*_) holds.

To facilitate further analysis, we reformulate the equation *F* (*C*_*i*_) = *G*(*C*_*i*_) as a combination of a function of *C*_*i*_ and a constant term:

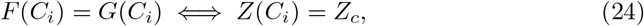

where *Z*(*C*_*i*_) represents a function of *C*_*i*_, and *Z*_*c*_ is a constant. The solution to the *A*_*n*_-*E*-*g*_*s*_ model is thus defined as the value of *C*_*i*_ that satisfies *Z*(*C*_*i*_) = *Z*_*c*_.

Using the Criteria for appropriate solutions (Section 2) and Eq. (24), the appropriate solutions to the *A*_*n*_-*E*-*g*_*s*_ model are defined as follows:

#### Definition 1

(Appropriate Solutions to the *A*_*n*_-*E*-*g*_*s*_ Model (I)). *C*_*i*_ *such that Z*(*C*_*i*_) = *Z*_*c*_, *g*_*s*_ *>* 0, *and C*_*i*_ *>* 0.

Since both *F* (*C*_*i*_) and *G*(*C*_*i*_) represent *g*_*s*_, equivalent definitions can be stated as:

#### Definition 2

(Appropriate Solutions to the *A*_*n*_-*E*-*g*_*s*_ Model (II)). *C*_*i*_ *such that Z*(*C*_*i*_) = *Z*_*c*_, *F* (*C*_*i*_) *>* 0, *and C*_*i*_ *>* 0.

#### Definition 3

(Appropriate Solutions to the *A*_*n*_-*E*-*g*_*s*_ Model (III)). *C*_*i*_ *such that Z*(*C*_*i*_) = *Z*_*c*_, *G*(*C*_*i*_) *>* 0, *and C*_*i*_ *>* 0.

Definition 1 is the most fundamental since it directly uses the Criteria for appropriate solutions. However, it does not provide an explicit expression for *g*_*s*_, making it less practical for analytical purposes. Definitions 2 and 3 provide explicit forms of *g*_*s*_ through *F* (*C*_*i*_) and *G*(*C*_*i*_), respectively. Given the simpler expression of *F* (*C*_*i*_) (Eqs. (16) and (19)), we adopt Definition 2 for our analysis.

Using Definition 2, the proof of the Theorem (Section 2) proceeds through the following steps:

1. **Step 1:** Identify the possible range of appropriate solutions by selecting the interval of *C*_*i*_ where *F* (*C*_*i*_) *>* 0 and *C*_*i*_ *>* 0 (Definition 2).
2. **Step 2:** Determine the distinct geometric combinations of *F* (*C*_*i*_) and *G*(*C*_*i*_) to identify the corresponding distinct geometric combinations of *Z*(*C*_*i*_) and *Z*_*c*_. Both *F* (*C*_*i*_) and *G*(*C*_*i*_) exhibit different geometries depending on external variables. Consequently, *Z*(*C*_*i*_) and *Z*_*c*_ also take on distinct geometries. By identifying the distinct geometric combinations of *F* (*C*_*i*_) and *G*(*C*_*i*_), the corresponding distinct geometric combinations of *Z*(*C*_*i*_) and *Z*_*c*_ can be determined.
3. **Step 3:** Analyze the geometry of *Z*(*C*_*i*_) for each combination identified in Step 2, within the possible range of appropriate solutions determined in Step 1.
4. **Step 4:** Determine the number of solutions for *C*_*i*_ that satisfy *Z*(*C*_*i*_) = *Z*_*c*_, *F* (*C*_*i*_) *>* 0, and *C*_*i*_ *>* 0, based on the geometries of *Z*(*C*_*i*_) analyzed in Step 3. This involves finding the intersections between *Z*(*C*_*i*_) and the line *Z*_*c*_ within the possible range of appropriate solutions. If it can be demonstrated that such an intersection always exists and is unique, the Theorem is proven.

## 4 Results

### 4.1 The Possible Range of Appropriate Solutions [Step 1]

To determine the possible range of appropriate solutions, defined as the interval of *C*_*i*_ where *F* (*C*_*i*_) *>* 0 and *C*_*i*_ *>* 0 (Definition 2), we analyze the geometries of *F* (*C*_*i*_) (Eq. (16)) and its primary component, *A*_*n*_(*C*_*i*_) (Eq. (12)).

The function *A*_*n*_(*C*_*i*_) takes three distinct forms: *A*_*nR*_(*C*_*i*_), *A*_*nR*0_, and *A*_*nP*_ (Eq. (12)). Among these, *A*_*nR*0_ and *A*_*nP*_ are trivial functions, such as negative and positive constants (Eq. (15)), and therefore do not require further analysis. As a result, this section focuses on *A*_*nR*_(*C*_*i*_).

Our analysis shows that *A*_*nR*_(*C*_*i*_) can exhibit five distinct geometries, denoted *A*_*nR*1_(*C*_*i*_) through *A*_*nR*5_(*C*_*i*_), depending on the limiting factors and external variables (see Section SI 1 for detailed analysis). Including the constant forms *A*_*nR*0_ and *A*_*P*_, this leads to seven distinct geometries for *A*_*n*_(*C*_*i*_). These geometries are illustrated in Figures 1a to 1g, along with key elements such as the air CO_2_ concentration (*C*_*a*_), the zero point 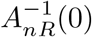,and the singularity point 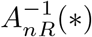.Specifically, 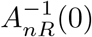 represents the zero of *A*_*nR*1_(*C*_*i*_) to *A*_*nR*4_(*C*_*i*_) (Eq. (SI 16)), while 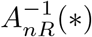 indicates the singularity of *A*_*nR*1_(*C*_*i*_) to *A*_*nR*5_(*C*_*i*_) (Eq. (SI 3)).

**Fig. 1:**
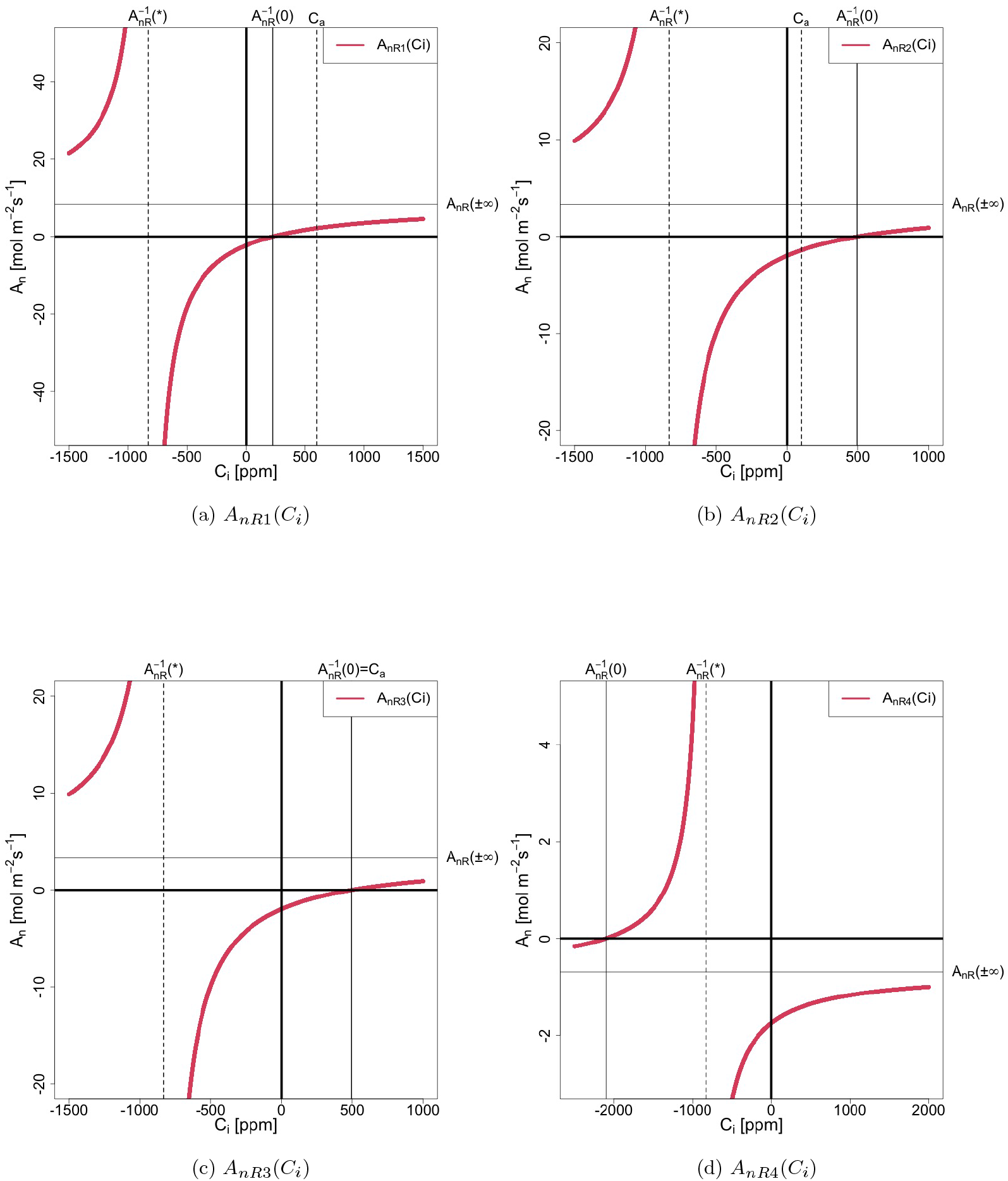
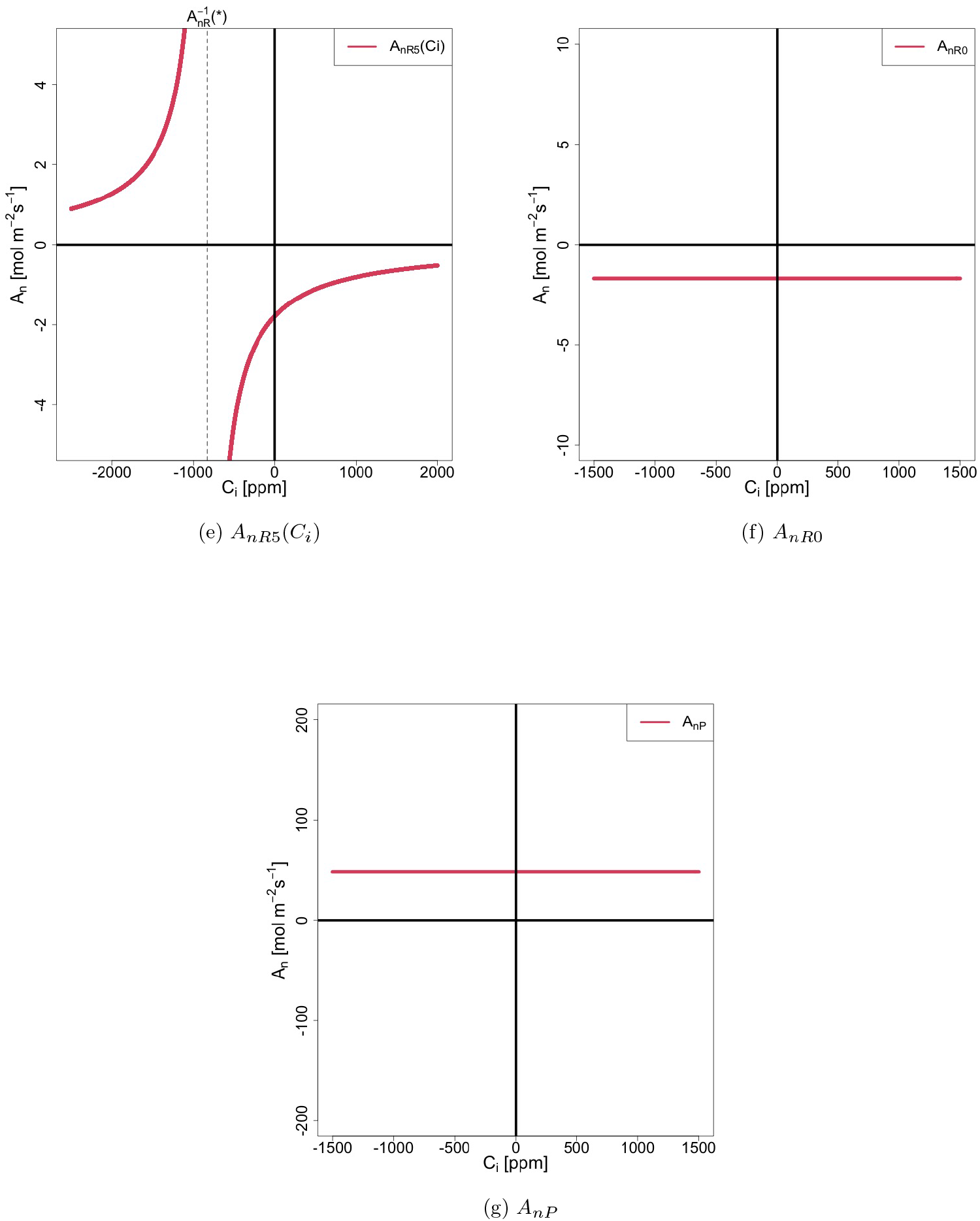
Geometry of *A*_*n*_(*C*_*i*_)

It is important to note that while the geometries of *A*_*nR*1_(*C*_*i*_) through *A*_*nR*5_(*C*_*i*_) differ depending on external variables, their mathematical expressions are identical to *A*_*nR*_(*C*_*i*_) (Eq. (12a)). Additionally, although *A*_*nR*1_(*C*_*i*_), *A*_*nR*2_(*C*_*i*_), and *A*_*nR*3_(*C*_*i*_) may appear similar (Figures 1a to 1c), they are distinguished by the relationship between *C*_*a*_ and 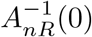.

Table 2 summarizes the conditions defining these seven distinct geometries of *A*_*n*_(*C*_*i*_). These conditions include factors such as limiting conditions (Rubisco/RuBP-limited or Product-limited) and the signs of *a, A*_*nR*_(*±*∞), and 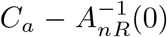.Here, *a* represents the maximum rate of carboxylation under Rubisco/RuBP-limited conditions or the electron transport rate under Product-limited conditions (Eq. (13)); *A*_*nR*_(*±*∞) denotes the photosynthesis rate as *C*_*i*_ → *±*∞ (Eq. (SI 7)). To classify the conditions for *A*_*nR*1_(*C*_*i*_) through *A*_*nR*5_(*C*_*i*_), we use [C1] to [C5] as labels (Table SI 1), which are also included in Table 2.

**Table 2:** Conditions and possible range of appropriate solutions for each geometry of *A*_*n*_(*C*_*i*_) and *F* (*C*_*i*_)

| $A_n(C_i)$ | $F(C_i)$ | Limiting | $a$ | $A_{nR}(\pm\infty)$ | $C_a - A_{nR}^{-1}(0)$ | [C] | Possible range of appropriate solutions | $A_n(C_i)$ |
| --- | --- | --- | --- | --- | --- | --- | --- | --- |
| $A_{nR1}(C_i)$ | $F_{R1}(C_i)$ | Rubisco/RuBP | $a > 0$ | $A_{nR}(\pm\infty) > 0$ | $C_a - A_{nR}^{-1}(0) > 0$ | [C1] | $F_R^{-1}(0) < C_i < F_R^{-1}(*)^+$ | $A_n(C_i) > 0$ |
| $A_{nR2}(C_i)$ | $F_{R2}(C_i)$ | Rubisco/RuBP | $a > 0$ | $A_{nR}(\pm\infty) > 0$ | $C_a - A_{nR}^{-1}(0) < 0$ | [C2] | $F_R^{-1}(*)^+ < C_i < F_R^{-1}(0)$ | $A_n(C_i) < 0$ |
| $A_{nR3}(C_i)$ | $F_{R3}(C_i)$ | Rubisco/RuBP | $a > 0$ | $A_{nR}(\pm\infty) > 0$ | $C_a - A_{nR}^{-1}(0) = 0$ | [C3] | $C_i = F_R^{-1}(*)^+ = F_R^{-1}(0)$ | $A_n(C_i) = 0$ |
| $A_{nR4}(C_i)$ | $F_{R4}(C_i)$ | Rubisco/RuBP | $a > 0$ | $A_{nR}(\pm\infty) < 0$ | | [C4] | $F_R^{-1}(*)^+ < C_i$ | $A_n(C_i) < 0$ |
| $A_{nR5}(C_i)$ | $F_{R5}(C_i)$ | Rubisco/RuBP | $a > 0$ | $A_{nR}(\pm\infty) = 0$ | | [C5] | $F_R^{-1}(*)^+ < C_i$ | $A_n(C_i) < 0$ |
| $A_{nR0}(C_i)$ | $F_{R0}(C_i)$ | Rubisco/RuBP | $a = 0$ | | | | $F_{R0}^{-1}(*) < C_i$ | $A_n(C_i) < 0$ |
| $A_{nP}(C_i)$ | $F_P(C_i)$ | Product | | | | | $0 < C_i < F_P^{-1}(*)$ | $A_n(C_i) > 0$ |

According to the seven distinct geometries of *A*_*n*_(*C*_*i*_), the function *F* (*C*_*i*_) also exhibits seven distinct geometries (Eq. (16)). These geometries are denoted with the same subscripts as those for *A*_*n*_(*C*_*i*_): *F*_*R*1_(*C*_*i*_) through *F*_*R*5_(*C*_*i*_), *F*_*R*0_(*C*_*i*_), and *F*_*P*_ (*C*_*i*_) (Table 2). Figures 2a to 2g illustrate these geometries, explicitly indicating the zeros and singularities of *F* (*C*_*i*_). Specifically, 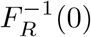 represents the zero of *F*_*R*1_(*C*_*i*_) through *F*_*R*4_(*C*_*i*_) (Eq. (SI 89)), while 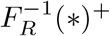 and 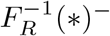 represent the singularities of these same functions (Eq. (SI 82)). Additionally, 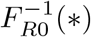 and 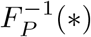 denote the singularities of *F*_*R*0_(*C*_*i*_) and *F*_*P*_ (*C*_*i*_), respectively (Eqs. (SI 117) and (SI 133)). Note that *F*_*R*3_(*C*_*i*_) takes the form 0*/*0 at 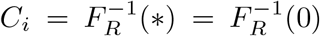,rendering it indeterminate at this point. As with *A*_*nR*_(*C*_*i*_), the geometries of *F*_*R*1_(*C*_*i*_) through *F*_*R*5_(*C*_*i*_) differ depending on external variables, although their mathematical expressions are identical to *F*_*R*_(*C*_*i*_) (Eq. (SI 20)).

**Fig. 2:**
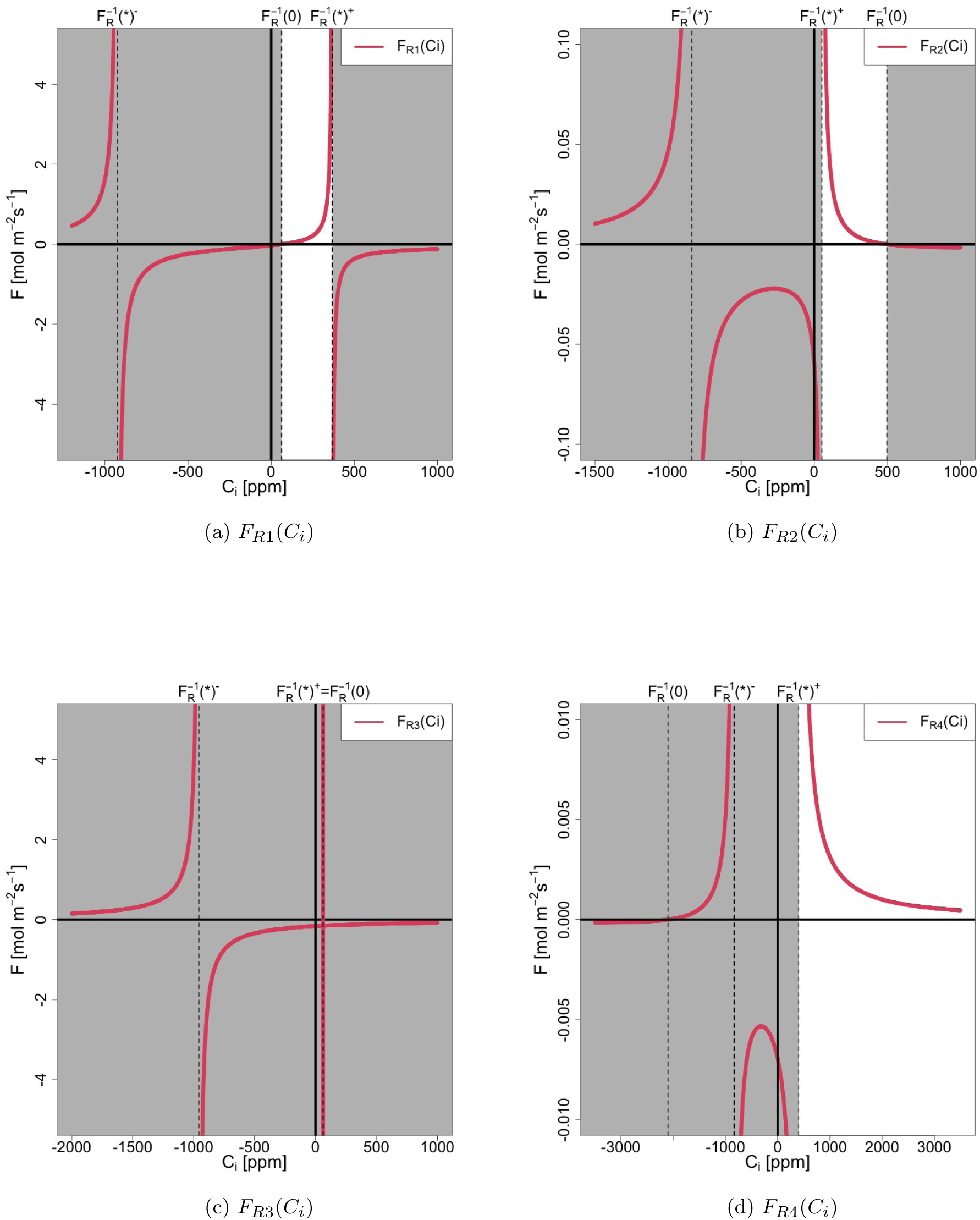
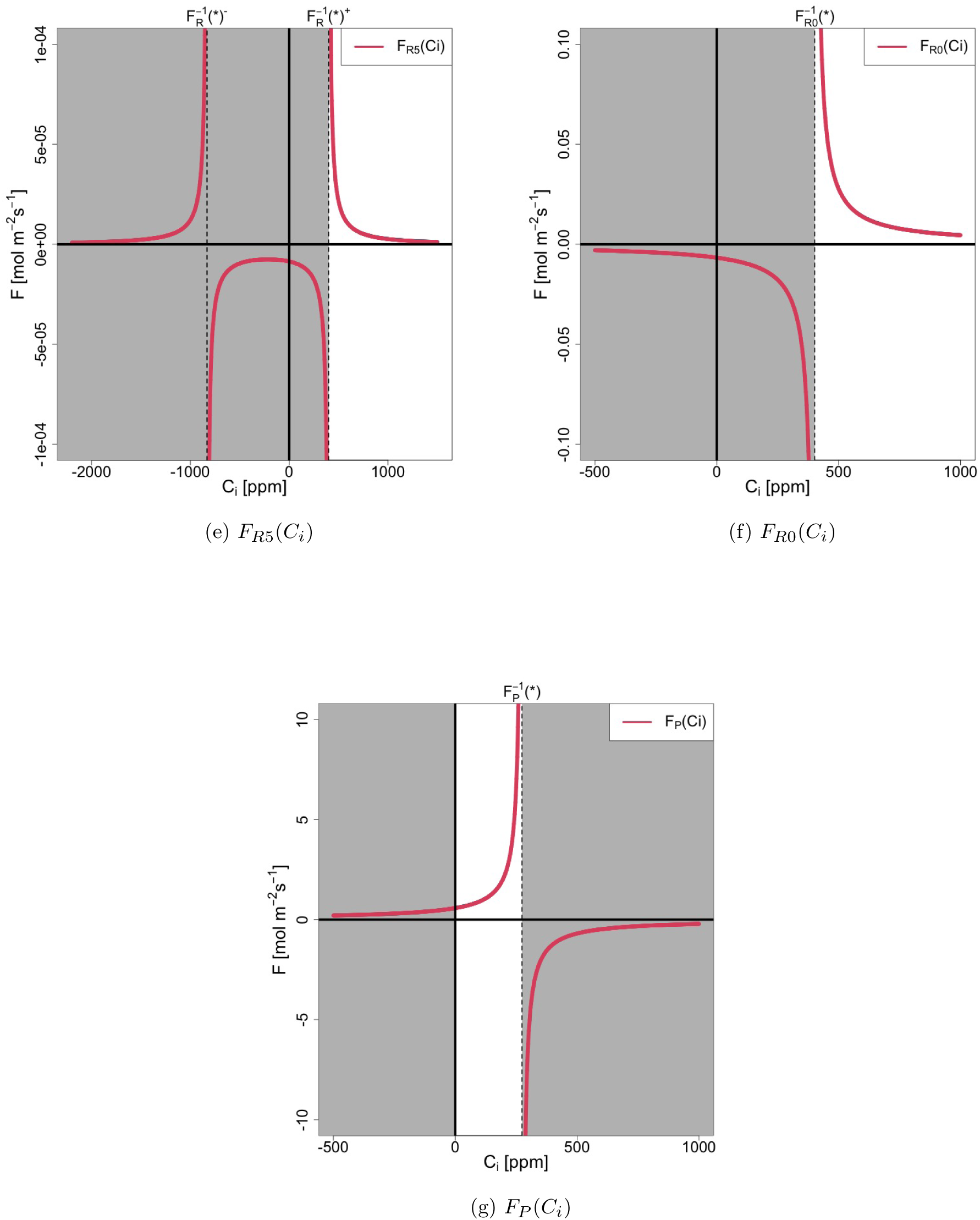
Geometry of *F* (*C*_*i*_)

From Figures 2a to 2g, the possible range of appropriate solutions can be identified by selecting the interval where *F* (*C*_*i*_) *>* 0 and *C*_*i*_ *>* 0. Table 2 summarizes the possible ranges of appropriate solutions for each geometry of *F* (*C*_*i*_). In these figures, regions outside the range are shaded in grey. It is important to note that the possible range of appropriate solutions for *F*_*R*3_(*C*_*i*_) consists solely of the indeterminate point 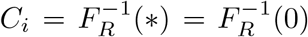.Therefore, special treatment is required for the analysis of *F*_*R*3_(*C*_*i*_). Additionally, *F* (*C*_*i*_) does not have any zeros within the possible range of appropriate solutions, as only the intervals where *F* (*C*_*i*_) *>* 0 are selected.

Once the possible range of appropriate solutions is determined, the sign of *A*_*n*_(*C*_*i*_) within that range can be analyzed using the geometries of *A*_*n*_(*C*_*i*_) (Figures 1a to 1g). The sign of *A*_*n*_(*C*_*i*_) is critical because it determines the specific form of *G*(*C*_*i*_) in the next step (Eq. (19)). Table 2 also summarizes the sign of *A*_*n*_(*C*_*i*_) for each case.

### 4.2 Combination of *Z*(*C*_*i*_) and *Z*_*c*_ [Step 2]

We explore the distinct geometric combinations of *F* (*C*_*i*_) and *G*(*C*_*i*_) to identify the corresponding distinct geometric combinations of *Z*(*C*_*i*_) and *Z*_*c*_.

According to Table 2, only *F*_*R*1_(*C*_*i*_) and *F*_*P*_ (*C*_*i*_) yield *A*_*n*_(*C*_*i*_) *>* 0 under Rubisco/RuBP-limited and Product-limited conditions, respectively. Based on these results and Eq. (19), when *A*_*n*_(*C*_*i*_) *>* 0, there are four possible combinations of *F* (*C*_*i*_) and *G*(*C*_*i*_): under the Rubisco/RuBP-limited condition, *F*_*R*1_(*C*_*i*_) can combine with either *G*_*R*_(*C*_*i*_) + *g*_*min*_ (when *e*_*a*_ *< e*_*i*_) or ∞ (when *e*_*a*_ ≥ *e*_*i*_); under the Product-limited condition, *F*_*P*_ (*C*_*i*_) can combine with either *G*_*P*_ (*C*_*i*_) + *g*_*min*_ (when *e*_*a*_ *< e*_*i*_) or ∞ (when *e*_*a*_ ≥ *e*_*i*_).

For the other five geometries of *F* (*C*_*i*_), namely *F*_*R*2_(*C*_*i*_) through *F*_*R*5_(*C*_*i*_) and *F*_*R*0_(*C*_*i*_), we have *A*_*n*_(*C*_*i*_) ≤ 0, as indicated in Table 2. Since *G*(*C*_*i*_) = *g*_*min*_ when *A*_*n*_(*C*_*i*_) ≤0 (Eq. (19)), these five forms of *F* (*C*_*i*_) combine only with *g*_*min*_.

Consequently, there are nine possible distinct geometric combinations between *F* (*C*_*i*_) and *G*(*C*_*i*_), summarized in Table 3. This table also shows the corresponding inequalities between *e*_*a*_ and *e*_*i*_, and the cases [C1] to [C5] for the conditions on *A*_*nR*1_(*C*_*i*_) to *A*_*nR*5_(*C*_*i*_) (Table 2).

**Table 3:**
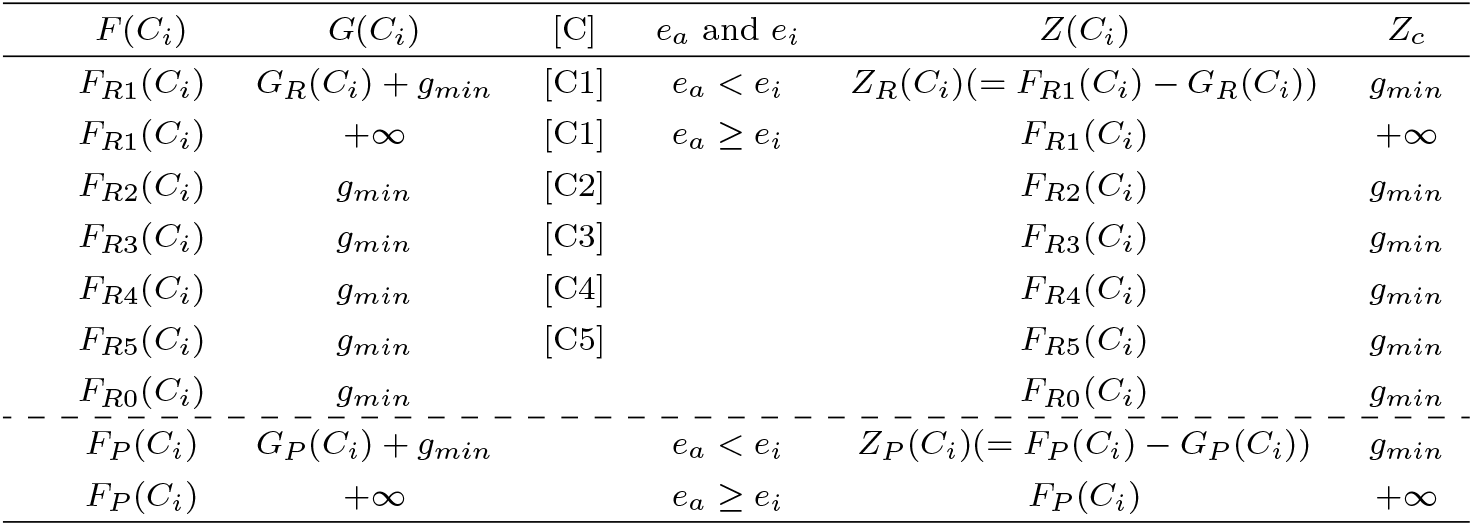
Distinct geometric combinations between *Z*(*C*_*i*_) and *Z*_*c*_.

| $F(C_i)$ | $G(C_i)$ | [C] | $e_a$ and $e_i$ | $Z(C_i)$ | $Z_c$ |
| --- | --- | --- | --- | --- | --- |
| $F_{R1}(C_i)$ | $G_R(C_i) + g_{min}$ | [C1] | $e_a < e_i$ | $Z_R(C_i)(= F_{R1}(C_i) - G_R(C_i))$ | $g_{min}$ |
| $F_{R1}(C_i)$ | $+\infty$ | [C1] | $e_a \geq e_i$ | $F_{R1}(C_i)$ | $+\infty$ |
| $F_{R2}(C_i)$ | $g_{min}$ | [C2] | | $F_{R2}(C_i)$ | $g_{min}$ |
| $F_{R3}(C_i)$ | $g_{min}$ | [C3] | | $F_{R3}(C_i)$ | $g_{min}$ |
| $F_{R4}(C_i)$ | $g_{min}$ | [C4] | | $F_{R4}(C_i)$ | $g_{min}$ |
| $F_{R5}(C_i)$ | $g_{min}$ | [C5] | | $F_{R5}(C_i)$ | $g_{min}$ |
| $F_{R0}(C_i)$ | $g_{min}$ | | | $F_{R0}(C_i)$ | $g_{min}$ |
| $F_P(C_i)$ | $G_P(C_i) + g_{min}$ | | $e_a < e_i$ | $Z_P(C_i)(= F_P(C_i) - G_P(C_i))$ | $g_{min}$ |
| $F_P(C_i)$ | $+\infty$ | | $e_a \geq e_i$ | $F_P(C_i)$ | $+\infty$ |

Once the combinations of *F* (*C*_*i*_) and *G*(*C*_*i*_) are identified, the corresponding distinct geometric combinations of *Z*(*C*_*i*_) and *Z*_*c*_ can be derived using Eq. (24). Table 3 summarizes these combinations of *Z*(*C*_*i*_) and *Z*_*c*_. Two new functions, *Z*_*R*_(*C*_*i*_) and *Z*_*P*_ (*C*_*i*_), are introduced here and are also included in Table 3.

### 4.3 Geometry of *Z*(*C*_*i*_) [Step 3]

We examine the geometries of the functions *Z*(*C*_*i*_) identified in Step 2, focusing on the possible range of appropriate solutions identified in Step 1. Among the distinct forms of *Z*(*C*_*i*_) listed in Table 3, the geometries of the functions *F*_*R*1_(*C*_*i*_) through *F*_*R*5_(*C*_*i*_), *F*_*R*0_(*C*_*i*_), and *F*_*P*_ (*C*_*i*_) have already been identified in Section 4.1. Therefore, we will focus on *Z*_*R*_(*C*_*i*_) and *Z*_*P*_ (*C*_*i*_), introduced in Section 4.2.

We find that *Z*_*R*_(*C*_*i*_) exhibits two distinct geometries within the possible range of appropriate solutions, depending on external variables (Section SI 5.7). One geometry always has a zero within this range, while the other does not, depending on the signs of 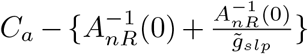 (Eq. (SI 410)). These geometries are denoted as *Z*_*RI*_ (*C*_i_) and *Z*_*RII*_ (*C*_*i*_), and these two cases are classified as [ZI] and [ZII]. Figures 3a and 3b display the curves of *Z*_*RI*_ (*C*_*i*_) and *Z*_*RII*_ (*C*_*i*_), respectively. The zero of *Z*_*RI*_ (*C*_*i*_) is denoted as 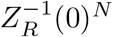 (Eq. (SI 437)) and is shown in Figure 3a.

**Fig. 3:**
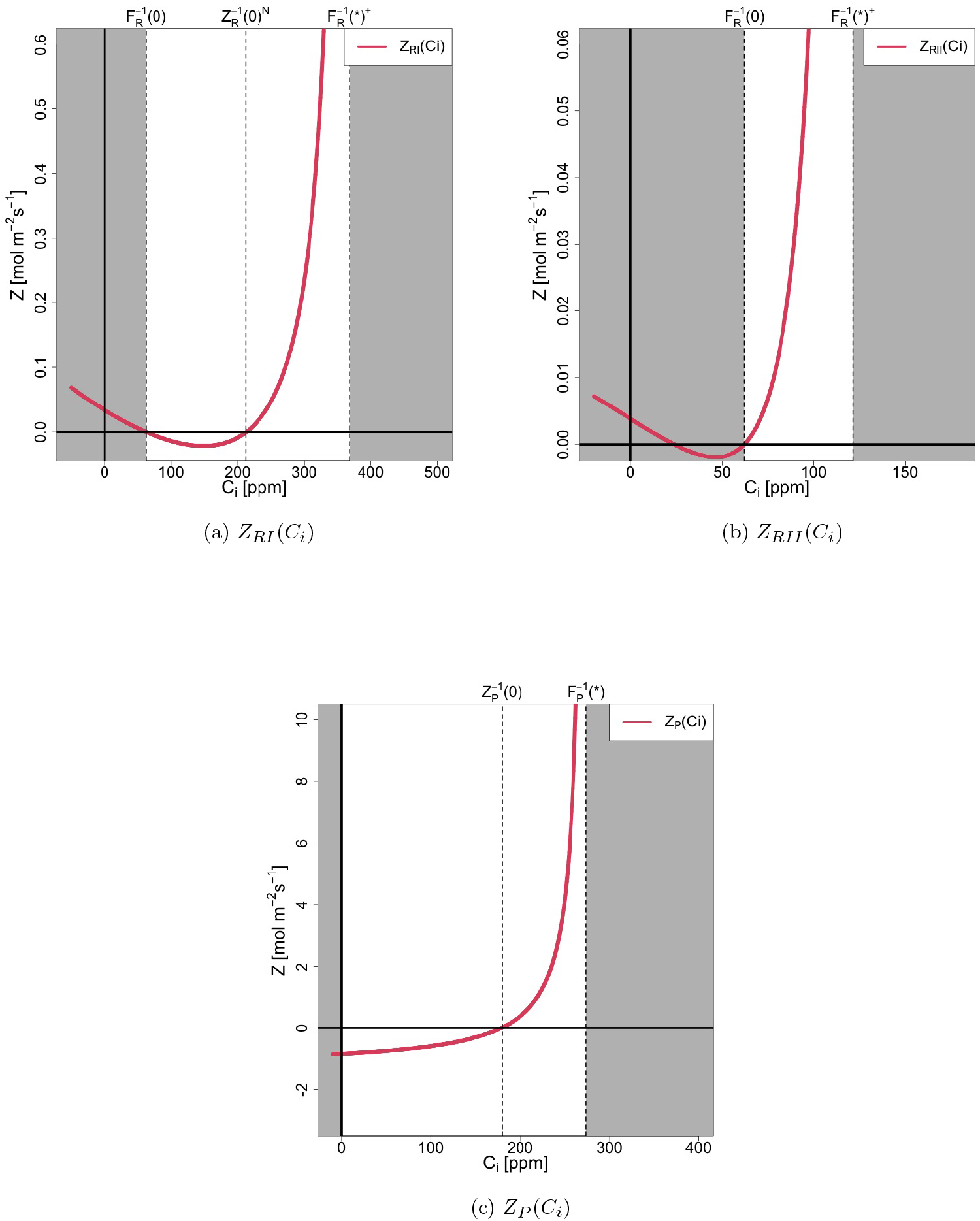
Geometry of *Z*_*RI*_ (*C*_*i*_), *Z*_*RII*_ (*C*_*i*_), and *Z*_*P*_ (*C*_*i*_)

For *Z*_*P*_ (*C*_*i*_), we find that it exhibits only one geometry, which always has a zero within the possible range of appropriate solutions. Figure 3c shows the curve of *Z*_*P*_ (*C*_*i*_). The zero of *Z*_*P*_ (*C*_*i*_) is denoted as 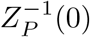 (Eq. (SI 508)) and is also shown in Figure 3c.

Finally, there are ten distinct geometric combinations of *Z*(*C*_*i*_) and *Z*_*c*_, denoted as [R-I] to [R-VIII] for Rubisco/RuBP-limited conditions and [P-I] to [P-II] for Product-limited conditions. Table 4 summarizes these combinations and their associated conditions. Note that there is a zero within the possible range of appropriate solutions only for cases [R-I] and [P-I], with no zeros in the other cases. In addition, since *g*_*min*_ *>* 0 (Table 1), *Z*_*c*_ *>* 0 for all cases (Table 3).

**Table 4:** Combinations of *F* (*C*_*i*_), *G*(*C*_*i*_), *Z*(*C*_*i*_), and *Z*_*c*_, and their conditions.

| | $F(C_i)$ | $G(C_i)$ | $Z(C_i)$ | $Z_c$ | Limited conditions | $a$ | $A_{nR}(\pm\infty)$ | $C_a - A_{nR}^{-1}(0)$ | $e_a$ and $e_i$ | $C_a - A_{nR}^{-1}(0) \left\{ 1 + \frac{1}{g_{slp}} \right\}$ | [C] | [Z] |
| --- | --- | --- | --- | --- | --- | --- | --- | --- | --- | --- | --- | --- |
| R-I | $F_{R1}(C_i)$ | $G_R(C_i) + g_{min}$ | $Z_{RI}(C_i)$ | $g_{min}$ | Rubisco/RuBP | $a > 0$ | $A_{nR}(\pm\infty) > 0$ | $C_a - A_{nR}^{-1}(0) > 0$ | $e_a < e_i$ | $C_a - A_{nR}^{-1}(0) \left\{ 1 + \frac{1}{g_{slp}} \right\} > 0$ | [C1] | [ZI] |
| R-II | $F_{R1}(C_i)$ | $G_R(C_i) + g_{min}$ | $Z_{RII}(C_i)$ | $g_{min}$ | Rubisco/RuBP | $a > 0$ | $A_{nR}(\pm\infty) > 0$ | $C_a - A_{nR}^{-1}(0) > 0$ | $e_a < e_i$ | $C_a - A_{nR}^{-1}(0) \left\{ 1 + \frac{1}{g_{slp}} \right\} \leq 0$ | [C1] | [ZII] |
| R-III | $F_{R1}(C_i)$ | $+\infty$ | $F_{R1}(C_i)$ | $+\infty$ | Rubisco/RuBP | $a > 0$ | $A_{nR}(\pm\infty) > 0$ | $C_a - A_{nR}^{-1}(0) > 0$ | $e_a \geq e_i$ | | [C1] | |
| R-IV | $F_{R2}(C_i)$ | $g_{min}$ | $F_{R2}(C_i)$ | $g_{min}$ | Rubisco/RuBP | $a > 0$ | $A_{nR}(\pm\infty) > 0$ | $C_a - A_{nR}^{-1}(0) < 0$ | | | [C2] | |
| R-V | $F_{R3}(C_i)$ | $g_{min}$ | $F_{R3}(C_i)$ | $g_{min}$ | Rubisco/RuBP | $a > 0$ | $A_{nR}(\pm\infty) > 0$ | $C_a - A_{nR}^{-1}(0) = 0$ | | | [C3] | |
| R-VI | $F_{R4}(C_i)$ | $g_{min}$ | $F_{R4}(C_i)$ | $g_{min}$ | Rubisco/RuBP | $a > 0$ | $A_{nR}(\pm\infty) < 0$ | | | | [C4] | |
| R-VII | $F_{R5}(C_i)$ | $g_{min}$ | $F_{R5}(C_i)$ | $g_{min}$ | Rubisco/RuBP | $a > 0$ | $A_{nR}(\pm\infty) = 0$ | | | | [C5] | |
| R-VIII | $F_{R0}(C_i)$ | $g_{min}$ | $F_{R0}(C_i)$ | $g_{min}$ | Rubisco/RuBP | $a = 0$ | | | | | | |
| P-I | $F_P(C_i)$ | $G_P(C_i) + g_{min}$ | $Z_P(C_i)$ | $g_{min}$ | Product | | | | $e_a < e_i$ | | | |
| P-II | $F_P(C_i)$ | $+\infty$ | $F_P(C_i)$ | $+\infty$ | Product | | | | $e_a \geq e_i$ | | | |

If *Z*(*C*_*i*_) has zeros within the possible range of appropriate solutions, the range can be refined by excluding the interval of *C*_*i*_ where *Z*(*C*_*i*_) *<* 0. This is because the solutions for *Z*(*C*_*i*_) = *Z*_*c*_ (Eq. 24) must exist within the range of *C*_*i*_ where *Z*(*C*_*i*_) *>* 0, given that *Z*_*c*_ *>* 0. Table 5 presents the modified range of possible solutions for cases [R-I] and [P-I].

**Table 5:** Possible Range of Appropriate Solutions.

|  | Range | Modified Range |
| --- | --- | --- |
| R-I | $F_R^{-1}(0) < C_i < F_R^{-1}(*)^+$ | $Z_R^{-1}(0)^N < C_i < F_R^{-1}(*)^+$ |
| R-II | $F_R^{-1}(0) < C_i < F_R^{-1}(*)^+$ | |
| R-III | $F_R^{-1}(0) < C_i < F_R^{-1}(*)^+$ | |
| R-IV | $F_R^{-1}(*)^+ < C_i < F_R^{-1}(0)$ | |
| R-V | $C_i = F_R^{-1}(*)^+ = F_R^{-1}(0)$ | |
| R-VI | $F_R^{-1}(*)^+ < C_i$ | |
| R-VII | $F_R^{-1}(*)^+ < C_i$ | |
| R-VIII | $F_R^{-1}(*) < C_i$ | |
| P-I | $0 < C_i < F_P^{-1}(*)$ | $Z_P^{-1}(0) < C_i < F_P^{-1}(*)$ |
| P-II | $0 < C_i < F_P^{-1}(*)$ | |

To identify the geometry of *Z*(*C*_*i*_) within the possible range of appropriate solutions, we calculated the values of *Z*(*C*_*i*_) at the endpoints of these ranges (Table 5) and the derivatives of *Z*(*C*_*i*_) within the ranges. The results of these calculations are summarized in Table 6. Note that for the cases [R-I] and [P-I], the modified possible range of appropriate solutions was used (Table 5). Additionally, for the case [R-V], these calculations could not be performed as the range of appropriate solutions consists only of the single indeterminate point 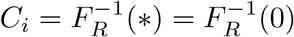.

**Table 6:**
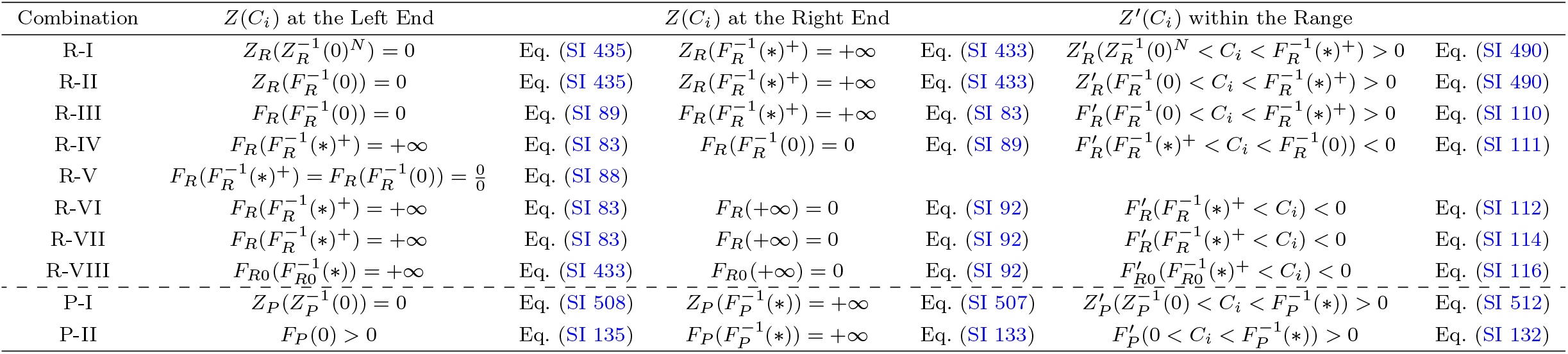
Geometry of *Z*(*C*_*i*_)

We find that the derivatives of *Z*(*C*_*i*_) are non-zero within the possible range of appropriate solutions, except for the case of [R-V], which indicates that *Z*(*C*_*i*_) is either monotonically increasing or decreasing within the range. Furthermore, for all cases except [R-V] and [P-II], one end of the possible range of appropriate solutions is a zero of *Z*(*C*_*i*_), while the other end is a singularity.

### 4.4 Number of Intersections between *Z*(*C*_*i*_) and *Z*_*c*_ [Step 4]

For cases [R-I, II, IV, VI to VIII, and P-I], *Z*(*C*_*i*_) = *Z*_*c*_ ⇔ *Z*(*C*_*i*_) = *g*_*min*_ (Table 4). At one end of the possible range of appropriate solutions, *Z*(*C*_*i*_) is zero or approaches zero, while at the other end, *Z*(*C*_*i*_) approaches +∞ (Table 6). Note that the modified possible range of appropriate solutions is applied for cases [R-I] and [P-I] (Table 5).

Since *g*_*min*_ *>* 0 (Table 1), *Z*(*C*_*i*_) satisfies *Z*(*C*_*i*_) *< g*_*min*_ at one end and *Z*(*C*_*i*_) *> g*_*min*_ at the other end. Furthermore, within this range, *Z*(*C*_*i*_) is either monotonically increasing or monotonically decreasing, depending on the inequality between *Z*(*C*_*i*_) and *g*_*min*_ at both ends (Table 6). Based on these geometric characteristics, cases [R-I, II, IV, VI to VIII, and P-I] can be divided into two groups. Table 7 summarizes these groups, as follows:

**Table 7:** *Z*(*C*_*i*_) and *Z*^*′*^(*C*_*i*_) for cases [R-I, II, IV, VI to VIII, and P-I].

| Group | Left end | Right end | $Z'(C_i)$ |
| --- | --- | --- | --- |
| R-I, II, and P-I | $Z(C_i) < g_{min}$ | $Z(C_i) > g_{min}$ | $Z'(C_i) > 0$ |
| R-IV, VI, VII, and VIII | $Z(C_i) > g_{min}$ | $Z(C_i) < g_{min}$ | $Z'(C_i) < 0$ |

In both groups, *Z*(*C*_*i*_) and *g*_*min*_ intersect exactly once within the range of appropriate solutions. Therefore, we have proven that there is always a unique appropriate solution for cases [R-I, II, IV, VI to VIII, and P-I]. Figures 4(a), (b), (d), and (f) through (i) illustrate the single intersection between *Z*(*C*_*i*_) and *Z*_*c*_ within the possible range of appropriate solutions for these cases.

**Fig. 4:**
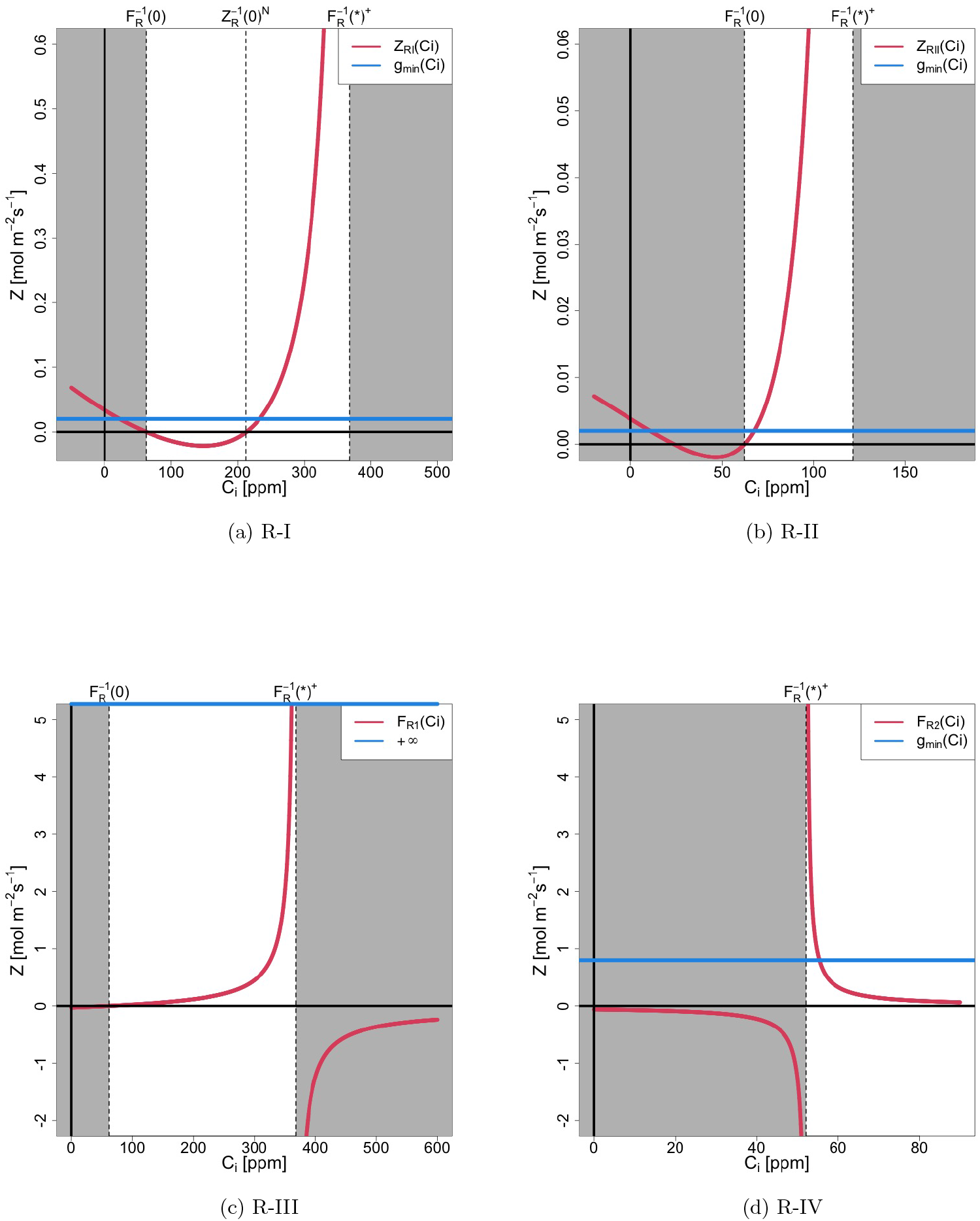
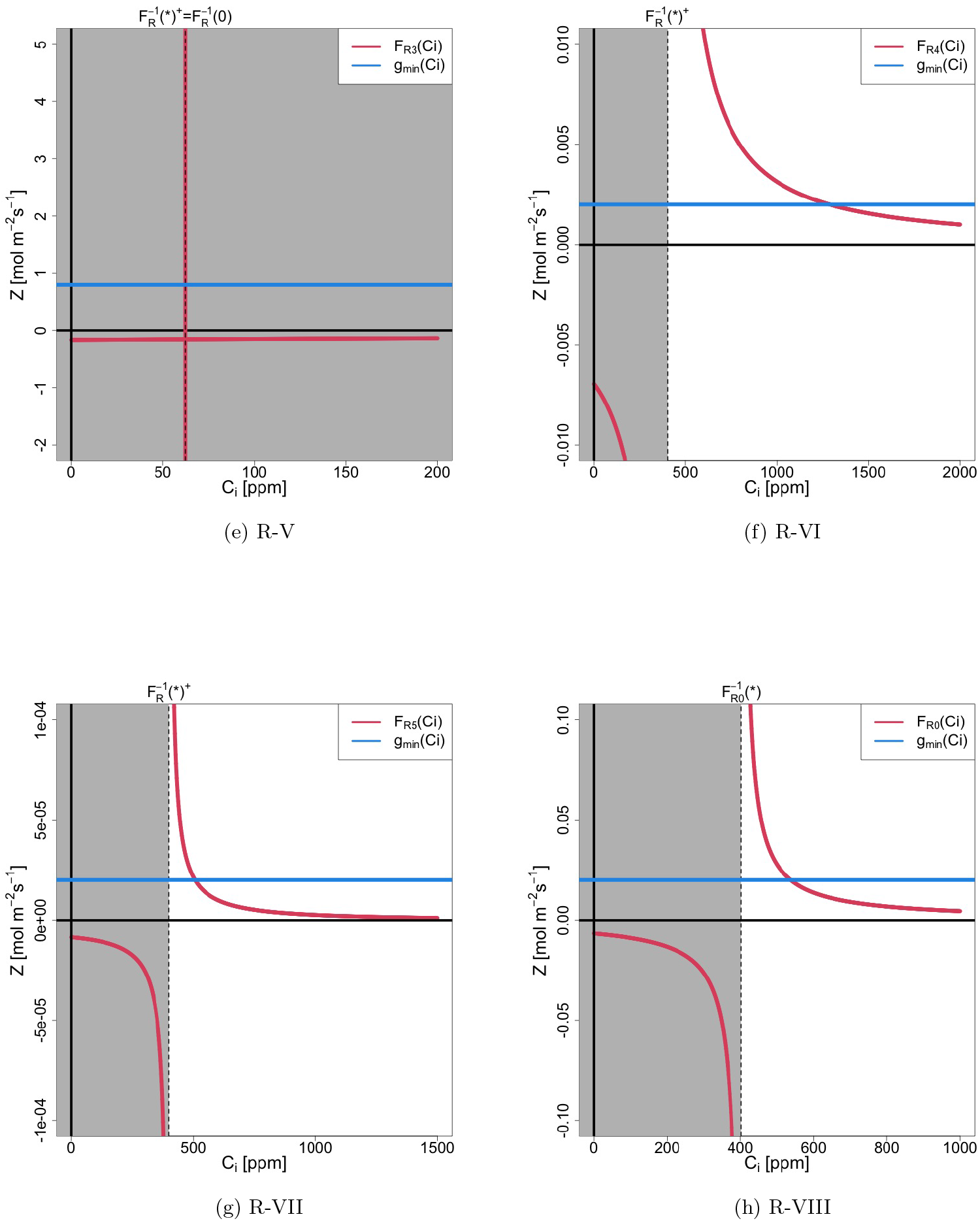
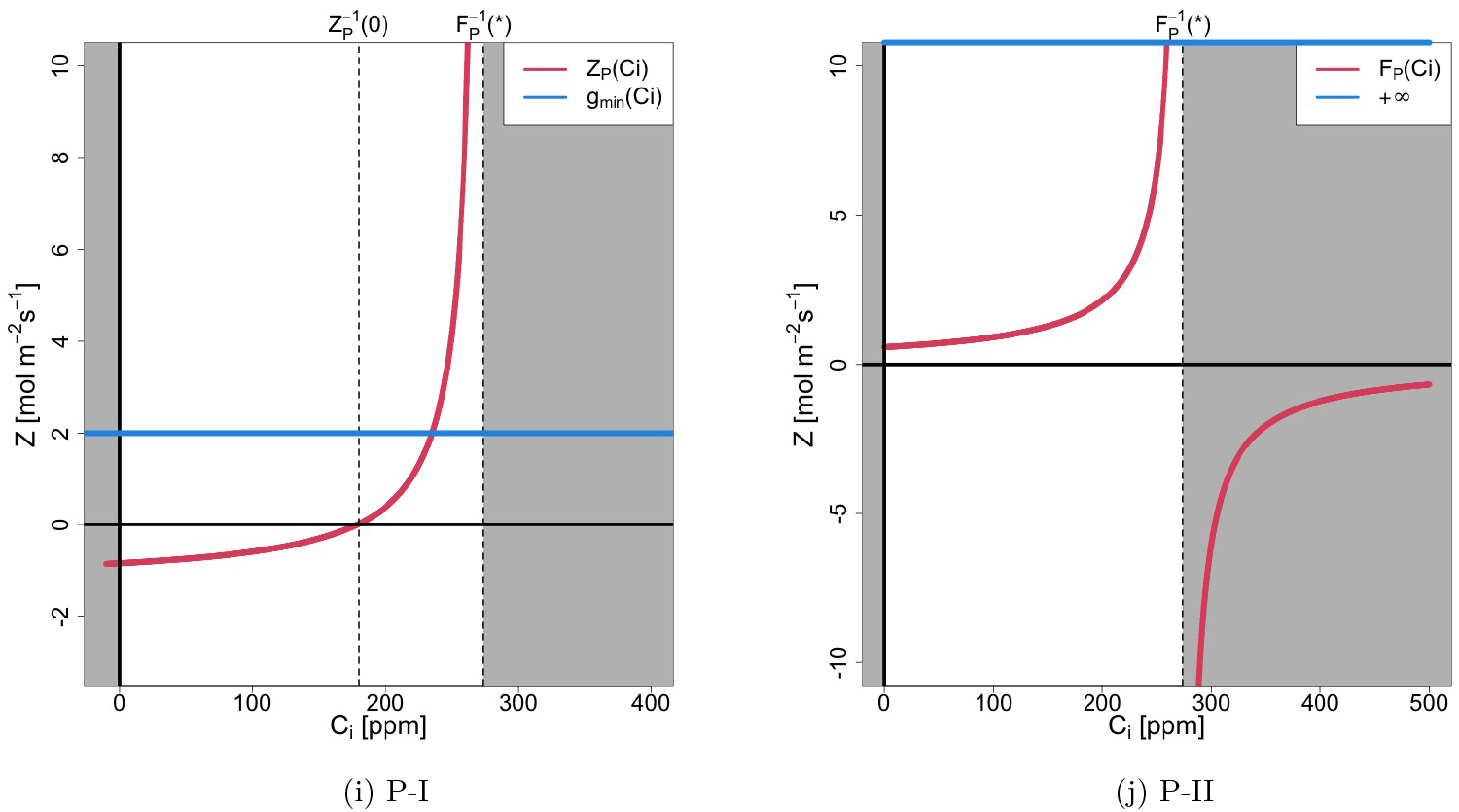
*Z*(*C*_*i*_) (red line) and *Z*_*c*_ (blue line)

For [R-V], *Z*(*C*_*i*_) = *Z*_*c*_ ⇔ *F*_*R*_(*C*_*i*_) = *g*_*min*_. According to Table 2, the only possible solution is 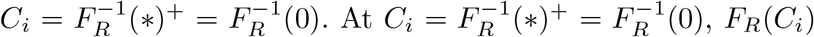 is indeterminate, meaning it can take any value at this point. Therefore, 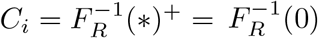 represents the unique solution for *F*_*R*_(*C*_*i*_) = *g*_*min*_. Thus, we have proven that there is always a unique solution for [R-V]. Figure 4(e) illustrates the single intersection between *Z*(*C*_*i*_) and *Z*_*c*_ at the only possible solution for this case.

For the remaining cases [R-III] and [P-II], *Z*(*C*_*i*_) = *Z*_*c*_ ⇔ *F*_*R*_(*C*_*i*_) = +∞ for [R-III] and *F*_*P*_ (*C*_*i*_) = +∞ for [P-II], as indicated in Table 4. The only *C*_*i*_ that satisfies *F*_*R*_(*C*_*i*_) = +∞ or *F*_*P*_ (*C*_*i*_) = +∞ within the range of *C*_*i*_ *>* 0 corresponds to their respective singularities: 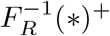 for [R-III] and 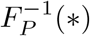 for [P-II]. Thus, 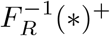 for [R-III] and 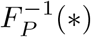 for [P-II] are the unique solutions for these cases. Therefore, we have proven that there is always a unique appropriate solution for the remaining cases [R-III] and [P-II]. Figures 4(c) and (j) illustrate the single intersection between *Z*(*C*_*i*_) and *Z*_*c*_.

It is important to note that *F*_*R*_(*C*_*i*_) and *F*_*P*_ (*C*_*i*_) are strictly mathematically undefined at their respective singularities. This issue arises from the divergence of Medlyn’s stomatal conductance equation at *D* = 0 (see Eq. (3)). To address this, stomatal resistance *r*_*s*_, expressed as the inverse of *g*_*s*_, can be used when *D* = 0. By using *r*_*s*_, we rigorously mathematically obtain the unique solution 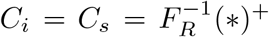 from Eqs. (5), (6) and (8) when *D* = 0. Thus the mathematically rigorous solution obtained using *r*_*s*_ is identical to the solution obtained through the less rigorous treatment that directly considers *g*_*s*_ at *D* = 0.

Thus, as demonstrated above, for all cases, there is always exactly one solution satisfying *Z*(*C*_*i*_) = *Z*_*c*_ within the possible range of appropriate solutions. Consequently, the Theorem is proven.

## 5 Discussion

### 5.1 Existence and uniqueness of solutions

In this study, we mathematically proved the existence and uniqueness of biologically and physically appropriate solutions for the *A*_*n*_-*E*-*g*_*s*_ model, a standard model for estimating photosynthesis, transpiration, and stomatal conductance. This theorem resolves the longstanding “solution selection” problem that has persisted for over 30 years since the model was first proposed by Collatz et al. (1991) ([11]). By identifying the only biologically and physically appropriate solution, the theorem ensures the unambiguous selection of the correct solution. Resolving the “solution selection” problem has far-reaching implications for diverse fields that rely on the *A*_*n*_-*E*-*g*_*s*_ model, including botany, plant ecology, agriculture, hydrology, and earth sciences. It strengthens the model’s reliability for estimating photosynthesis, transpiration, and stomatal conductance, which are critical for understanding global carbon and water cycles, assessing climate change impacts, and improving agricultural resource management.

Prior to resolving this problem, there was no guarantee that the solutions obtained by the model were correct, as there were multiple valid solutions. For example, most vegetation and ecosystem models used in the Global Carbon Project (GCP) to annually estimate the global carbon cycle rely on the *A*_*n*_-*E*-*g*_*s*_ model. This raised the possibility that such estimates, along with the conclusions derived from them, could be incorrect. However, the theorem presented in this study guarantees the correctness of past estimates, including those by the GCP, provided that they confirm *C*_*i*_ *>* 0 and *g*_*s*_ *>* 0. Therefore, we strongly recommend verifying whether *C*_*i*_ *>* 0 and *g*_*s*_ *>* 0 were satisfied in past estimates. In fact, since it is likely that many of these estimates already confirmed *C*_*i*_ *>* 0 and *g*_*s*_ *>* 0, the possibility that previous estimates were incorrect is low. Nonetheless, this does not diminish the significance of the mathematical proof presented here. Even if *C*_*i*_ *>* 0 and *g*_*s*_ *>* 0 were confirmed in past estimates, there was previously no guarantee that the resulting estimates were correct. Similarly, in future studies, by ensuring that estimates satisfy *C*_*i*_ *>* 0 and *g*_*s*_ *>* 0, it can be concluded that these estimates are correct. Accordingly, this study’s theorem will have significant benefits for research in related fields, spanning from past to future endeavors.

The requirement of two criteria (*C*_*i*_ *>* 0 and *g*_*s*_ *>* 0) for ensuring the existence and uniqueness of a solution is far from trivial. This is because it is possible that either condition alone might suffice, or that additional conditions could be necessary. Let us first consider the case where only *C*_*i*_ *>* 0 is assumed. In this case, as shown in Figure 4a, there are two intersections satisfying *Z*(*C*_*i*_) = *Z*_*c*_, which means the uniqueness of the solution cannot be guaranteed. By introducing the condition *g*_*s*_ *>* 0, the range of solutions is restricted, and it can be demonstrated that there is always a single solution within this range. Similarly, if we were to assume only *g*_*s*_ *>* 0, we would consider the region where *F* (*C*_*i*_) *>* 0. For example, as shown in Figure 2d, there exists a solution satisfying *Z*(*C*_*i*_) = *F*_*R*4_(*C*_*i*_) = *Z*_*c*_ in the region where *C*_*i*_ *<* 0. Therefore, it is only with the combination of these two criteria (*C*_*i*_ *>* 0 and *g*_*s*_ *>* 0) that the existence and uniqueness of the solution can be proven. Furthermore, this proof also demonstrates that no additional conditions are necessary beyond these two.

Mathematical theorems are relatively rare in biology and botany, making this study a valuable contribution to the field’s body of knowledge. Furthermore, this research stands out as one of the few that elucidates the mathematical foundations in the long history of photosynthesis research, marking it as a significant milestone. This mathematical framework not only enhances confidence in current modeling approaches but also paves the way for new research directions and practical applications in disciplines that depend on precise estimates of photosynthesis, transpiration, and stomatal conductance.

### 5.2 Challenges

The *A*_*n*_-*E*-*g*_*s*_ model has several variants, and proving the existence and uniqueness of solutions for these variants is not straightforward. In particular, numerous models have been proposed for stomatal conductance ([27]). This study specifically provides the proof for the Medlyn model, one of the major stomatal conductance models. However, if the existence and uniqueness of solutions cannot be demonstrated for a given stomatal conductance model, it may indicate that the model is either flawed or requires additional conditions. From a broader perspective, the next challenge is to identify the general form of stomatal conductance models that guarantee the existence and uniqueness of solutions. Once identified, such a general form could impose constraints on stomatal conductance models and clarify whether modifications to equations or additional conditions are necessary for the many existing models.

This study focused on C3 plants, but similar research could be conducted for C4 plants. For C4 plants, using relatively simple models ([15, 32]) might not be overly challenging for this proof. However, if more complex models like those proposed by Yin and Struik (2009)([33]) are used, the task might be somewhat more difficult. Nevertheless, the approach for C3 plants has been established, so it is likely only a matter of time before this is also proven for C4 plants. If this cannot be proven for C4 plants, conversely it would be a very intriguing finding.

As pointed out in Masutomi (2023)([24]), it has been recently understood that nocturnal stomatal conductance is not constantly minimum assumed in this study (Eq. (3)) and seems to be correlated with respiration rate ([34–36]). However, the relationship with environmental factors such as humidity and CO_2_ during nighttime is not well understood, and the final model equation incorporating these environmental variables remains unknown. Once this becomes clear, proving the existence and uniqueness of solutions for the *A*_*n*_-*E*-*g*_*s*_ model incorporating such nocturnal stomatal conductance models will become an important future challenge.

As described in the supplementary information, the proof presented in this study is highly lengthy and complex. While we expect that a simpler proof exists, we were unable to find one. This challenge remains unresolved and will be passed on to the next generation of researchers to tackle.

In this study, we focused on the *A*_*n*_-*E*-*g*_*s*_ model, which deals with only steady-state photosynthesis, transpiration, and stomatal conductance. However, more general models that describe non-steady states can also be considered. These models are expected to be formulated as dynamic systems of differential equations involving time, offering a much richer and more intricate landscape. Nevertheless, in the real world, a single value is ultimately selected, and the concept of existence and uniqueness of solutions demonstrated in this study undoubtedly applies to such dynamic systems of differential equations as well.

## Supplementary information

The details of the calculation for the proof are given as supplementary information.

## Acknowledgements

This study was partly supported by the Environment Research and Technology Development Fund (JPMEERF21S12007) of the Environmental Restoration and Conservation Agency (ERCA), and JSPS KAKENHI Grant Number JP23H00351. We would like to express our gratitude to Professor Yoshimura and Professor Nitta of The University of Tokyo. This research would not have been completed without the fruitful discussions with researchers at the “Land Model Research Meeting” they have organized. Additionally, we acknowledge the use of ChatGPT, developed by OpenAI, for its assistance in translating this manuscript from Japanese to English, as well as for proofreading and refining the English text, which greatly improved its clarity and readability.

## Declarations

- Competing interests The authors declare no competing interest.
- Author contribution Y.M designed and carried out the study, and wrote the manuscript. K.K discussed and supervised all aspects of the study.

## Appendix A Mathematical notations and conventions

In this study, parenthesis “()” is used for denoting the argument of functions. For example, a function “*F* “of *C*_*i*_ is expressed as *F* (*C*_*i*_). When an inequality is used as the argument of a function, it represents a condition. For example, *F* (*a < C*_*i*_ *< b*) *<* 0 means that *F* (*C*_*i*_) *<* 0 for *a < C*_*i*_ *< b*. Curly brackets “{}” is used for separating a group of equations, such as *a {b* + *c}* indicating *ab* + *ac*. Square brackets “[]” is used for expressing conditions under which the respective equation is valid.

Suppose that ℱ (*C*_*i*_) is a function of *C*_*i*_. The following notations are used in the present study

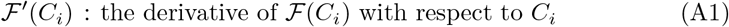

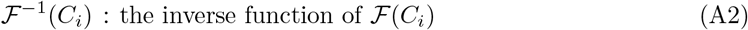

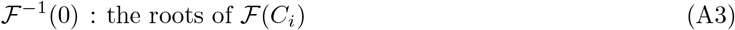

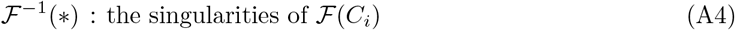

Note that we also treat *C*_*i*_ as a root if *ℱ*(*C*_*i*_) results in indeterminate like *ℱ*(*C*_*i*_) = 0*/*0. We use the following definition for singularity: *C*_*i*_ satisfying *ℱ*(*C*_*i*_) = ∞, −∞, or 0*/*0 at −∞ *< C*_*i*_ *<* ∞

## Supplementary Information

### SI 1 *A*_*nR*_(*C*_*i*_) [*a >* 0]

In this chapter, we examine the geometry of *A*_*nR*_(*C*_*i*_), which is defined by Eq. (12a) as follows:

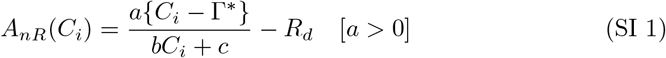

Note that *a >* 0 is assumed for *A*_*nR*_(*C*_*i*_) (Eq. 12a).

a. Derivative: 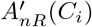 By differentiating Eq. (SI 1) with respect to *C*_*i*_, we have

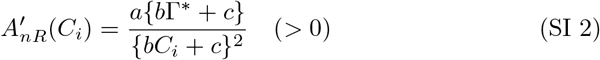 Therefore, *A*_*nR*_(*C*_*i*_) increases monotonically with *C*_*i*_ except at the singularity derived in the next section.
b. Singularity: 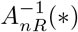 From Eq. (SI 1), the singularity is given by:

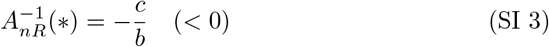 To clarify the signs of infinity on both sides of the singularity, an infinitesimal positive quantity, “*ϵ*(*>* 0)”, is introduced. Using “*ϵ*”, the neighborhoods on both sides of 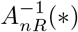 are expressed as:

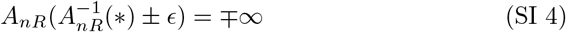
c. Root: 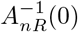 Setting *A*_*nR*_(*C*_*i*_) = 0 yields

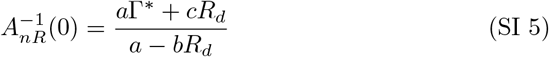 Therefore,

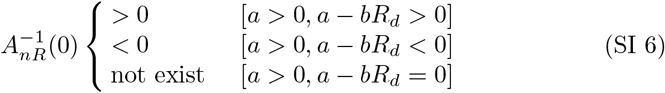
d. *A*_*nR*_(*±*∞) From Eq. (SI 1), *A*_*nR*_ (*±*∞) is given as:

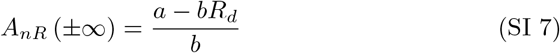 Therefore,

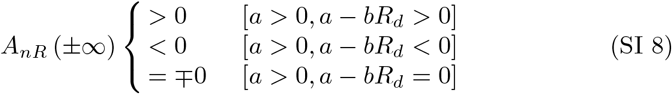 Eq. (SI 8) leads to:

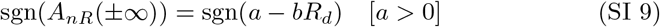
e. Inequality relations among 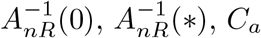 and 0 In the case where *a − bR*_*d*_ *>* 0, Eqs. (SI 3) and (SI 6) yield the following inequality:

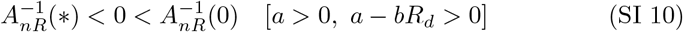 Thus, when *a >* 0, *a* − *bR*_*d*_ *>* 0 and 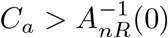,we have

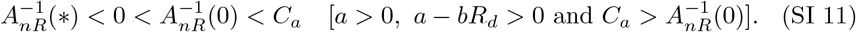 Similarly, when *a >* 0, *a*−*bR*_*d*_ *>* 0 and 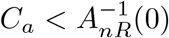,or when *a >* 0, *a*−*bR*_*d*_ *>* 0 and 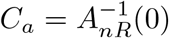,we have

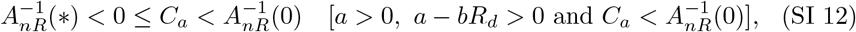

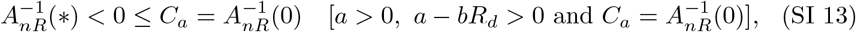 In the case where *a >* 0 and *a* − *bR*_*d*_ *<* 0, *A*_*nR*_(−∞) is negative (Eq. (SI 8)), while 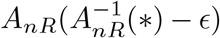 is positive (Eq. (SI 4)), indicating opposite signs at these two points. Since *A*_*nR*_(*C*_*i*_) increases monotonically with *C*_*i*_ for 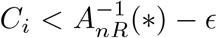 (Eq. (SI 2)), 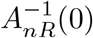 must exist between −∞ and 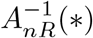.By combining this with *C*_*a*_ ≥ 0, we have

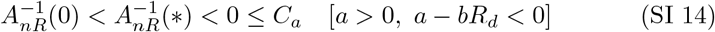 In the case where *a >* 0 and *a* − *bR*_*d*_ = 0, Eqs. (SI 3) and (SI 6), combined with *C*_*a*_ ≥ 0, yield

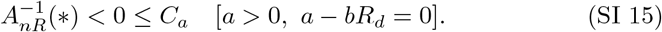 In summary, there are five patterns of inequality relations among 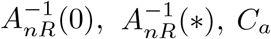 and 0 (Eqs. (SI 11) to (SI 15)). Table SI 1 summarizes these five patterns of inequality relations and the associated conditions. For the conditions, *A*_*nR*_(*±*∞) is used instead of *a* −*bR*_*d*_ since the signs are identical under [*a >* 0] (Eq. (SI 9)). To distinguish the conditions, the symbols [C1] to [C5] are used hereafter.
f. *A*_*nR*_(0) From Eq. (SI 1), *A*_*nR*_(0) is given by:

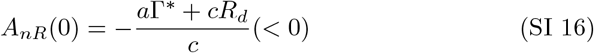
g. Expression of *A*_*nR*_(*C*_*i*_) Using 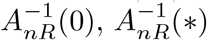,and *A*_*nR*_(0), *A*_*nR*_(*C*_*i*_) can be represented as follows:

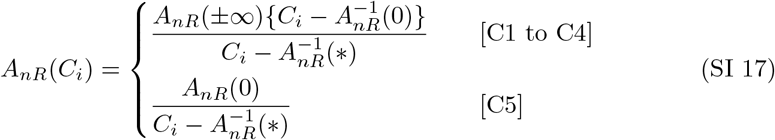
h. *A*_*nR*_(*C*_*a*_) From Eq. (SI 1), *A*_*nR*_(*C*_*a*_) is given by:

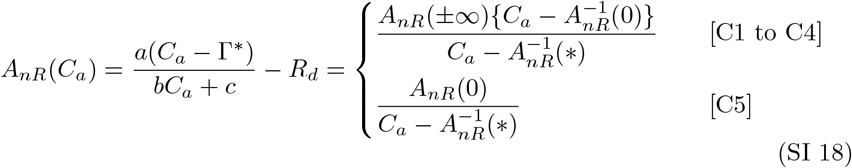 By using the signs of *A*_*nR*_(*±*∞) (Eq. (SI 8)), *A*_*nR*_(0) *<* 0 (Eq. (SI 16)), and the inequalities for each case (Table SI 1), the signs of *A*_*nR*_(*C*_*a*_) are given by:

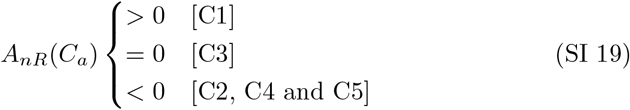
i. Geometry of *A*_*nR*_(*C*_*i*_) According to cases [C1] to [C5], *A*_*nR*_(*C*_*i*_) exhibits distinct geometries for each case. To differentiate these, we denote them as *A*_*nR*1_(*C*_*i*_) to *A*_*nR*5_(*C*_*i*_) for cases [C1] to [C5], respectively. Figures 1a to 1e display the curves of *A*_*nR*1_(*C*_*i*_) to *A*_*nR*5_(*C*_*i*_), and Tables SI 2 to SI 6 provide the sign charts of *A*_*nR*1_(*C*_*i*_) to *A*_*nR*5_(*C*_*i*_).

**Table SI 1:** Five patterns of conditions and inequality relations.

| Case | Condition | Inequality relation |
| --- | --- | --- |
| [C1] | $a > 0, A_{nR}(\pm\infty) > 0, C_a - A_{nR}^{-1}(0) > 0$ | $A_{nR}^{-1}(*) < 0 < A_{nR}^{-1}(0) < C_a$ |
| [C2] | $a > 0, A_{nR}(\pm\infty) > 0, C_a - A_{nR}^{-1}(0) < 0$ | $A_{nR}^{-1}(*) < 0 \leq C_a < A_{nR}^{-1}(0)$ |
| [C3] | $a > 0, A_{nR}(\pm\infty) > 0, C_a - A_{nR}^{-1}(0) = 0$ | $A_{nR}^{-1}(*) < 0 < C_a = A_{nR}^{-1}(0)$ |
| [C4] | $a > 0, A_{nR}(\pm\infty) < 0$ | $A_{nR}^{-1}(0) < A_{nR}^{-1}(*) < 0 \leq C_a$ |
| [C5] | $a > 0, A_{nR}(\pm\infty) = 0$ | $A_{nR}^{-1}(*) < 0 \leq C_a$ |

**Table SI 2:**
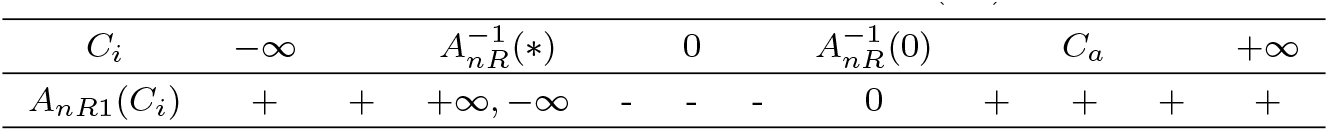
Sign chart for *A*_*nR*1_(*C*_*i*_)

**Table SI 3:** Sign chart for *A*_*nR*2_(*C*_*i*_)

|  |  |  |  |  |  |  |  |  |  |  |  |
| --- | --- | --- | --- | --- | --- | --- | --- | --- | --- | --- | --- |
| $C_i$ | $-\infty$ | | $A_{nR}^{-1}(*)$ | | | 0 | $C_a$ | $A_{nR}^{-1}(0)$ | | | $+\infty$ |
| $A_{nR2}(C_i)$ | + | + | $+\infty, -\infty$ | - | - | - | - | - | 0 | + | + |

**Table SI 4:** Sign chart for *A*_*nR*3_(*C*_*i*_)

|  |  |  |  |  |  |  |  |  |  |
| --- | --- | --- | --- | --- | --- | --- | --- | --- | --- |
| $C_i$ | $-\infty$ | $A_{nR}^{-1}(*)$ | | | 0 | $C_a = A_{nR}^{-1}(0)$ | | | $+\infty$ |
| $A_{nR3}(C_i)$ | + | + | $+\infty, -\infty$ | - | - | - | 0 | + | + |

**Table SI 5:**
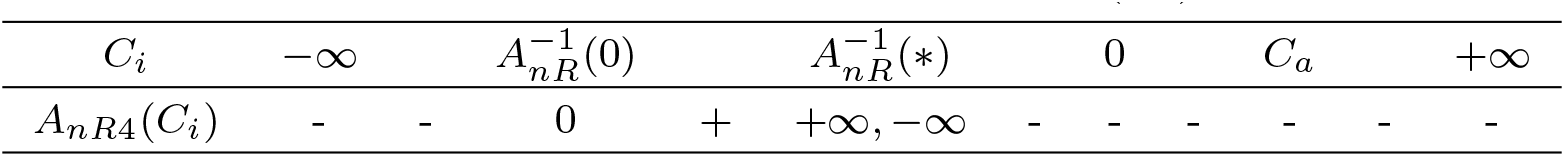
Sign chart for *A*_*nR*4_(*C*_*i*_)

**Table SI 6:**
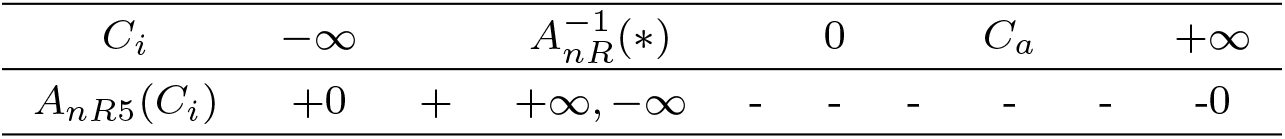
Sign chart for *A*_*nR*5_(*C*_*i*_)

### SI 2 *F*_*R*_(*C*_*i*_)

We examine the geometry of the function *F*_*R*_(*C*_*i*_), which is used when *a >* 0 under Rubisco/RuBP-limited conditions (Eq. (17)). Therefore, in analyzing *F*_*R*_(*C*_*i*_), we assume *a >* 0. Additionally, when *a >* 0, there are five different cases for the external variables, labeled [C1] through [C5] (Table SI 1). Accordingly, the geometries of *F*_*R*_(*C*_*i*_) for [C1] to [C5] are examined in this chapter.

From Eq. (17), the function *F*_*R*_(*C*_*i*_) is given by :

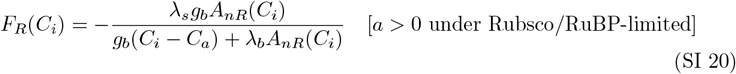

#### SI 2.1 Expression of *F*_*R*_(*C*_*i*_)

Using Eq. (SI 17), *F*_*R*_(*C*_*i*_) can be rewritten based on Eq. (SI 20) as follows:

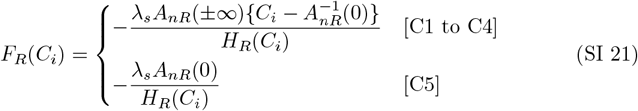

where *H*_*R*_(*C*_*i*_) is given by a quadratic equation in terms of *C*_*i*_, as follows:

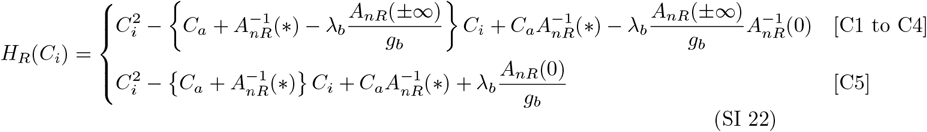

#### SI 2.2 *H*_*R*_(*C*_*i*_)

In this section, we investigate *H*_*R*_(*C*_*i*_) as defined in Eq. (SI 22).

(a) Root: 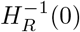 The equation *H*_*R*_(*C*_*i*_) = 0 (Eq. (SI 22)) is a quadratic polynomial in *C*_*i*_, expressed as 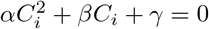,where the coefficients are given by:

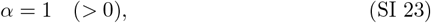

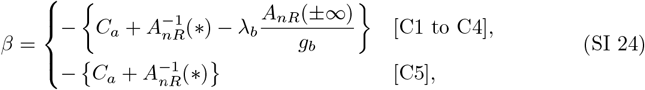

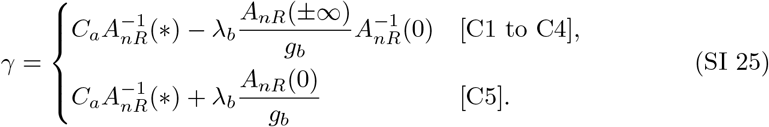

Table SI 1 shows 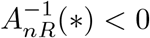 under [C1] to 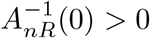 and 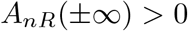 under [C1] to 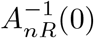 and *A*_*nR*_(*±*∞) *<* 0 under [C4]; and *A*_*nR*_(0) *<* 0 under [C5] (Eq. (SI 16)). Based on Eq. (SI 25), the sign of *γ* is therefore determined as:

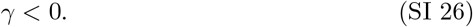

Given *α >* 0 (Eq. (SI 23)) and *γ <* 0 (Eq. (SI 26)), *H*_*R*_(*C*_*i*_) = 0 has two real roots, one positive and one negative, given by:

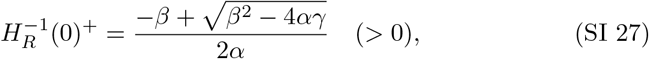

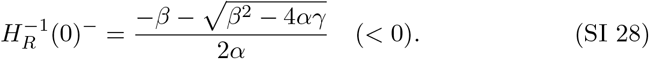

The superscripts “+ “and “− “denote the positive and negative roots, respectively.

Using 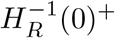 and 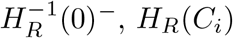 can be factored as

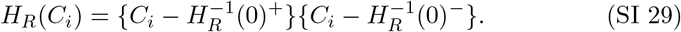

From the relationships between roots and coefficients, we obtain

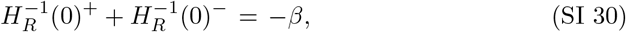

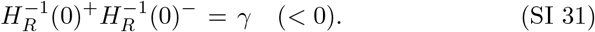

Note that *α* = 1 (Eq. (SI 23)) is assumed in Eqs. (SI 30) and (SI 31). Since *H*_*R*_(*C*_*i*_) is a convex quadratic function, we have:

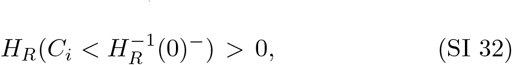

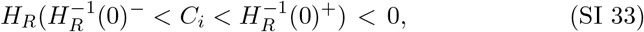

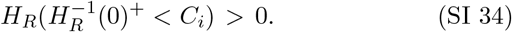

(b) 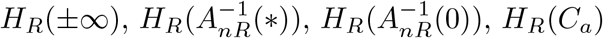,and *H*_*R*_(0)

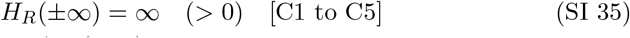

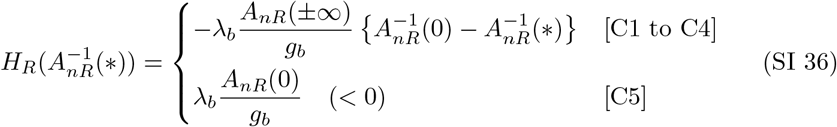

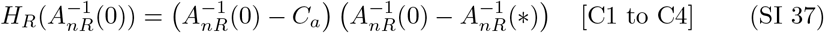

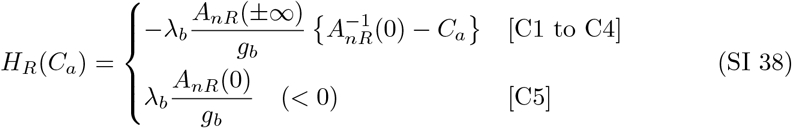

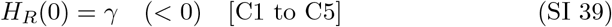
(c) Inequality relation among 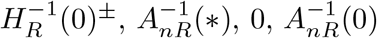,and *C*_*a*_

We examine inequality relations among 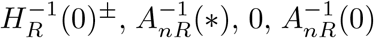,and *C*_*a*_ for each case from [C1] to [C5].

In case [C1], we have 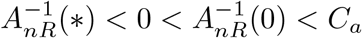 (Eq. (SI 11)). From Eqs. (SI 35) to (SI 39), and Table SI 1, it follows that:

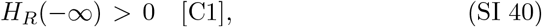

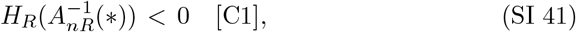

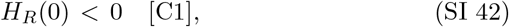

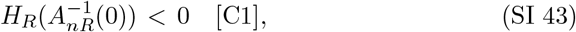

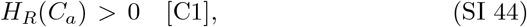

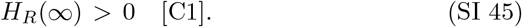

Since 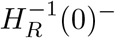 and 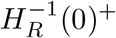 must lie between points where *H*_*R*_(*C*_*i*_) changes sign, we derive the following inequality relation:

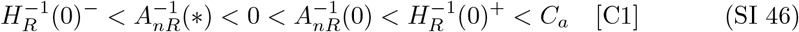

In case [C2], we have 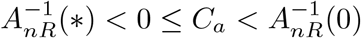 (Eq. (SI 12)). From Eqs. (SI 35) to (SI 39), and Table SI 1, it follows that:

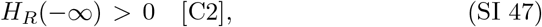

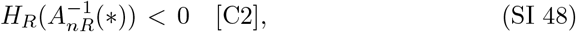

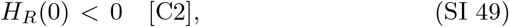

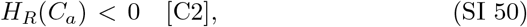

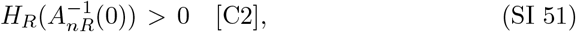

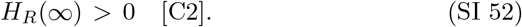

Following the same reasoning, we obtain the following inequality relation:

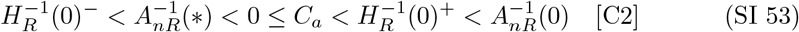

In case [C3], we have 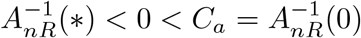 (Eq. (SI 13)). From Eqs. (SI 35) to (SI 39), and Table SI 1, we find:

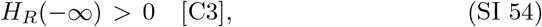

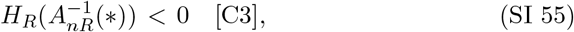

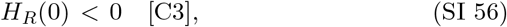

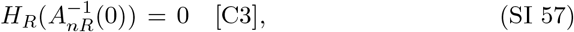

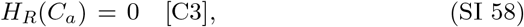

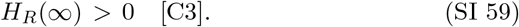

Since 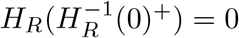,we find 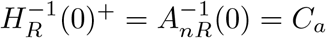,Thus, we have:

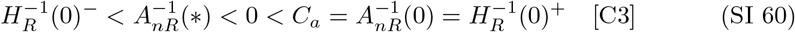

In case [C4], 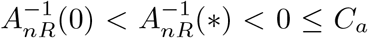 (Eq. (SI 14)). From Eqs. (SI 35) to (SI 39), Table SI 1, we obtain:

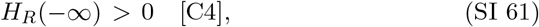

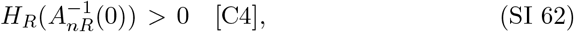

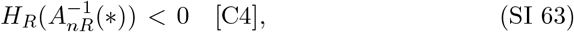

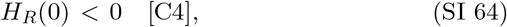

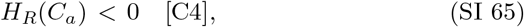

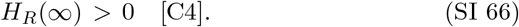

Therefore, following the same reasoning, we derive the inequality relation:

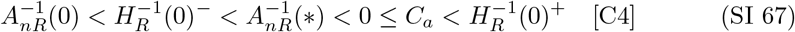

In case [C5], 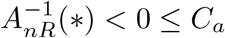 (Eq. (SI 15)). From Eqs. (SI 35) to (SI 39), we have:

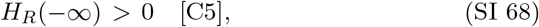

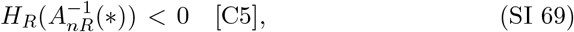

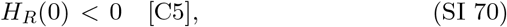

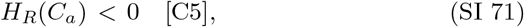

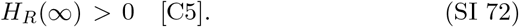

Thus, from the same discussion as above, we have the following inequality relation:

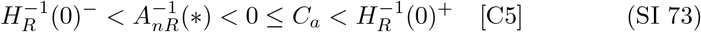

Table SI 7 summarizes the inequality relationships for *H*_*R*_(*C*_*i*_) across cases [C1] to [C5].

(d) Geometry of *H*_*R*_(*C*_*i*_)

Figure SI 1 shows the curve of *H*_*R*_(*C*_*i*_) for case [C1]. The shapes of the curves for the other cases are similar to that of [C1], but the inequalities among points such as 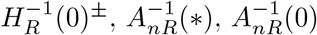 and *C*_*a*_ vary for each case. Tables SI 8 to SI 12 present the sign charts of *H*_*R*_(*C*_*i*_) for cases [C1] to [C5].

**Table SI 7:**
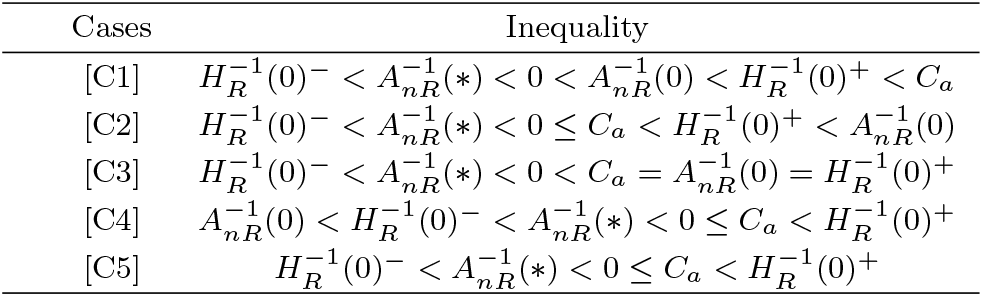
Inequalities for *H*_*R*_(*C*_*i*_)

**Fig. SI 1:**
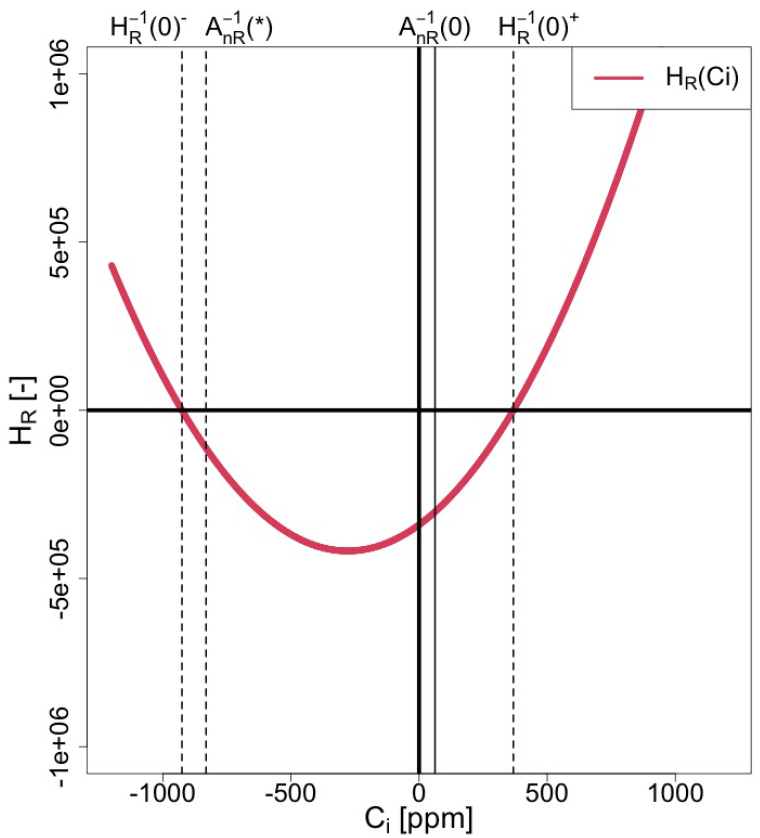
*H*_*R*_(*C*_*i*_) [C1]

**Table SI 8:**
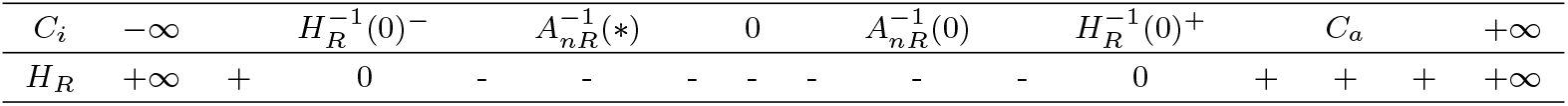
Sign chart of *H*_*R*_(*C*_*i*_) [C1].

**Table SI 9:**
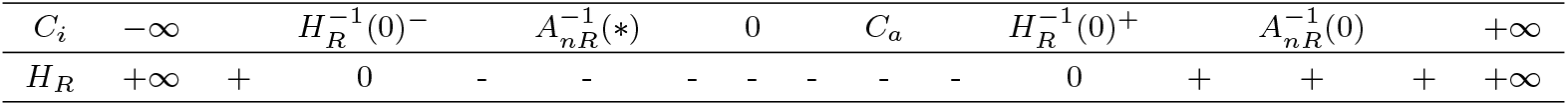
Sign chart of *H*_*R*_(*C*_*i*_) [C2].

|  |  |  |  |  |  |  |  |  |  |  |  |  |  |  |  |
| --- | --- | --- | --- | --- | --- | --- | --- | --- | --- | --- | --- | --- | --- | --- | --- |
| $C_i$ | $-\infty$ | | $H_R^{-1}(0)^-$ | | $A_{nR}^{-1}(*)$ | | 0 | | $C_a$ | | $H_R^{-1}(0)^+$ | | $A_{nR}^{-1}(0)$ | | $+\infty$ |
| $H_R$ | $+\infty$ | + | 0 | - | - | - | - | - | - | - | 0 | + | + | + | $+\infty$ |

**Table SI 10:** Sign chart of *H*_*R*_(*C*_*i*_) [C3].

|  |  |  |  |  |  |  |  |  |  |  |  |
| --- | --- | --- | --- | --- | --- | --- | --- | --- | --- | --- | --- |
| $C_i$ | $-\infty$ | | $H_R^{-1}(0)^-$ | | $A_{nR}^{-1}(*)$ | | 0 | | $C_a = H_R^{-1}(0)^+ = A_{nR}^{-1}(0)$ | | $+\infty$ |
| $H_R$ | $+\infty$ | + | 0 | - | - | - | - | - | 0 | + | $+\infty$ |

**Table SI 11:** Sign chart of *H*_*R*_(*C*_*i*_) [C4].

|  |  |  |  |  |  |  |  |  |  |  |  |  |  |  |  |
| --- | --- | --- | --- | --- | --- | --- | --- | --- | --- | --- | --- | --- | --- | --- | --- |
| $C_i$ | $-\infty$ | | $A_{nR}^{-1}(0)$ | | $H_R^{-1}(0)^-$ | | $A_{nR}^{-1}(*)$ | | 0 | | $C_a$ | | $H_R^{-1}(0)^+$ | | $+\infty$ |
| $H_R$ | $+\infty$ | + | + | + | 0 | - | - | - | - | - | - | - | 0 | + | $+\infty$ |

**Table SI 12:** Sign chart of *H*_*R*_(*C*_*i*_) [C5].

|  |  |  |  |  |  |  |  |  |  |  |  |  |  |
| --- | --- | --- | --- | --- | --- | --- | --- | --- | --- | --- | --- | --- | --- |
| $C_i$ | $-\infty$ | | $H_R^{-1}(0)^-$ | | $A_{nR}^{-1}(*)$ | | 0 | | $C_a$ | | $H_R^{-1}(0)^+$ | | $+\infty$ |
| $H_R$ | $+\infty$ | + | 0 | - | - | - | - | - | - | - | 0 | + | $+\infty$ |

#### SI 2.3 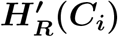

By differentiating *H*_*R*_(*C*_*i*_) with respect to *C*_*i*_ under cases [C1] to [C5] (Eq. (SI 22)), we have:

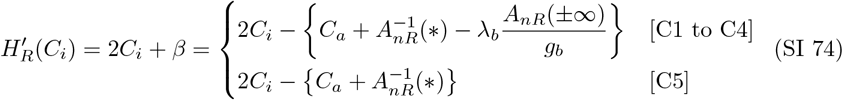

(a) 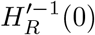 From Eq. (SI 74), we obtain

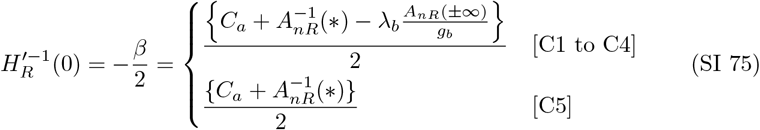 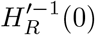 represents the value of *C*_*i*_ at which the slope of *H*_*R*_(*C*_*i*_) is zero. Since 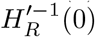 must lie between the two roots of *H*_*R*_(*C*_*i*_) = 0, it satisfies the following inequality:

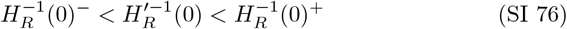
(b) Signs of 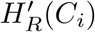 Since *H*_*R*_(*C*_*i*_) is a convex quadratic function of *C*_*i*_, the sign of 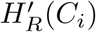(*C*_*i*_) changes from negative to positive at 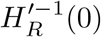.Hence, we have:

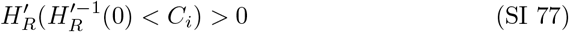

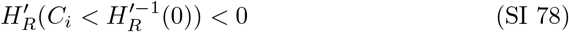 From Eqs. (SI 76) to (SI 78), we obtain:

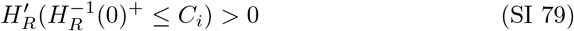

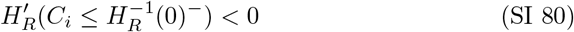

#### SI 2.4 *F*_*R*_(*C*_*i*_)

*F*_*R*_(*C*_*i*_) under cases [C1] to [C5] is given by Eq. (SI 21):

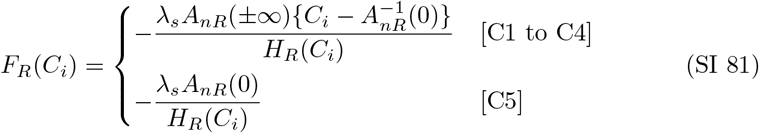

(a) Singularities: 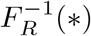 Eq. (SI 81) shows that *F*_*R*_(*C*_*i*_) has singularities at points *C*_*i*_ where *H*_*R*_(*C*_*i*_) = 0. Thus, we can express the singularities as:

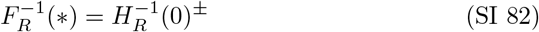 To differentiate between the two singularities, we denote them as:

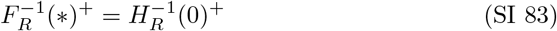

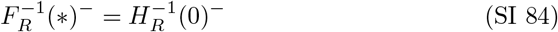 The behavior of *F*_*R*_(*C*_*i*_) in the neighborhood of 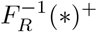 is expressed as

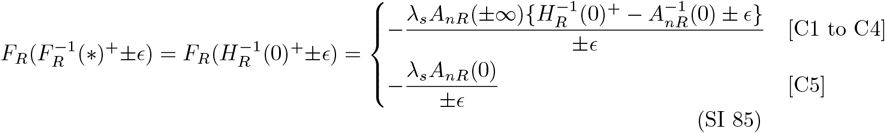 In the case of [C1], *A*_*nR*_(*±*∞) *>* 0 (Eq. (SI 8)) and 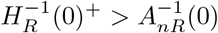 (Eq. (SI 46)). Therefore, we obtain 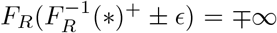 for [C1]. Similarly, for cases [C2] to [C5], we have:

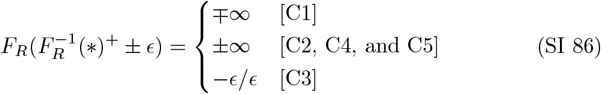 Following similar reasoning, we find:

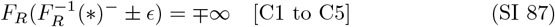 In case [C3], 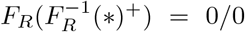 (Eq. (SI 86)), indicating that *F*_*R*_(*C*_*i*_) at 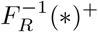 is indeterminate under [C3], as follows:

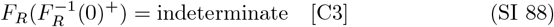
(b) Roots: 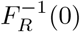 From Eqs. (SI 81), we have:

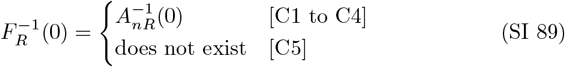 In case [C3], 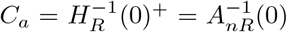 under [C3] (SI 60). Combining this with Eqs. (SI 83) and (SI 89), we have:

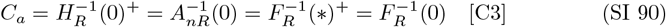
(c) *F*_*R*_(*±*∞) and 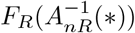

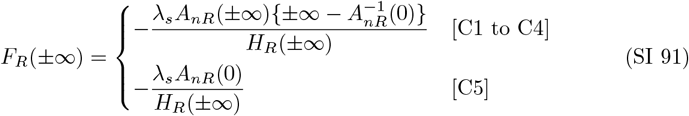 Noting that the denominator of *F*_*R*_(*C*_*i*_) is a quadratic function of *C*_*i*_ (Eq. (SI 22)), while the numerator is linear for [C1 to C4] or constant for [C5], we find:

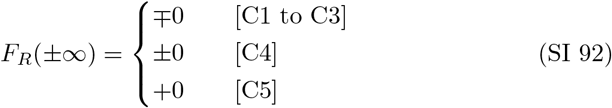

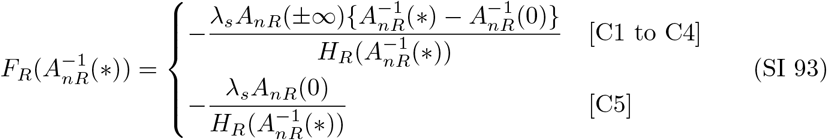 From the inequalities in Section SI 2.2 (c), we obtain:

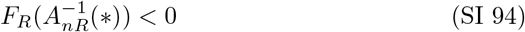
(d) Geometry of *F*_*R*_(*C*_*i*_) According to the distinct geometries of *A*_*nR*_(*C*_*i*_) and *H*_*R*_(*C*_*i*_) for cases [C1] to [C5] (Sections SI 1 and SI 2.2), *F*_*R*_(*C*_*i*_) also exhibits distinct geometries for each case. To differentiate these, we denote them as *F*_*R*1_(*C*_*i*_) to *F*_*R*5_(*C*_*i*_) for cases [C1] to [C5], respectively. Figures 2a to 2e display the curves of *F*_*R*1_(*C*_*i*_) to *F*_*R*5_(*C*_*i*_), and Tables SI 13 to SI 17 provide the sign charts of *F*_*R*1_(*C*_*i*_) to *F*_*R*5_(*C*_*i*_) as well as the signs of the numerator and denominator. In the figures and tables, we use the inequality relations between the points for cases [C1] to [C5] (Table SI 7), and two equations: 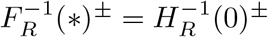(Eqs. (SI 83) and (SI 84)) and 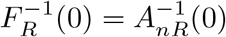 (Eq. (SI 89)).
(e) Possible Range of Appropriate Solutions for *F*_*R*_(*C*_*i*_)

Based on the geometries of *F*_*R*_(*C*_*i*_) identified in the previous section, the possible range of appropriate solutions is determined by selecting the interval of *C*_*i*_ where *F*_*R*_(*C*_*i*_) *>* 0 and *C*_*i*_ *>* 0. Table SI 18 summarizes the possible ranges of appropriate solutions for *F*_*R*_(*C*_*i*_).

**Table SI 13:**
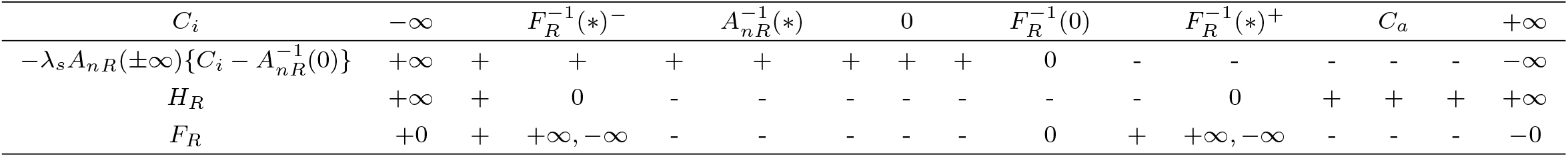
Sign chart of *F*_*R*1_(*C*_*i*_)

**Table SI 14:**
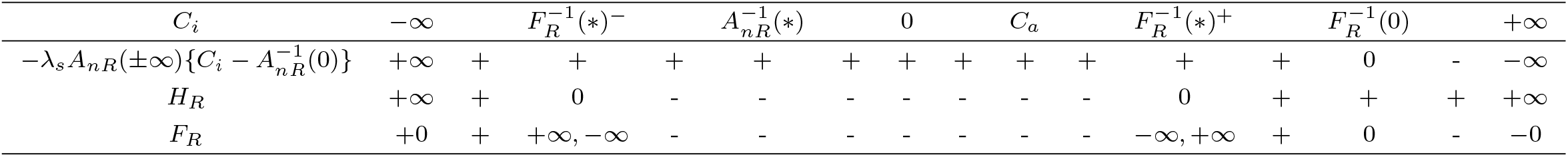
Sign chart of *F*_*R*2_(*C*_*i*_)

**Table SI 15:**
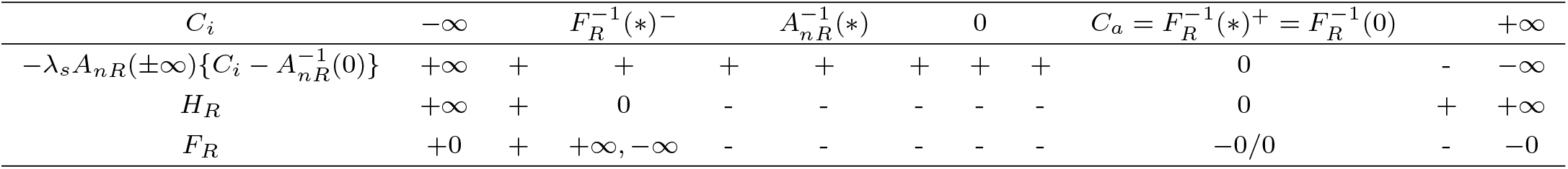
Sign chart of *F*_*R*3_(*C*_*i*_)

**Table SI 16:**
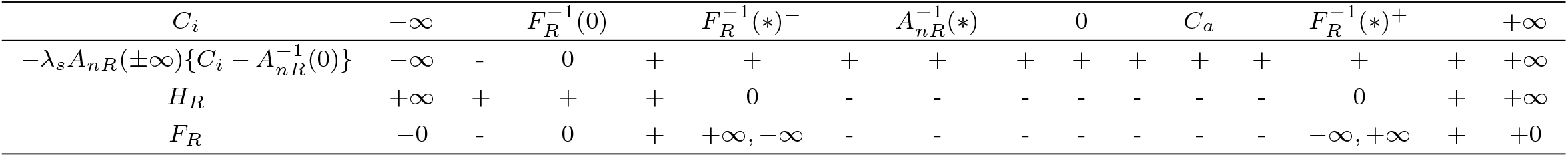
Sign chart of *F*_*R*4_(*C*_*i*_)

**Table SI 17:** Sign chart of *F*_*R*5_(*C*_*i*_)

| $C_i$ | $-\infty$ | $F_R^{-1}(*)^-$ | $A_{nR}^{-1}(*)$ | 0 | $C_a$ | $F_R^{-1}(*)^+$ | $+\infty$ |
| --- | --- | --- | --- | --- | --- | --- | --- |
| $-\lambda_s A_{nR}(0)$ | + | + | + | + | + | + | + |
| $H_R$ | $+\infty$ | 0 | - | - | - | 0 | $+\infty$ |
| $F_R$ | +0 | $+\infty, -\infty$ | - | - | - | $-\infty, +\infty$ | +0 |

**Table SI 18:** Possible range of appropriate solutions for *F*_*R*_(*C*_*i*_)

| [C] | $F_R(C_i)$ | Range |
| --- | --- | --- |
| [C1] | $F_{R1}(C_i)$ | $F_R^{-1}(0) < C_i < F_R^{-1}(*)^+$ |
| [C2] | $F_{R2}(C_i)$ | $F_R^{-1}(*)^+ < C_i < F_R^{-1}(0)$ |
| [C3] | $F_{R3}(C_i)$ | $C_i = C_a = F_R^{-1}(*)^+ = F_R^{-1}(0)$ |
| [C4] | $F_{R4}(C_i)$ | $F_R^{-1}(*)^+ < C_i$ |
| [C5] | $F_{R5}(C_i)$ | $F_R^{-1}(*)^+ < C_i$ |

#### SI 2.5 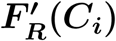

Differentiating *F*_*R*_(*C*_*i*_) with respect to *C*_*i*_ for cases [C1] to [C5] (Eq. (SI 81)) gives the derivative of *F*_*R*_(*C*_*i*_) as follows:

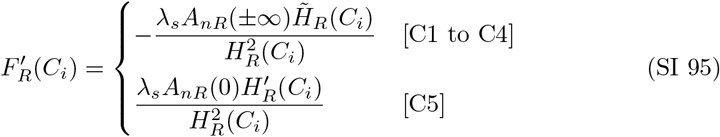

where 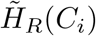 is defined as:

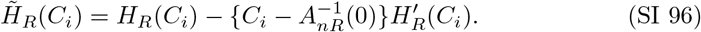

We focus exclusively on the possible range of appropriate solutions in this section (Table SI 18).

#### SI 2.5.1 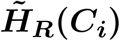

Using Eqs. (SI 22) and (SI 74), 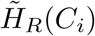 can be rewritten as follows:

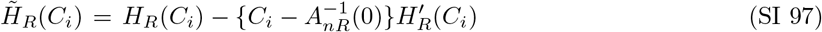

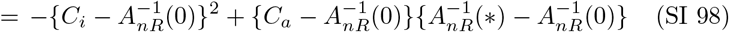

Eq. (SI 98) shows that 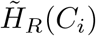 is a concave quadratic function of *C*_*i*_ and attains its maximum at 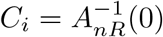.

(a) 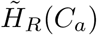

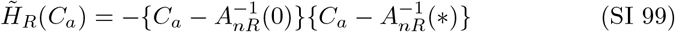 Considering the inequalities in Table SI 7, Eq. (SI 99) gives the following inequalities:

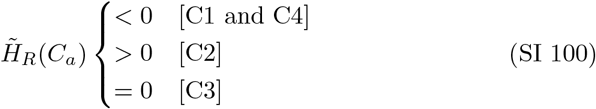
(b) 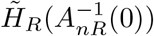

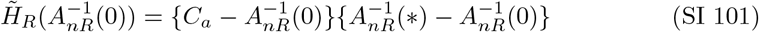 By using the inequalities in Table SI 7, we obtain

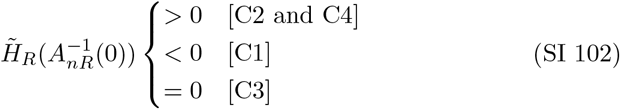
(c) 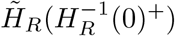

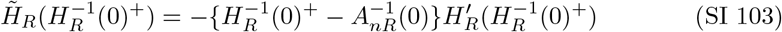 By using the inequalities in Table SI 7 and 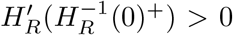 (Eq. (SI 79)), we obtain:

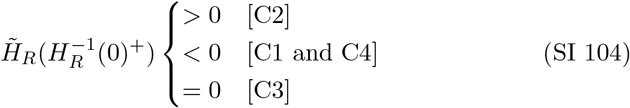
(d) Sign of 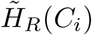 Note that 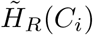 is a concave quadratic function of *C*_*i*_ that attains its maximum at 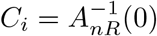.

In case [C1], since 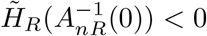(Eq. (SI 102)), we have:

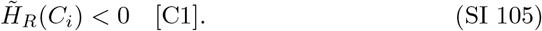

For case [C2], given that 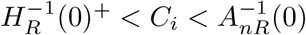 (Table SI 7), 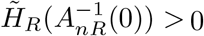 (Eq. (SI 102)), and 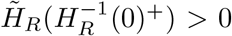 (Eq. (SI 104)), it follows from Eq. (SI 98) that:

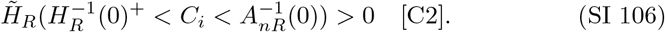

By combining Eq. (SI 106) with 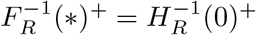 (Eq. (SI 83)) and 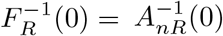 (Eq. (SI 89)), we have:

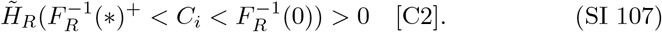

For case [C4], given that 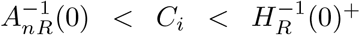 (Table SI 7) and 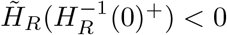 (Eq. (SI 104)), it follows from Eq. (SI 98) that:

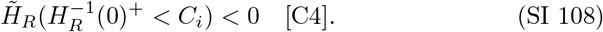

Similarly to case [C2], by combining Eq. (SI 108) with 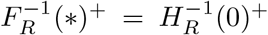 (Eq. (SI 83)), we have:

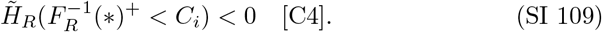

#### SI 2.5.2 Sign of 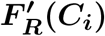

For case [C1], since *A*_*nR*_(*±*∞) *>* 0 (Eq. SI 7) and 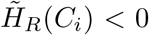 (Eq. SI 105), Eq. (SI 95) gives:

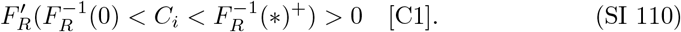

For case [C2], given that *A*_*nR*_(*±*∞) *>* 0 (Eq. SI 7) and 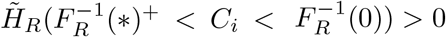 (Eq. SI 107), Eq. (SI 95) indicates:

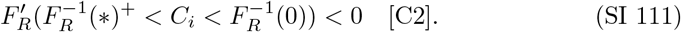

For case [C4], given that *A*_*nR*_(*±*∞) *<* 0 (Eq. SI 7) and 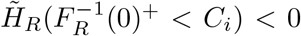 (Eq. SI 109), Eq. (SI 95) results in:

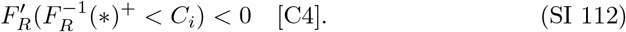

For case [C5], since *A*_*nR*_(0) *<* 0 (Eq. SI 16) and 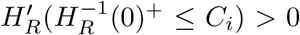 (Eq. SI 79), Eq. (SI 95) leads to:

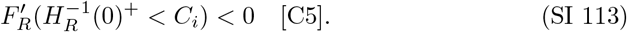

By combining Eq. (SI 113) with 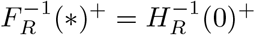 (Eq. SI 83), we obtain:

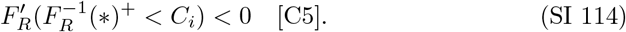

### SI 3 *F*_*R*0_(*C*_*i*_)

In this chapter, we explore *F*_*R*0_(*C*_*i*_), which is given by:

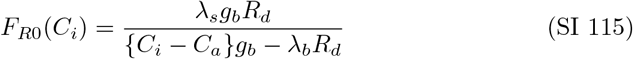

a. Derivative: 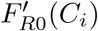

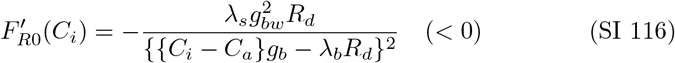 Thus, *F*_*R*0_(*C*_*i*_) monotonically decreases with *C*_*i*_, except at the singularity obtained in the next section.
b. Singularity: 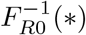 From Eq. (SI 115), 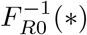 is given by:

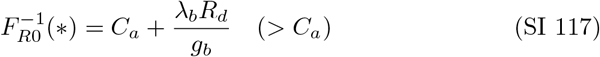 The behavior of *F*_*R*0_(*C*_*i*_) in the neighborhood of 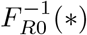 is given by:

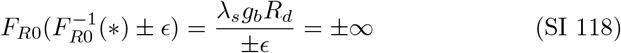
c. *F*_*R*0_(*±*∞) and *F*_*R*0_(*C*_*a*_)

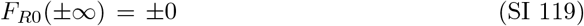

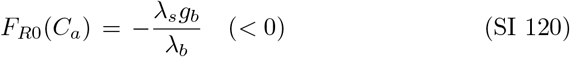
d. Geometry of *F*_*R*0_(*C*_*i*_) Figure 2f and Table SI 19 show the curve and sign chart of *F*_*R*0_(*C*_*i*_).
e. Possible range of appropriate solutions for *F*_*R*0_(*C*_*i*_) Based on the geometries of *F*_*R*0_(*C*_*i*_) identified in the previous section, the possible range of appropriate solutions can be determined by selecting the interval of *C*_*i*_ where *F*_*R*0_(*C*_*i*_) *>* 0 and *C*_*i*_ *>* 0. Table SI 20 summarizes the possible range of appropriate solutions for *F*_*R*0_(*C*_*i*_).

**Table SI 19:**
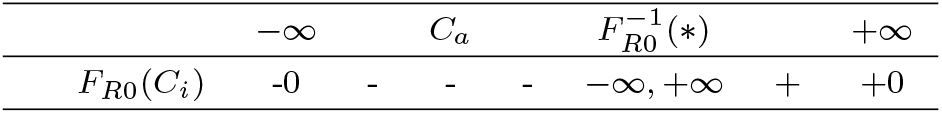
Sign chart of *F*_*R*0_(*C*_*i*_)

**Table SI 20:**
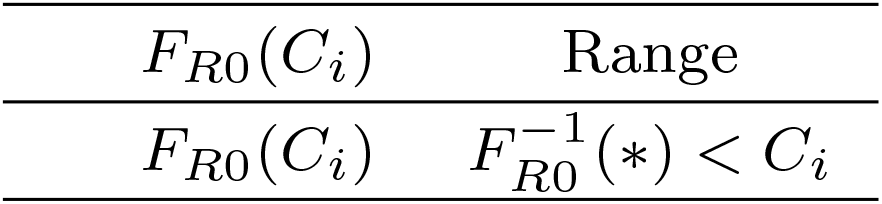
Possible range of appropriate solutions for *F*_*R*0_(*C*_*i*_)

### SI 4 *F*_*P*_ (*C*_*i*_)

In this chapter, we investigate the geometry of *F*_*P*_ (*C*_*i*_), which is given by:

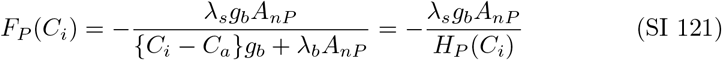

where *H*_*P*_ (*C*_*i*_) is given by:

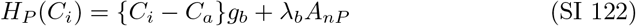

#### SI 4.1 *H*_*P*_ (*C*_*i*_)

(a) Root: 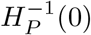 Setting *H*_*P*_ (*C*_*i*_) = 0, we obtain:

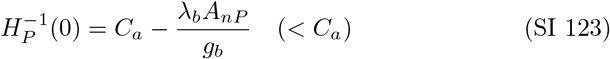 Therefore, we have:

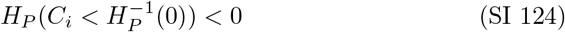

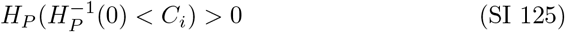 In the case of Product-limited conditions, from Eqs. (6), (8), and (12c), we find:

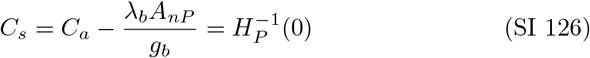 Since *C*_*s*_ *<* 0 is physically impossible, we require *C*_*s*_ *>* 0. Thus, we have:

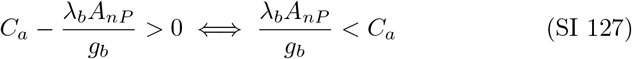 Eq. (SI 127) provides the lower limit of *C*_*a*_. Using Eqs. (SI 123), (SI 126), and (SI 127), we obtain:

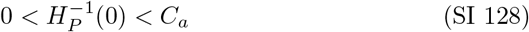
(b) *H*_*P*_ (0) By substituting *C*_*i*_ = 0 into Eq. (SI 122), we have:

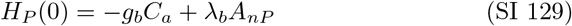 Using Eq. (SI 127), we find:

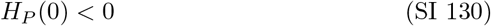
(c) 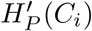 Differentiating *H*_*P*_ (*C*_*i*_) (Eq. (SI 122)) with respect to *C*_*i*_, we obtain:

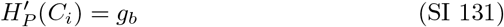

#### SI 4.2 *F*_*P*_ (*C*_*i*_)

(a) Derivative: 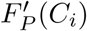

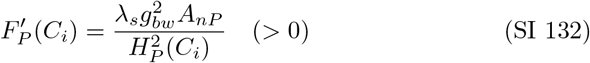 Thus, *F*_*P*_ (*C*_*i*_) monotonically increases with *C*_*i*_, except at the singularity calculated in the next section.
(b) Singularity: 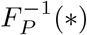 From Eq. (SI 121), 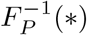 is given by:

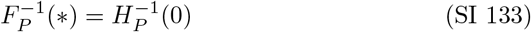
(c) *F*_*P*_ (*±*∞), *F*_*P*_ (0), and *F*_*P*_ (*C*_*a*_)

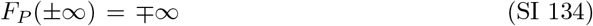

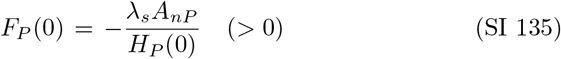

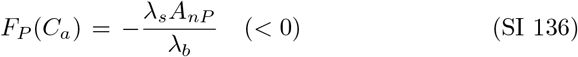
(d) Geometry of *F*_*P*_ (*C*_*i*_) Figure 2g and Table SI 21 show the curve and sign chart of *F*_*P*_ (*C*_*i*_).
(e) Possible range of appropriate solutions for *F*_*P*_ (*C*_*i*_) Based on the geometries of *F*_*P*_ (*C*_*i*_) identified in the previous section, the possible range of appropriate solutions can be determined by selecting the interval of *C*_*i*_ where *F*_*P*_ (*C*_*i*_) *>* 0 and *C*_*i*_ *>* 0. Table SI 22 summarizes the possible range of appropriate solutions for *F*_*P*_ (*C*_*i*_).

**Table SI 21:**
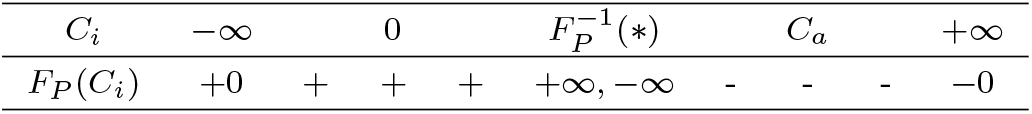
Geometry of *F*_*P*_ (*C*_*i*_)

**Table SI 22:** Possible range of appropriate solutions for *F*_*P*_ (*C*_*i*_)

| $F_P(C_i)$ | Range |
| --- | --- |
| $F_P(C_i)$ | $0 < C_i < F_P^{-1}(*)$ |

### SI 5 *Z*_*R*_(*C*_*i*_)

We analyze the geometry of *Z*_*R*_(*C*_*i*_), which represents *Z*(*C*_*i*_) under case [C1] and *e*_*a*_ *< e*_*i*_ (Table 3). It is defined as:

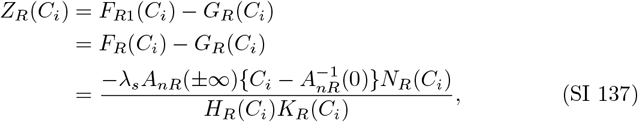

where *N*_*R*_(*C*_*i*_) is expressed as:

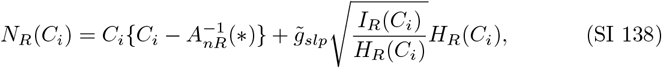

with *I*_*R*_(*C*_*i*_) and *K*_*R*_(*C*_*i*_) defined as:

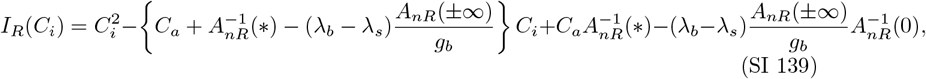

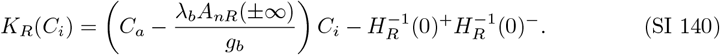

The function *N*_*R*_(*C*_*i*_) is real for *C*_*i*_ satisfying:

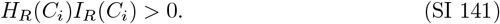

Hence, Eq. (SI 141) is assumed throughout this section. Additionally, as case [C1] is a condition for *Z*_*R*_(*C*_*i*_) (Table 4), [C1] is assumed to all analyses in this chapter.

#### SI 5.1 *I*_*R*_(*C*_*i*_)

(a) 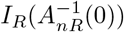 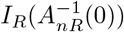is given by:

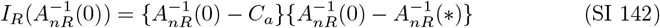 Since 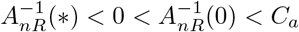 under [C1] (Table SI 1), we have:

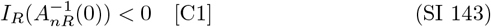 Thus, *I*_*R*_(*C*_*i*_) has a negative value at 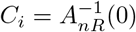.
(b) Root: 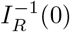 Setting *I*_*R*_(*C*_*i*_) = 0 yields a quadratic polynomial equation with respect to *C*_*i*_ in the form 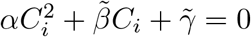,where the coefficients are given by:

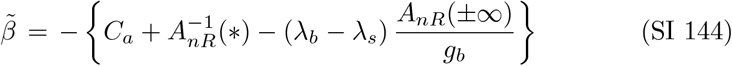

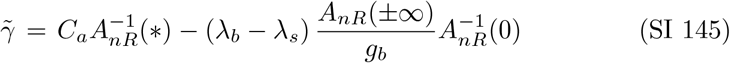 Note that *α* = 1 (Eq. (SI 23)). Eqs. (SI 139) and (SI 143) show that *I*_*R*_(*C*_*i*_) is a convex quadratic function of *C*_*i*_ and has a negative value at 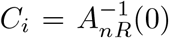. Therefore, *I*_*R*_(*C*_*i*_) = 0 has two real solutions, expressed as:

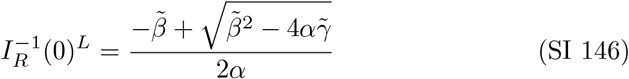

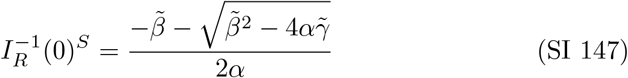 The two roots satisfy the following inequality:

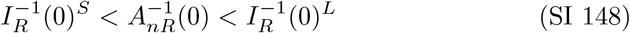 Since 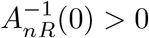 under [C1] (Table SI 1), we have:

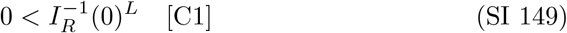 Using 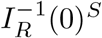 and 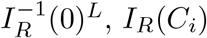 can be expressed as:

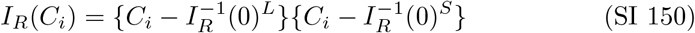 From the relationship between solutions and coefficients for quadratic equations, we have:

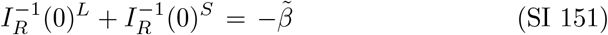

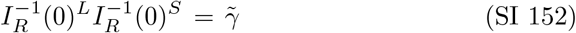 Note that *α* = 1 (Eq. (SI 23)) was considered in Eqs. (SI 151) and (SI 152). From Eqs. (SI 149) and (SI 152), the signs of 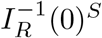 under [C1] are given as:

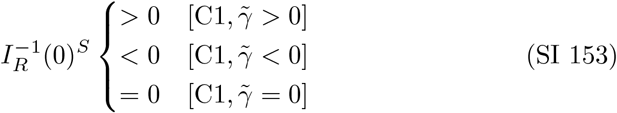 Eq. (SI 153) shows that the signs of 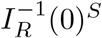 are the same as those of 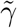 under [C1], as follows:

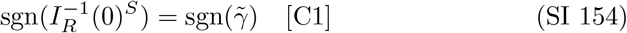 Considering that *I*_*R*_(*C*_*i*_) is convex with *α* = 1, we have:

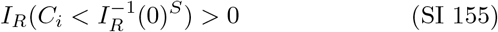

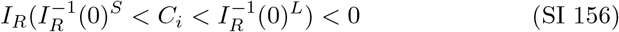

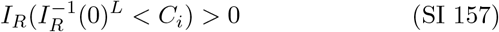
(c) 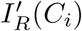 By differentiating *I*_*R*_(*C*_*i*_) with respect to *C*_*i*_, we have:

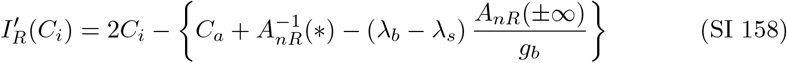 Since *I*_*R*_(*C*_*i*_) is a convex quadratic function of *C*_*i*_ (Eq. (SI 139)) and *I*_*R*_(*C*_*i*_) = 0 has two real solutions (Eqs. (SI 146) and (SI 147)), we have the following inequalities:

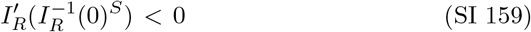

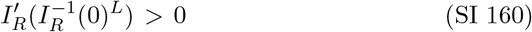
(d) 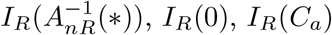,and *I*_*R*_(*±*∞)

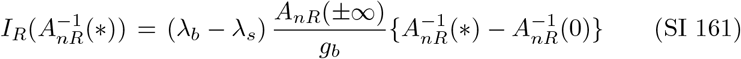

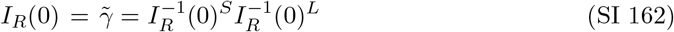

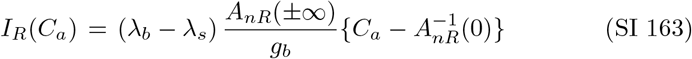

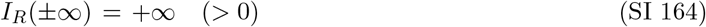 Since *A*_*nR*_(*±*∞) *>* 0 and 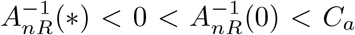 under [C1] (Table SI 1), we have:

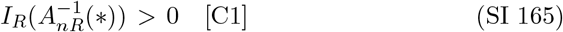

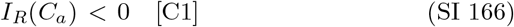
(e) Inequality relations between 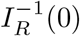 and other points In case 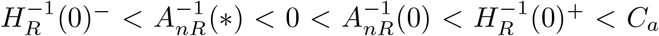 (Table SI 7).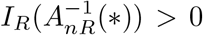 (Eq. (SI 165)) and 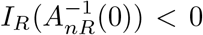 (Eq. (SI 143)), indicating opposite signs at these two points. Similarly, *I*_*R*_(*C*_*a*_) *<* 0 (Eq. (SI 166)) and *I*_*R*_(+∞) *>* 0 (Eq. (SI 164)), indicating opposite signs at these points. Considering the inequality between two solutions (Eq. (SI 148)) and the signs of 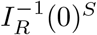 (Eq. (SI 153)), we have the following possible inequalities:

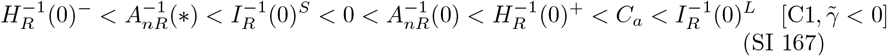

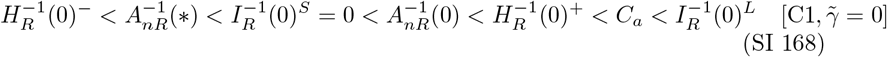

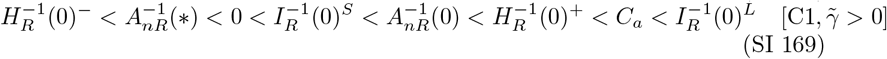
(f) Geometry of *I*_*R*_(*C*_*i*_) Figure SI 2 shows the curve of *I*_*R*_(*C*_*i*_) under 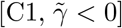.The curves of *I*_*R*_(*C*_*i*_) for other cases are the same as for 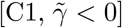,except for the inequality relationships between points. Tables SI 23 to SI 25 show the sign charts of *I*_*R*_(*C*_*i*_) for each case.

**Fig. SI 2:**
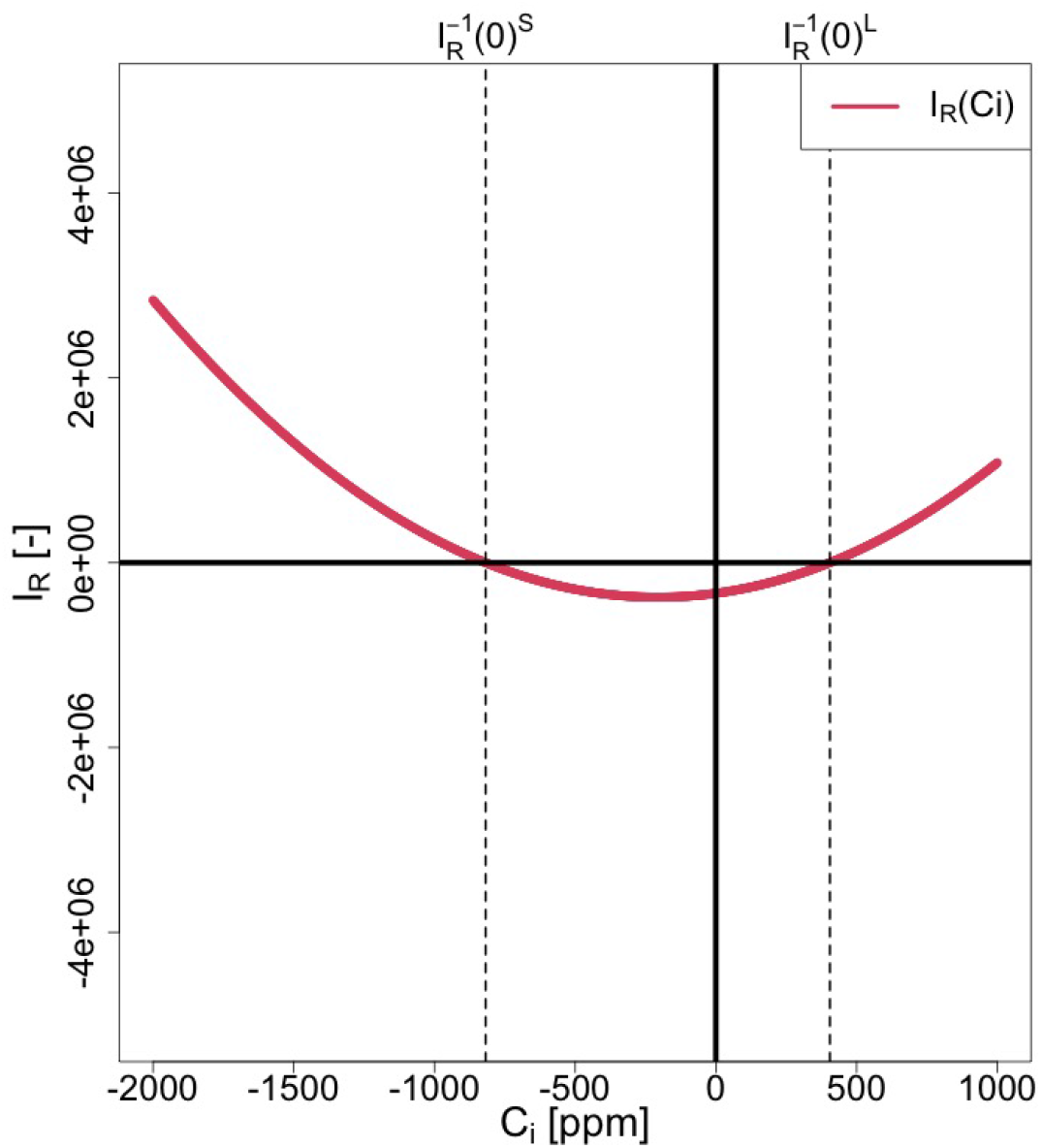
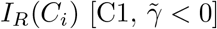

**Table SI 23:**
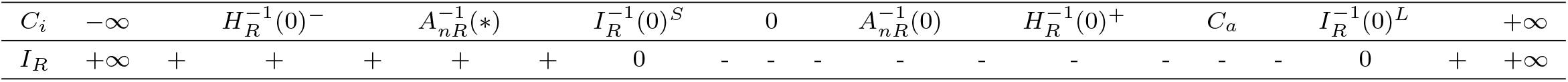
Sign Chart of *I*_*R*_(*C*_*i*_) under 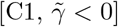.

**Table SI 24:**
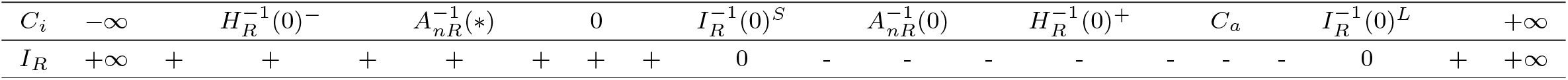
Sign Chart of *I*_*R*_(*C*_*i*_) under 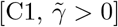.

**Table SI 25:**
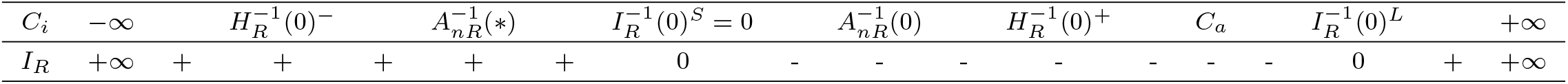
Sign Chart of *I*_*R*_(*C*_*i*_) under 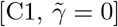.

#### SI 5.2 *K*_*R*_(*C*_*i*_)

This section examines the function *K*_*R*_(*C*_*i*_), which is given from Eq. (SI 140) by:

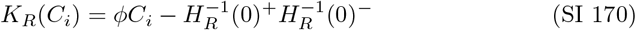

where *ϕ* is defined as

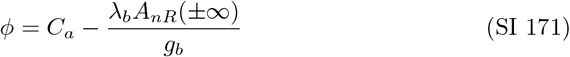

(a) Derivative: 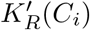

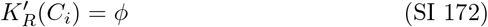 Thus *K*_*R*_(*C*_*i*_) is a linear function of *C*_*i*_ with a slope of *ϕ*.
(b) Root: 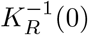 If *ϕ*≠ 0, *K*_*R*_(*C*_*i*_) = 0 yields:

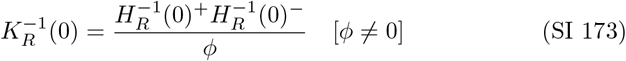 Given 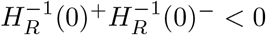 (Eq. (SI 31)), we obtain:

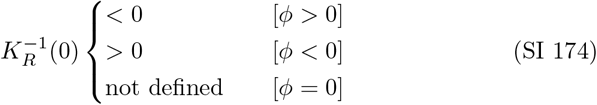 Since *K*_*R*_(*C*_*i*_) is a linear function with slope *ϕ*, we have:

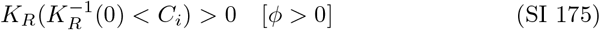

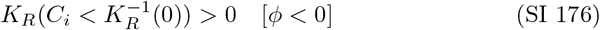

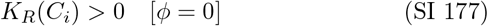
(c) Inequality Relation between 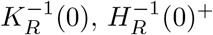,and 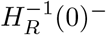 If *ϕ*≠ 0, then 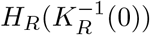 is given by:

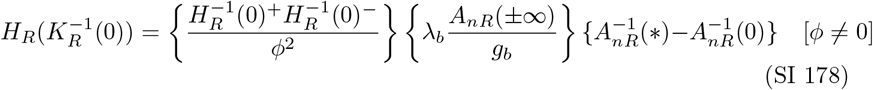 Given that 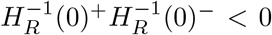 (Eq. (SI 31)), *A*_*nR*_(*±*∞) *>* 0 and 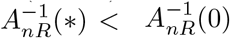 under [C1] (Table SI 1), we derive:

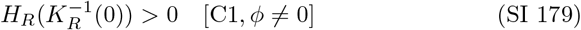 Since *H*_*R*_(*C*_*i*_) is a convex quadratic function with two real roots 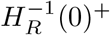 and 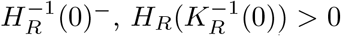 (Eq. (SI 179)) indicates that 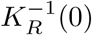 is located outside the region of 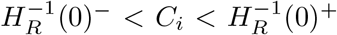.Combining this with 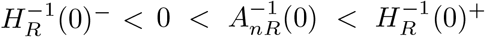 under [C1] (Eq. (SI 46)), we obtain the following inequalities:

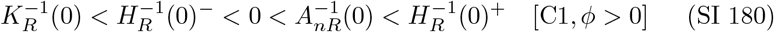

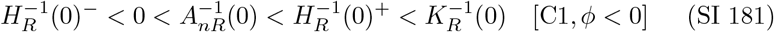 By combining Eqs. (SI 180) and (SI 181) with Eqs. (SI 175) to (SI 177), we obtain:

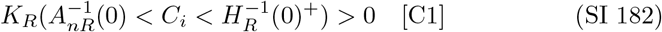
(d) 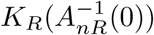 If *ϕ*≠ 0, we have:

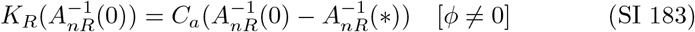 Since 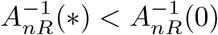 under [C1] (Table SI 1), it follows that:

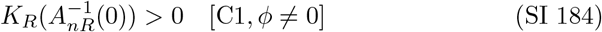 By combining Eq. (SI 184) with Eq. (SI 177), we obtain:

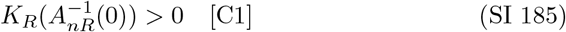
(e) Geometry of *K*_*R*_(*C*_*i*_) Figures SI 3 to SI 5 and Tables SI 26 to SI 28 illustrate the curves and sign charts of *K*_*R*_(*C*_*i*_).

**Fig. SI 3:**
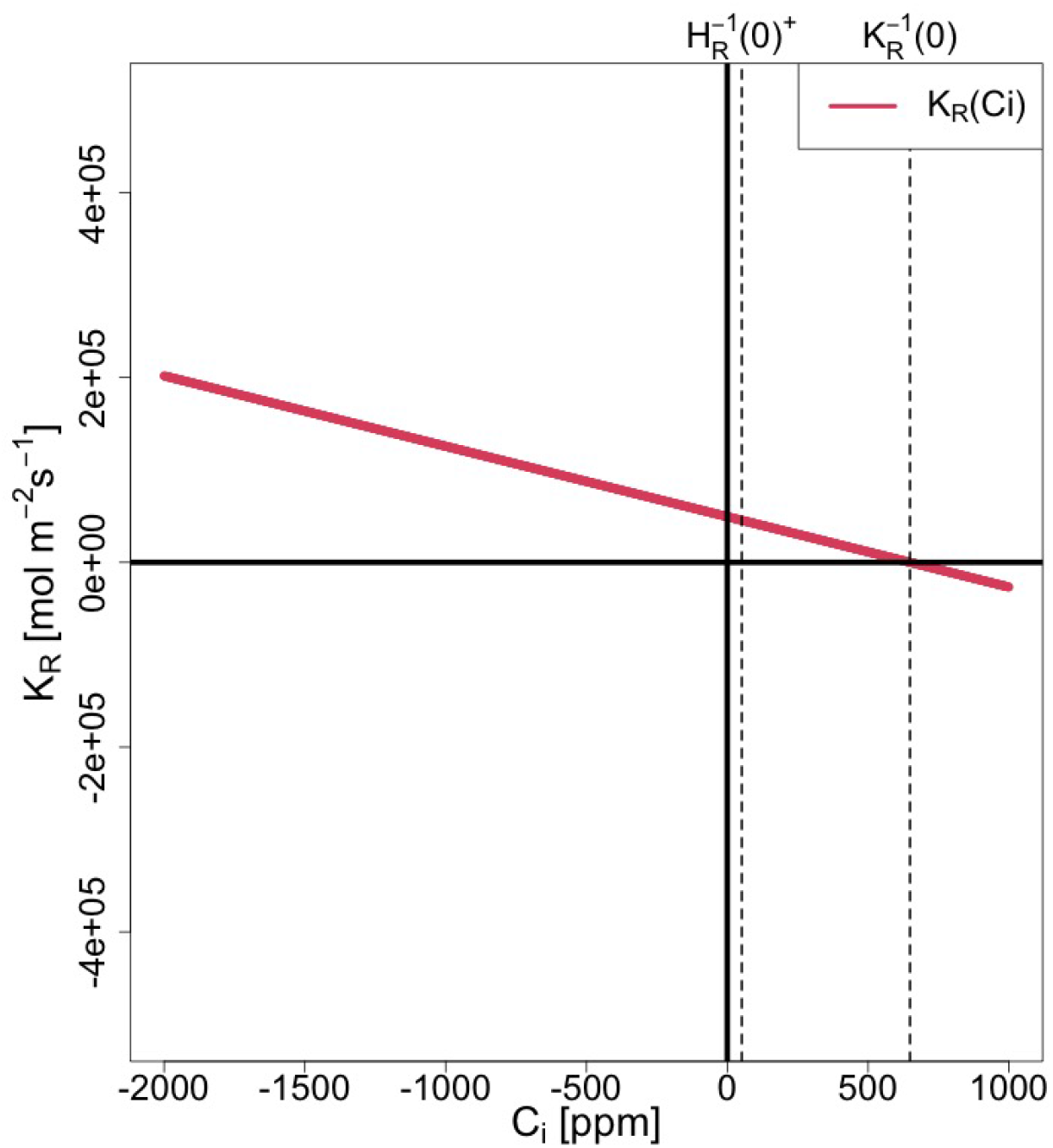
*K*_*R*_(*C*_*i*_) for [C1, *ϕ <* 0]

**Fig. SI 4:**
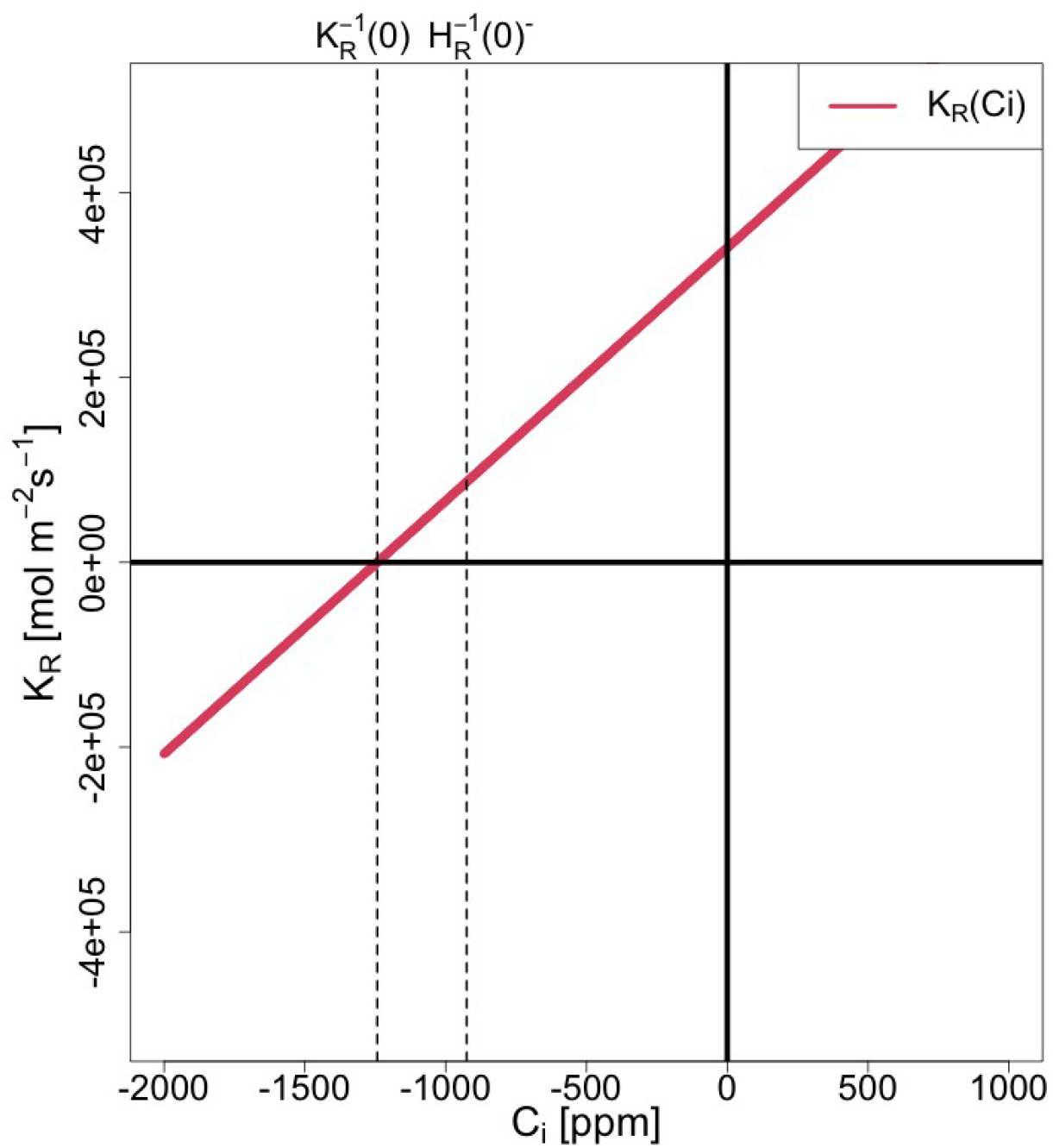
*K*_*R*_(*C*_*i*_) for [C1, *ϕ >* 0]

**Fig. SI 5:**
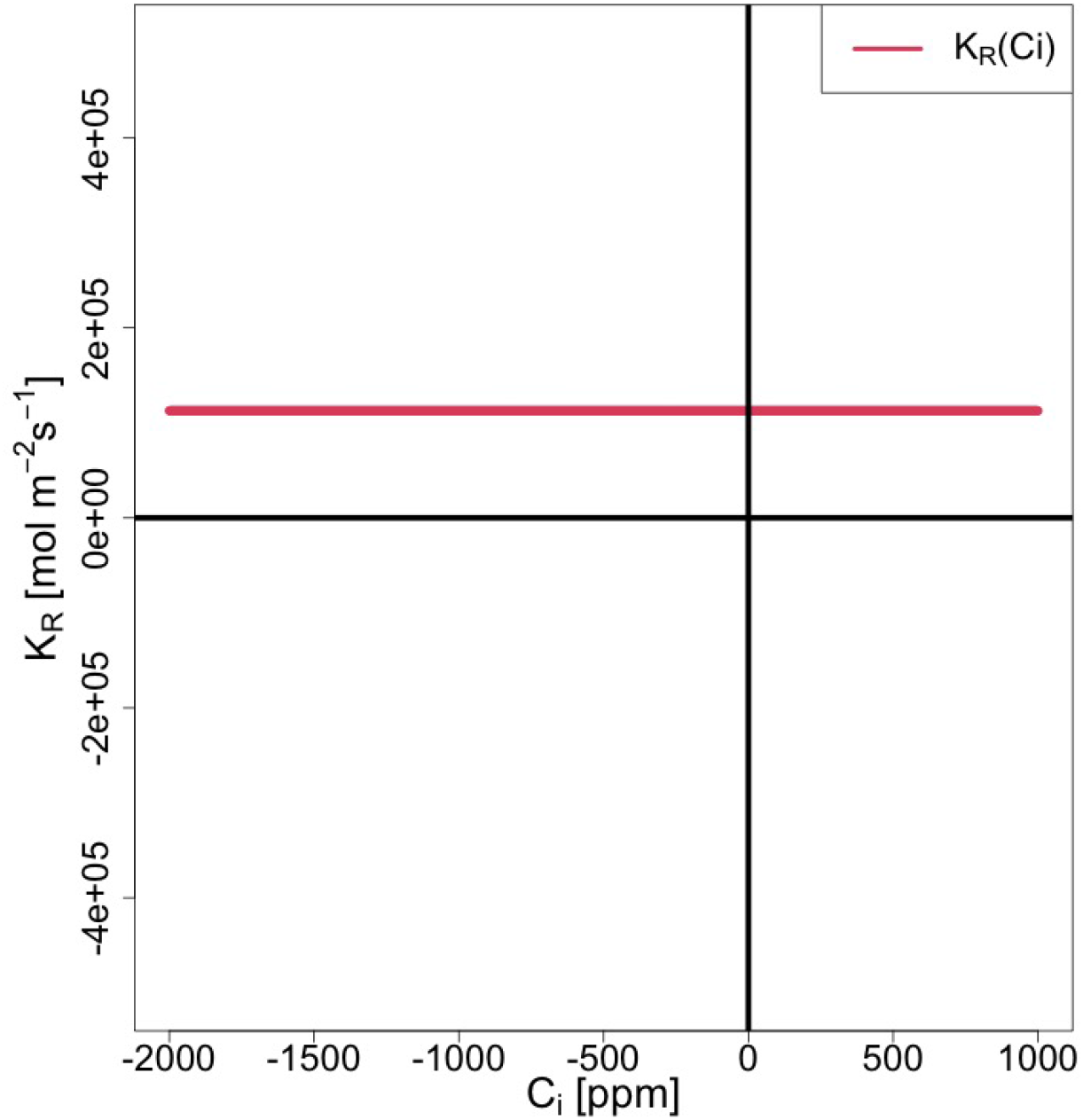
*K*_*R*_(*C*_*i*_) for [C1, *ϕ* = 0]

**Table SI 26:**
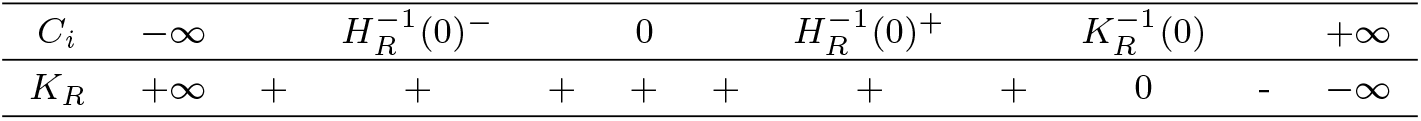
Sign Chart of *K*_*R*_(*C*_*i*_) [C1, *ϕ <* 0].

| $C_i$ | $-\infty$ | | $H_R^{-1}(0)^-$ | | 0 | | $H_R^{-1}(0)^+$ | | $K_R^{-1}(0)$ | | $+\infty$ |
| --- | --- | --- | --- | --- | --- | --- | --- | --- | --- | --- | --- |
| $K_R$ | $+\infty$ | + | + | + | + | + | + | + | 0 | - | $-\infty$ |

**Table SI 27:**
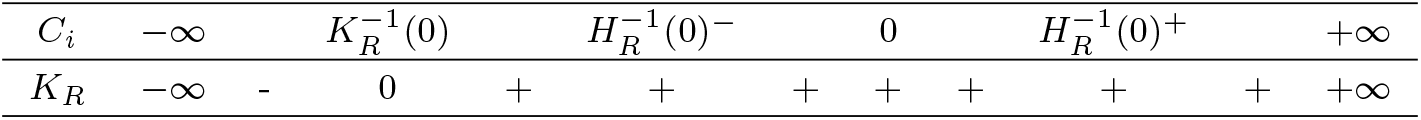
Sign Chart of *K*_*R*_(*C*_*i*_) [C1, *ϕ >* 0].

| $C_i$ | $-\infty$ | | $K_R^{-1}(0)$ | | $H_R^{-1}(0)^-$ | | 0 | | $H_R^{-1}(0)^+$ | | $+\infty$ |
| --- | --- | --- | --- | --- | --- | --- | --- | --- | --- | --- | --- |
| $K_R$ | $-\infty$ | - | 0 | + | + | + | + | + | + | + | $+\infty$ |

**Table SI 28:** Sign Chart of *K*_*R*_(*C*_*i*_) [C1, *ϕ* = 0].

| $C_i$ | $-\infty$ | | $H_R^{-1}(0)^-$ | | 0 | | $H_R^{-1}(0)^+$ | | $+\infty$ |
| --- | --- | --- | --- | --- | --- | --- | --- | --- | --- |
| $K_R$ | + | + | + | + | + | + | + | + | + |

#### SI 5.3 *N*_*R*_(*C*_*i*_) = 0

We explore the roots of *Z*_*R*_(*C*_*i*_) = 0 to examine the geometry of *Z*_*R*_(*C*_*i*_). From Eq. (SI 137), the roots of *Z*_*R*_(*C*_*i*_) = 0 are 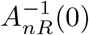 and those of *N*_*R*_(*C*_*i*_) = 0. Thus, we examine *N*_*R*_(*C*_*i*_) = 0. Note that [C1] is assumed for *N*_*R*_(*C*_*i*_) since it is assumed for *Z*_*R*_(*C*_*i*_).

*N*_*R*_(*C*_*i*_) can be expressed as the difference between two functions, *L*_*R*_(*C*_*i*_) and *M*_*R*_(*C*_*i*_), as follows:

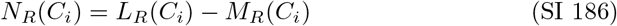

where *L*_*R*_(*C*_*i*_) and *M*_*R*_(*C*_*i*_) are given by

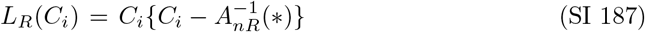

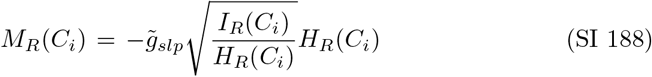

The roots of 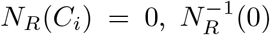,can be found as the values of *C*_*i*_ that satisfy *L*_*R*_(*C*_*i*_) = *M*_*R*_(*C*_*i*_), which indicates the points of intersection between *L*_*R*_(*C*_*i*_) and *M*_*R*_(*C*_*i*_).

#### SI 5.3.1 *L*_*R*_(*C*_*i*_)

Eq. (SI 187) shows that *L*_*R*_(*C*_*i*_) is a convex quadratic function of *C*_*i*_.

(a) Roots: 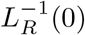 From Eq. (SI 187), 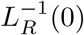 is given by:

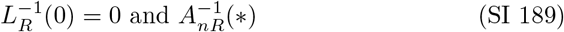
(b) Geometry of *L*_*R*_(*C*_*i*_) Figure SI 6 and Table SI 29 show the curve and sign chart of *L*_*R*_(*C*_*i*_).

**Fig. SI 6:**
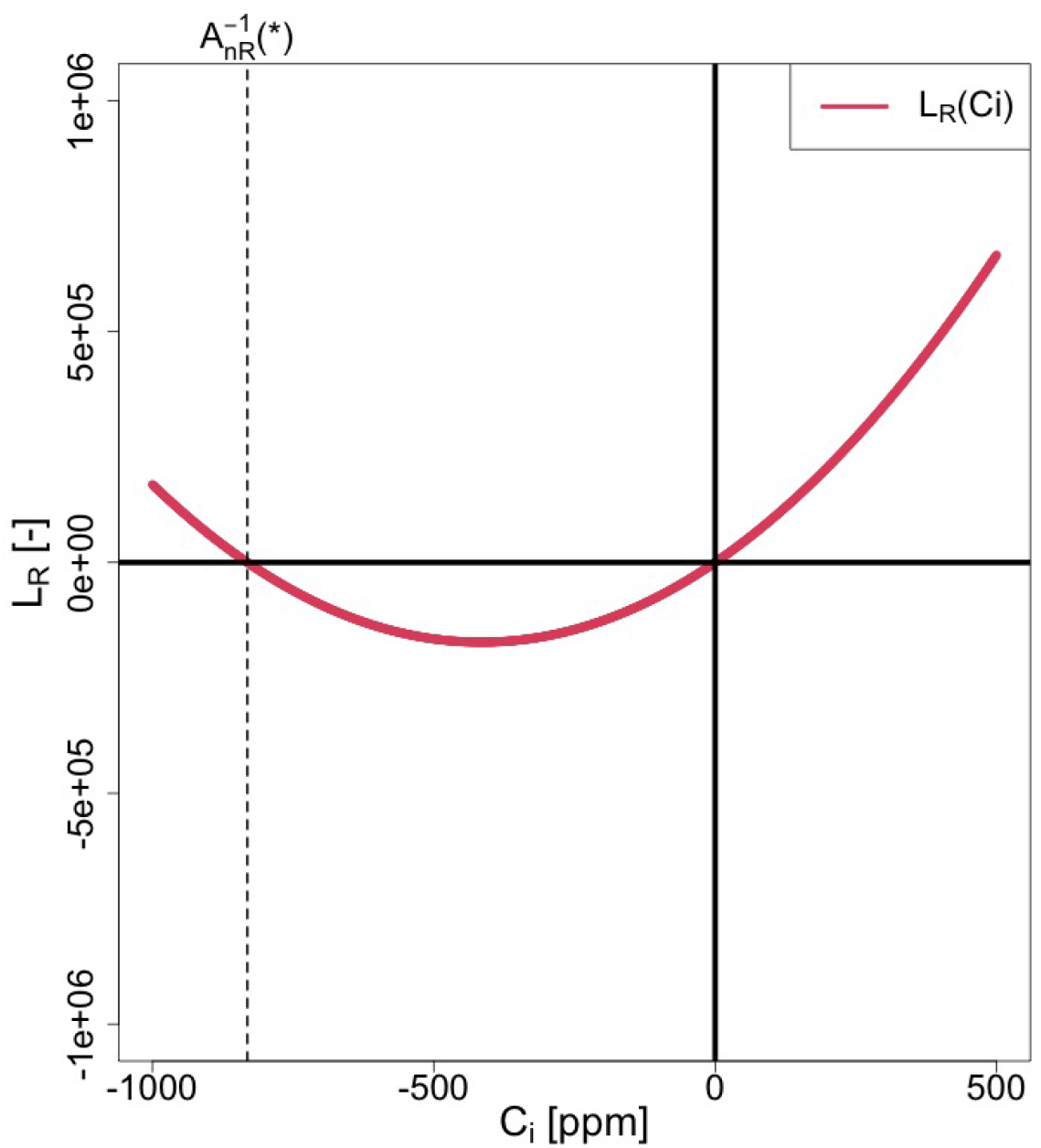
*L*_*R*_(*C*_*i*_)

**Table SI 29:**
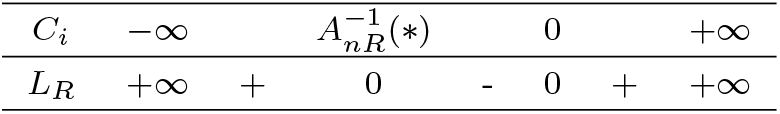
Sign chart of *L*_*R*_(*C*_*i*_)

#### SI 5.3.2 *M*_*R*_(*C*_*i*_)

(a) Roots: 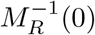 From Eq. (SI 188), 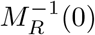 is given by:

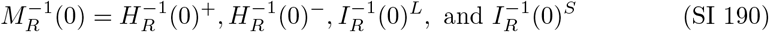
(b) Geometry of *M*_*R*_(*C*_*i*_) Since there are three distinct inequalities for *I*_*R*_(*C*_*i*_) under [C1], depending on the sign of 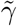,there are also three distinct inequalities for *M*_*R*_(*C*_*i*_). Figure SI 7 illustrates the curve of *M*_*R*_(*C*_*i*_) under the condition 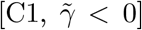. Notably, the curves of *M*_*R*_(*C*_*i*_) under 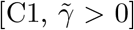 and 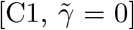 are identical to that under 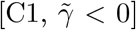, Table SI 30 summarizes the sign chart of *M*_*R*_(*C*_*i*_), focusing on the possible range of appropriate solutions under [C1] (see Table 2).

**Fig. SI 7:**
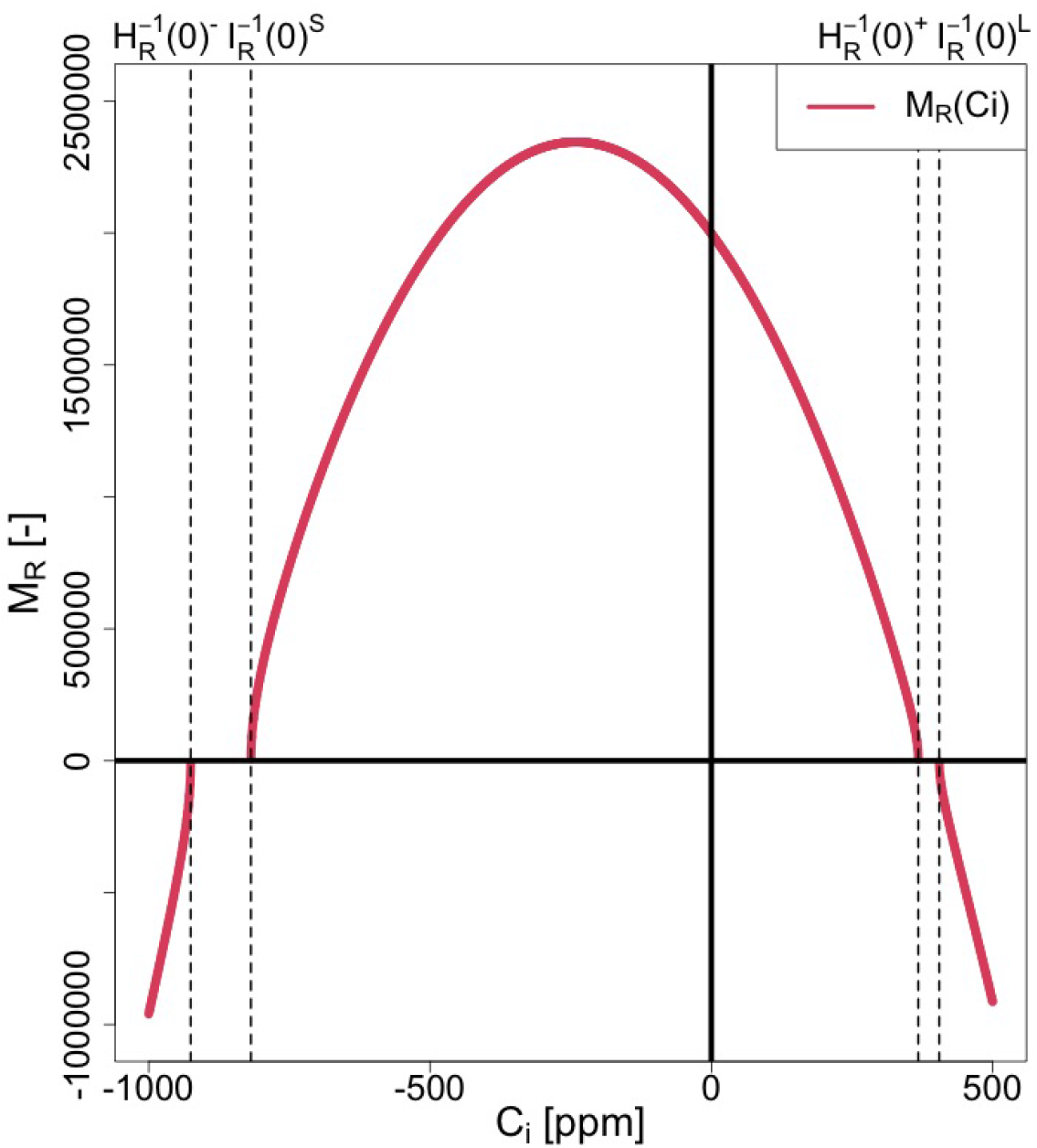
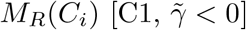

#### SI 5.3.3 Sign of *L*_*R*_(*C*_*i*_) and *M*_*R*_(*C*_*i*_)

By focusing on the interval 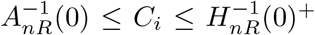, which represents the possible range of solutions under [C1] (as shown in Table 2), we can deduce from Tables SI 29 and SI 30 that both *L*_*R*_(*C*_*i*_) and *M*_*R*_(*C*_*i*_) are positive or zero within this range:

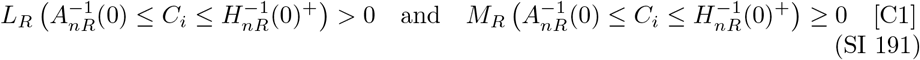

**Table SI 30:**
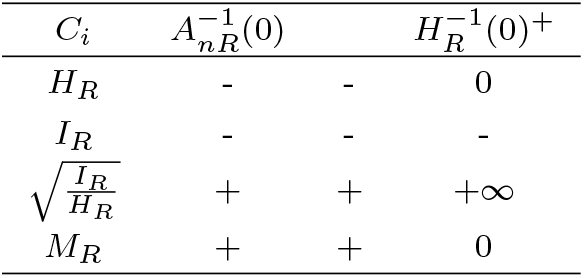
Sign chart of *M*_*R*_(*C*_*i*_) [C1].

### SI 5.4 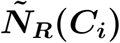

To facilitate analysis of *N*_*R*_(*C*_*i*_), which is challenging due to the square root terms in *M*_*R*_(*C*_*i*_), we introduce a modified function 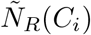 defined as follows:

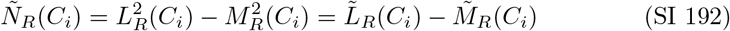

where 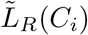 and 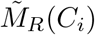 represent the squares of *L*_*R*_(*C*_*i*_) and *M*_*R*_(*C*_*i*_), respectively, and are given by:

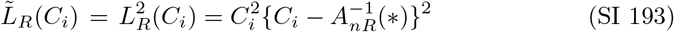

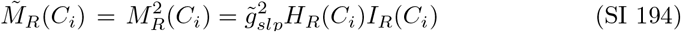

The function 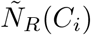 can also be expressed using *N*_*R*_(*C*_*i*_), *L*_*R*_(*C*_*i*_), and *M*_*R*_(*C*_*i*_) as follows:

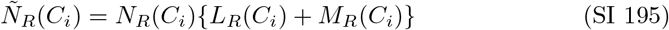

Eq. (SI 195) shows that *N*_*R*_(*C*_*i*_) and 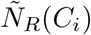 share the same roots and signs at *C*_*i*_ where *L*_*R*_(*C*_*i*_) + *M*_*R*_(*C*_*i*_) *>* 0. Therefore, we have:

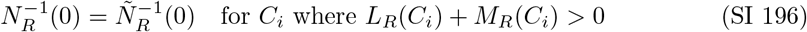

In fact, Eq. (SI 191) indicates that *L*_*R*_(*C*_*i*_) + *M*_*R*_(*C*_*i*_) *>* 0 hold true within the interval 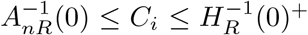 under condition [C1]. Thus, we proceed to analyze 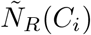 in detail in the following sections.

#### SI 5.4.1 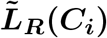

Eq. (SI 193) shows that 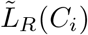 is a quartic function of *C*_*i*_.

(a) Roots: 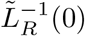 From Eq. (SI 193), 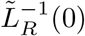 is given by:

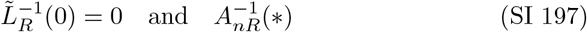
(b) 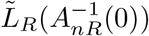 and 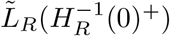

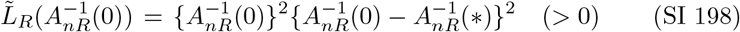

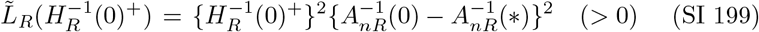
(c) Geometry of 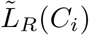 Figure SI 8 and Table SI 31 illustrate the curve and sign chart of 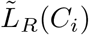.

**Fig. SI 8:**
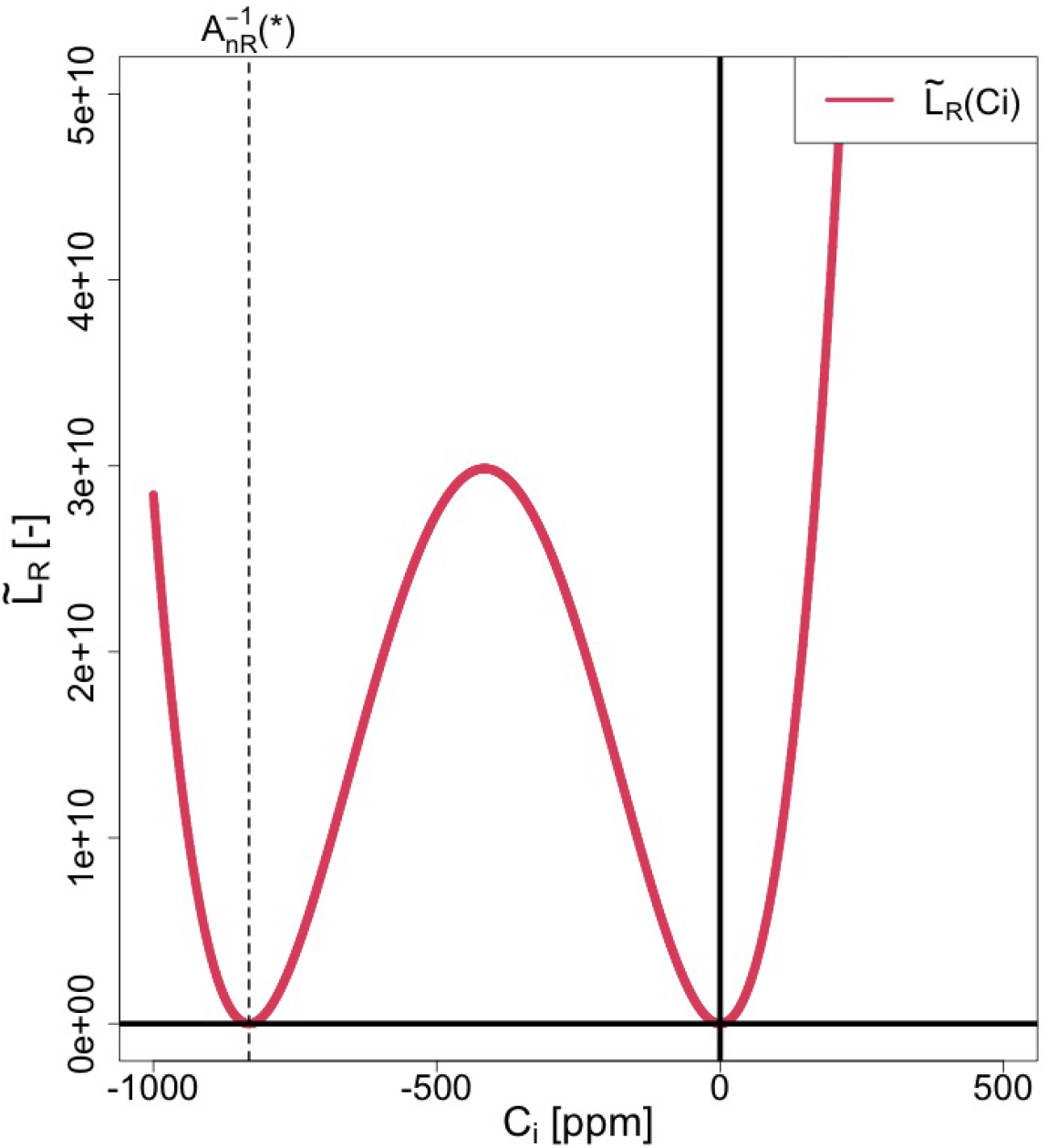
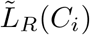

**Table SI 31:** Sign chart of 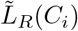.

|  |  |  |  |  |  |  |  |
| --- | --- | --- | --- | --- | --- | --- | --- |
| $C_i$ | $-\infty$ | | $A_{nR}^{-1}(*)$ | | 0 | | $+\infty$ |
| $\tilde{L}_R$ | $+\infty$ | + | 0 | + | 0 | + | $+\infty$ |

#### SI 5.4.2 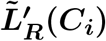

Differentiating Eq. (SI 193) with respect to *C*_*i*_ yields:

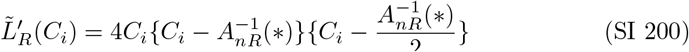

Eq. (SI 200) shows that 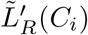 is a cubic function of *C*_*i*_.

(a) Roots: 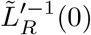 From Eq. (SI 200), 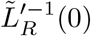 is given by:

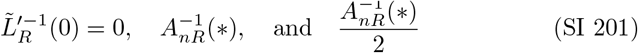 These values satisfy the following inequality since 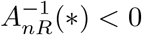 (Table SI 1):

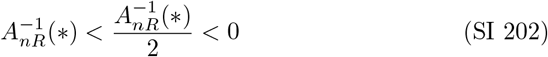 Since 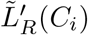 is a cubic function with a positive leading coefficient (Eq. SI 200), we have the following sign changes for 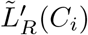:

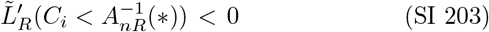

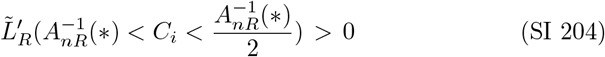

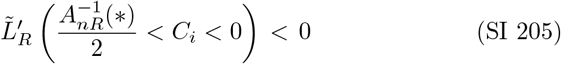

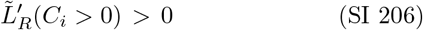
(b) 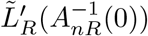

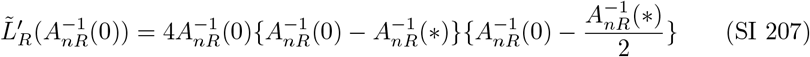
(c) Geometry of 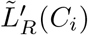

Figure SI 9 and Table SI 32 display the curve and sign chart of 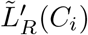.

**Fig. SI 9:**
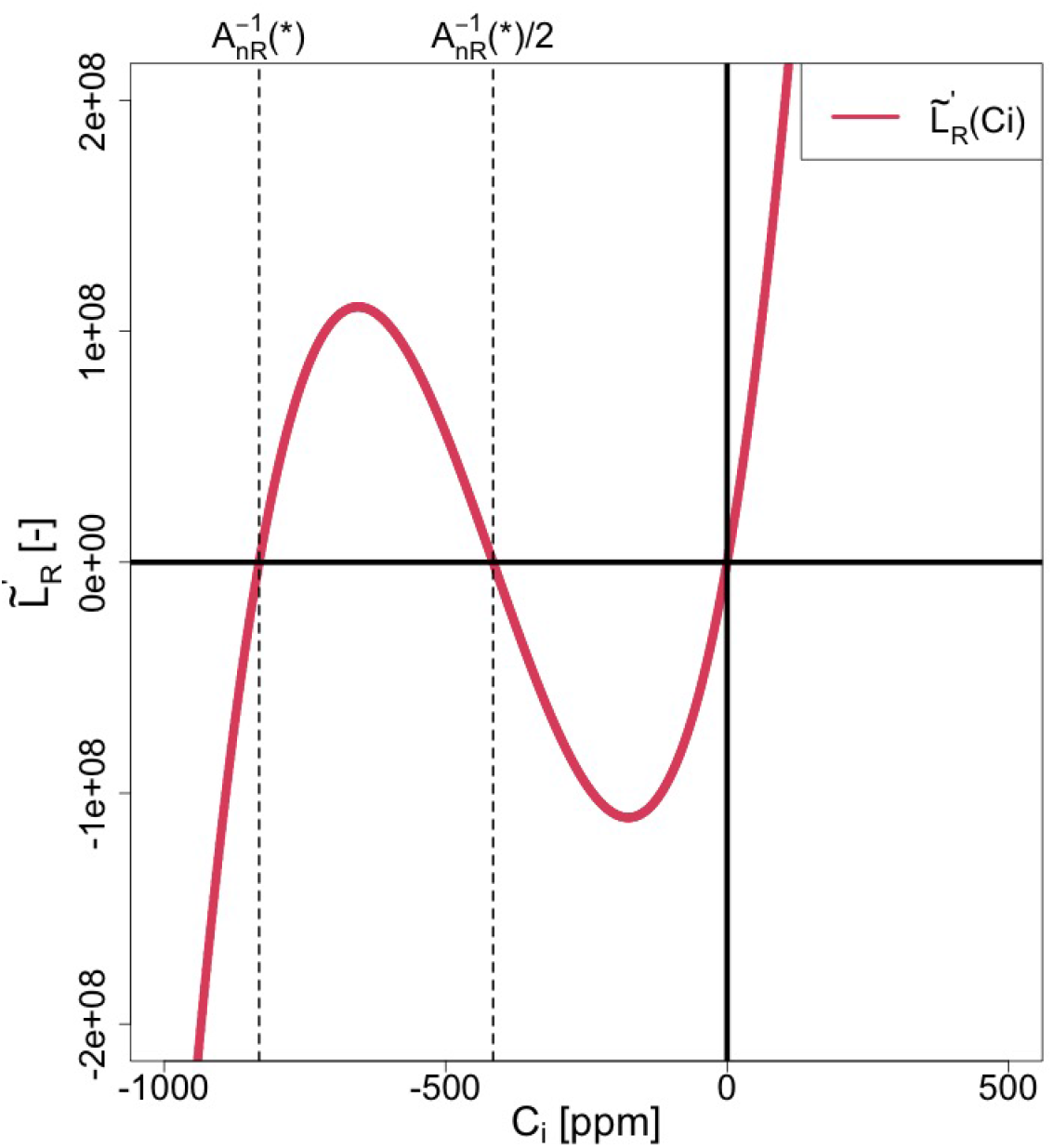
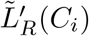

**Table SI 32:**
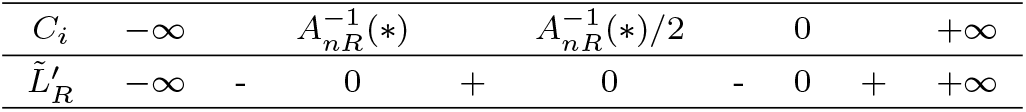
Sign chart of 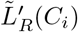.

#### SI 5.4.3 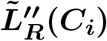

Differentiating Eq. (SI 200) with respect to *C*_*i*_ gives:

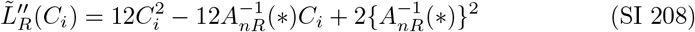

Eq. (SI 208) shows that 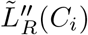 is a convex quadratic function of *C*_*i*_.

(a) Roots: 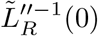 From Eq. (SI 208), 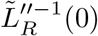 is given by:

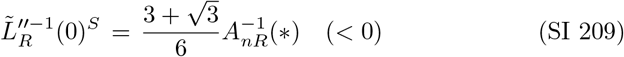

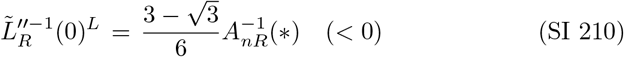 Note that 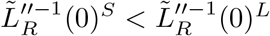 since 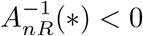 (Table SI 1). Since 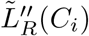 is a convex quadratic function of *C*_*i*_, we have:

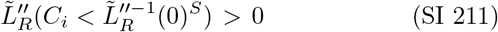

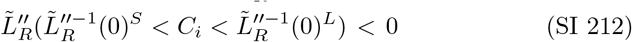

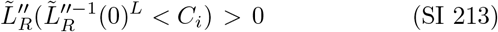 Since 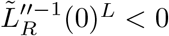 (Eq. (SI 210)), we have in particular:

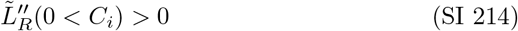
(b) 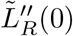

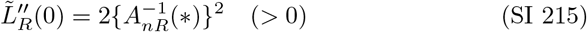
(c) Geometry of 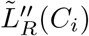 Figure SI 10 and Table SI 33 show the curve and sign chart of 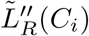.

**Fig. SI 10:**
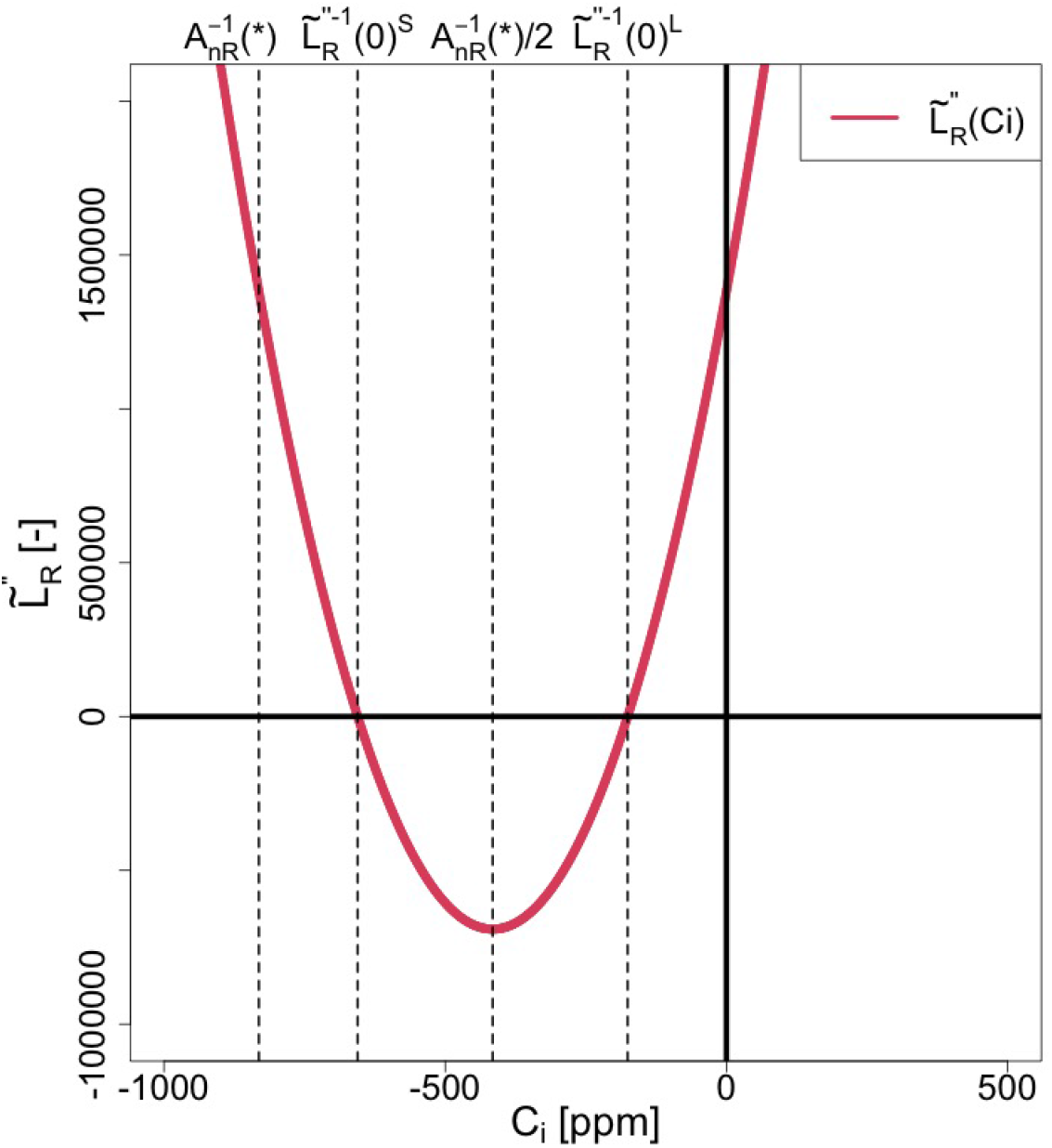
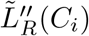

**Table SI 33:**
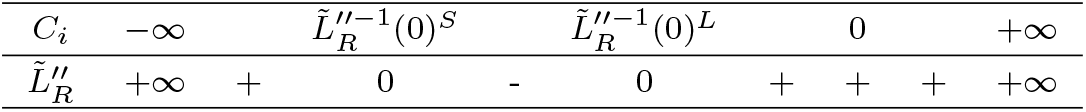
Sign chart of 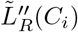.

#### SI 5.4.4 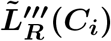

Differentiating Eq. (SI 208) with respect to *C*_*i*_, we obtain:

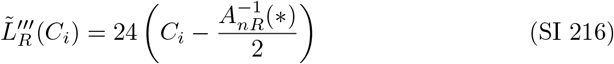

Eq. (SI 216) shows that 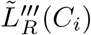 is a linear function of *C*_*i*_ with a positive slope.

(a) Root: 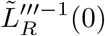 From Eq. (SI 216), 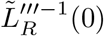 is given by:

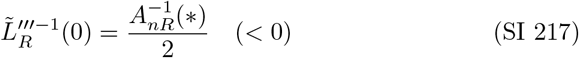 Since the slope of 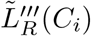 is positive (Eq. (SI 216)), we have the following inequalities:

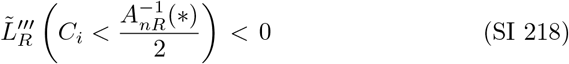

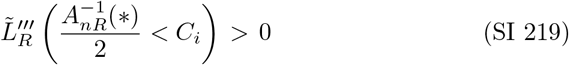
(b) 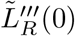

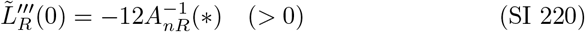
(c) Geometry of 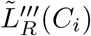 Figure SI 11 and Table SI 34 show the curve and sign chart of 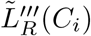.

**Fig. SI 11:**
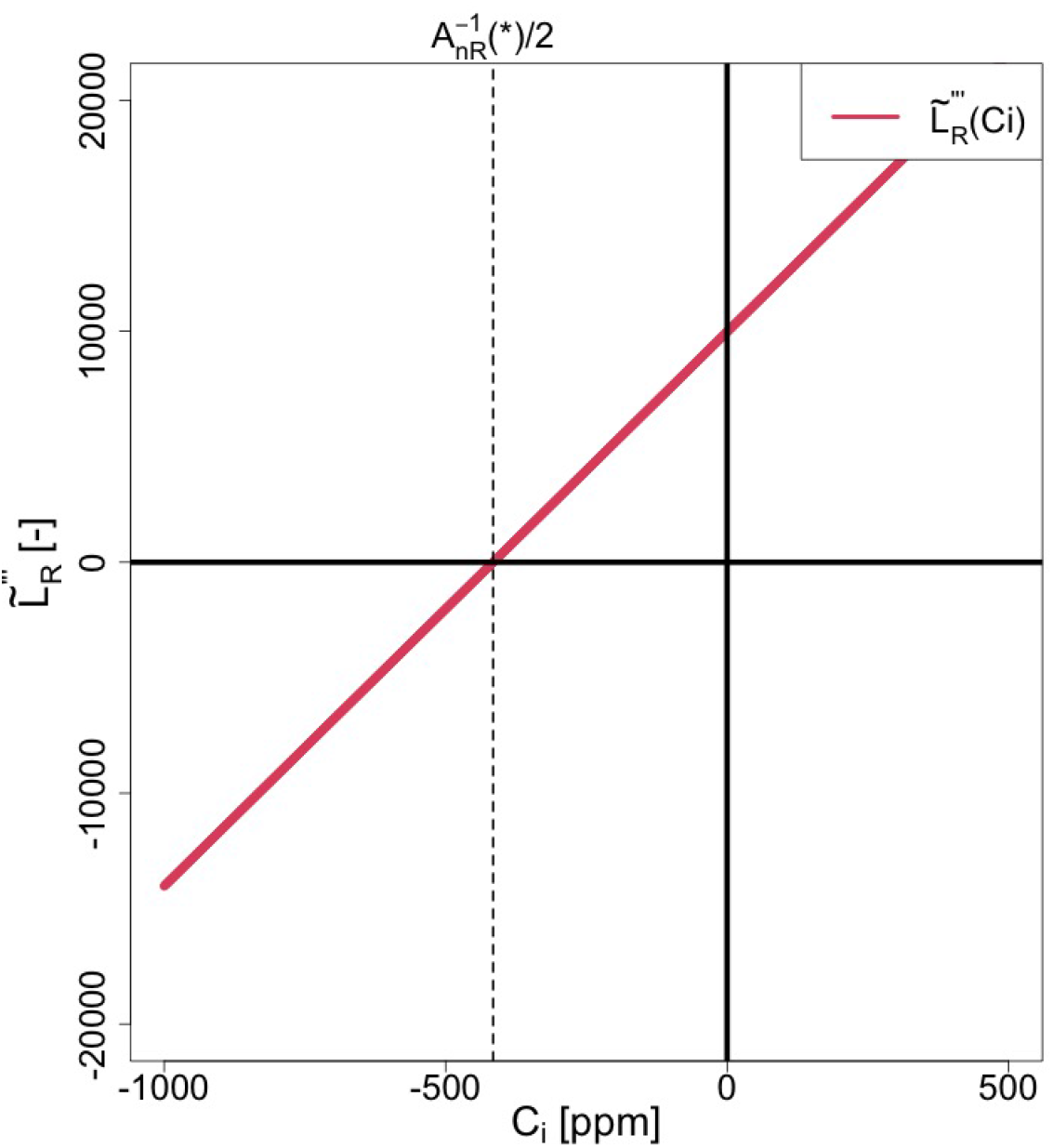
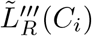

**Table SI 34:** Sign chart of 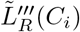.

|  |  |  |  |  |  |  |  |
| --- | --- | --- | --- | --- | --- | --- | --- |
| $C_i$ | $-\infty$ | | $\tilde{L}_R'''^{-1}(0)$ | | $0$ | | $+\infty$ |
| $\tilde{L}_R'''$ | $-\infty$ | - | $0$ | + | + | + | $+\infty$ |

#### SI 5.4.5 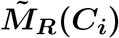

Using Eqs. (SI 29) and (SI 150) for *H*_*R*_(*C*_*i*_) and *I*_*R*_(*C*_*i*_), we can rewrite 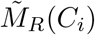 from Eq. (SI 194) as follows:

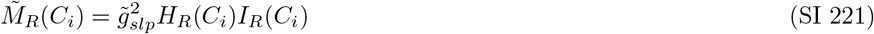

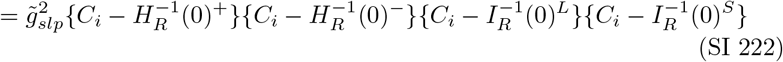

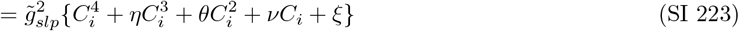

where

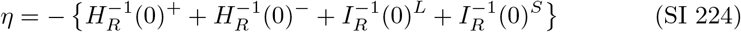

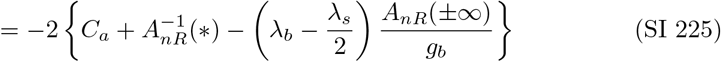

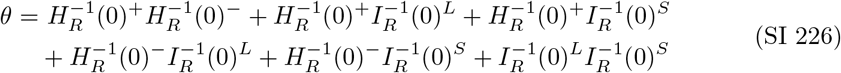

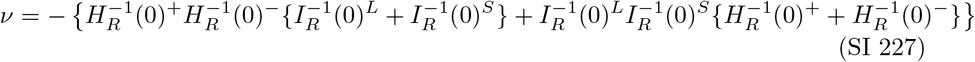

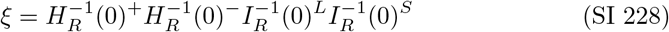

Eq. (SI 223) shows that 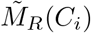 is a quartic function of *C*_*i*_.

(c) Root: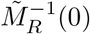 From Eq. (SI 222), 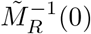 is given by:

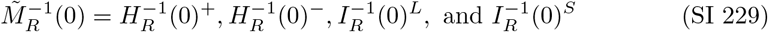
(d) Sings of 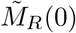 and 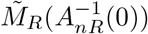 is given by:

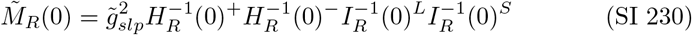 Since 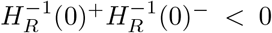 (Eq. (SI 31)) and 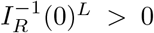 under [C1] (Eq. (SI 149)), we have:

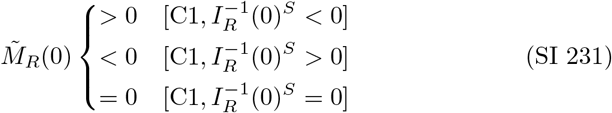 Eq. (SI 231) indicates that the signs of 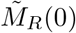 are opposite to those of 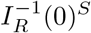 under [C1]. Note that the signs of 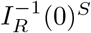 are the same as those of 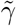 under [C1] (Eq. (SI 153)). Therefore, the signs of 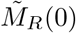 are also opposite to those of 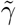 under [C1]. Thus, we have the following relationship under [C1]:

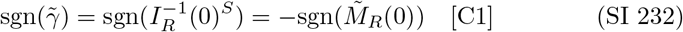 Using Eqs. (SI 37) and (SI 142), we have:

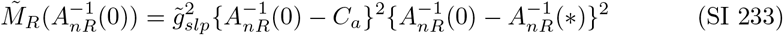
(e) Geometry of 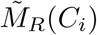 Figures SI 12 to SI 14 and Tables SI 35 to SI 37 show the curves and sign charts of 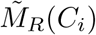.

**Fig. SI 12:**
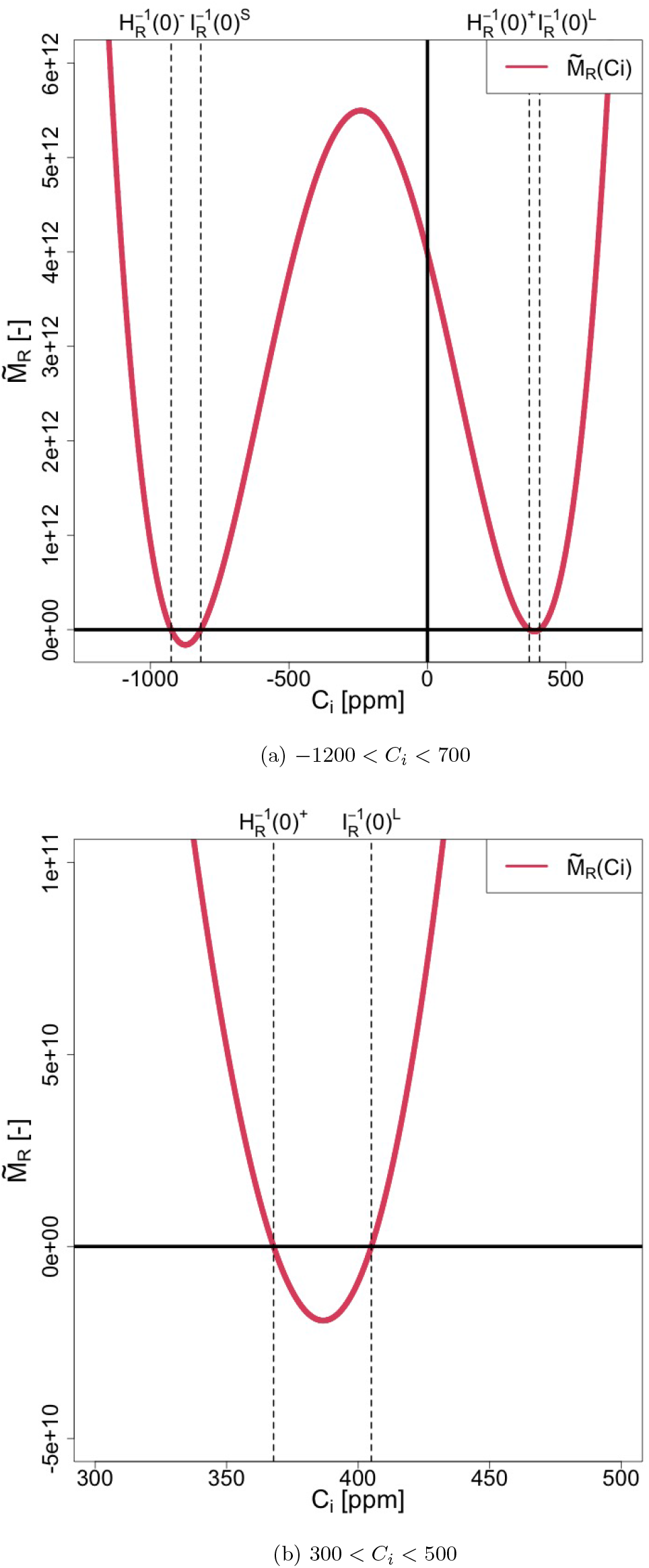
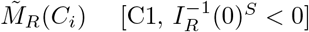

**Fig. SI 13:**
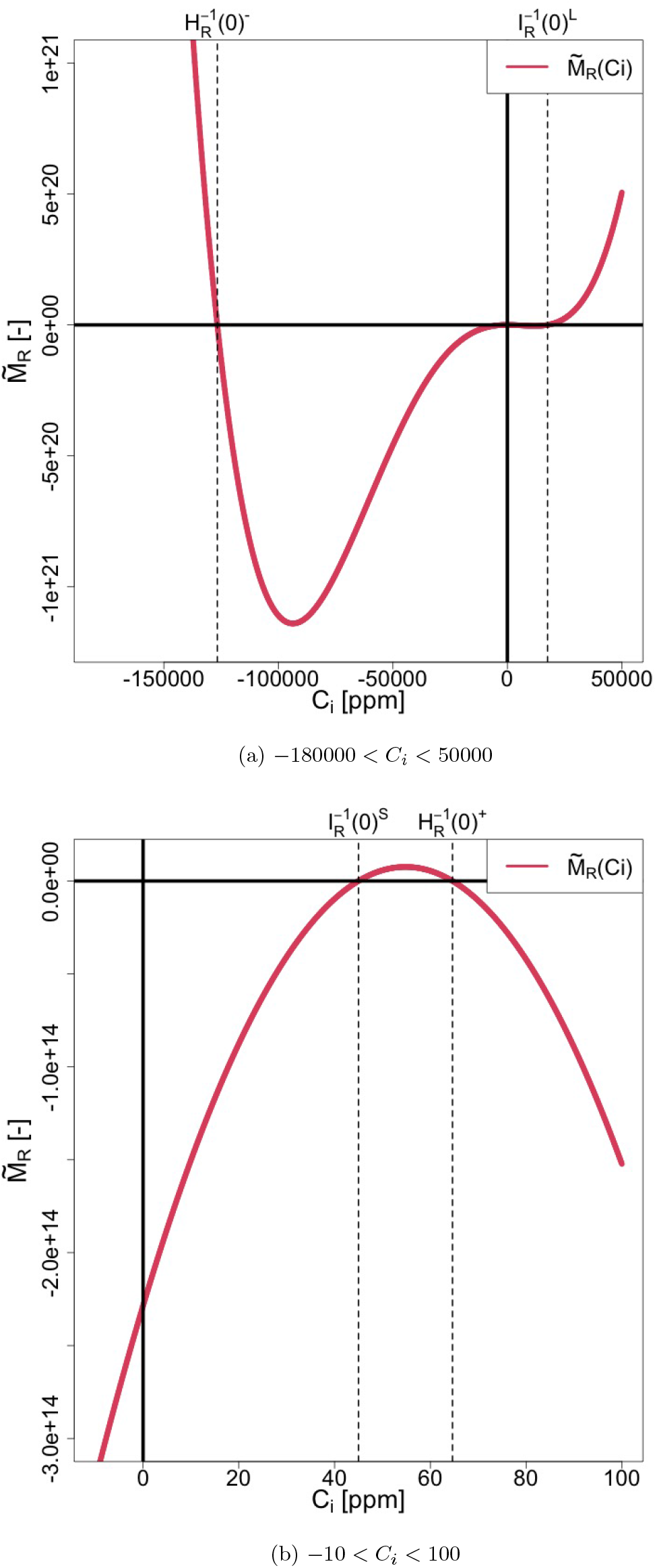
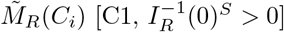

**Fig. SI 14:**
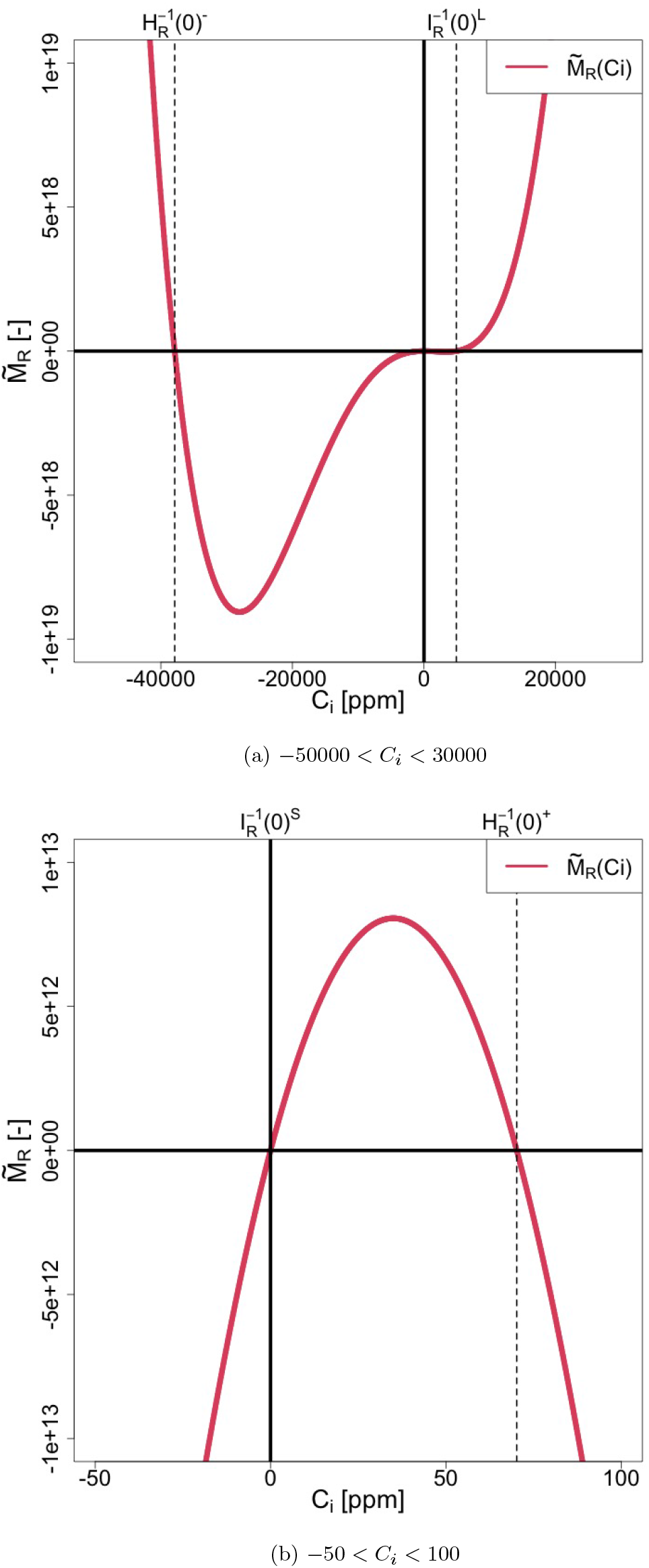
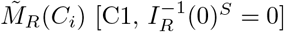.

**Table SI 35:**
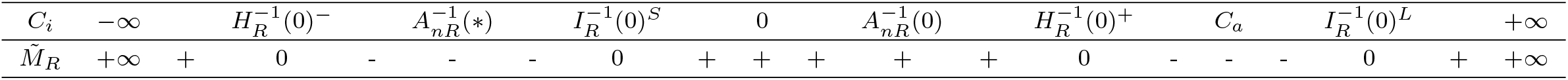
Sign chart of 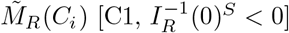.

**Table SI 36:**
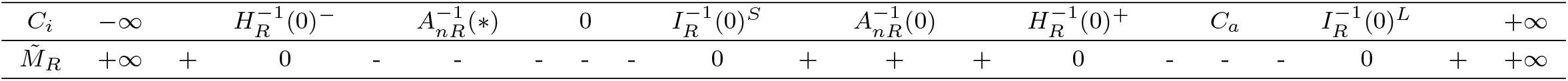
Sign chart of 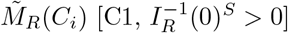.

**Table SI 37:**
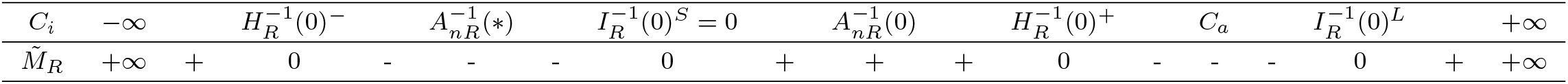
Sign chart of 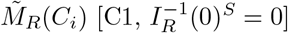.

#### SI 5.4.6 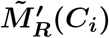

By differentiating Eq. (SI 221) or (SI 223) with respect to *C*_*i*_, we have:

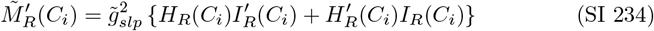

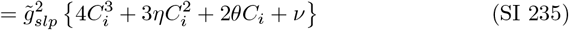

Eq. (SI 235) shows that 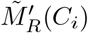 is a cubic function of *C*_*i*_.

(a) 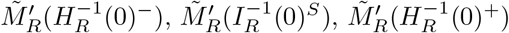,and 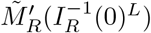 From Eq. (SI 234), we have:

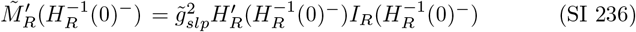

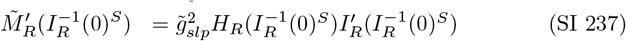

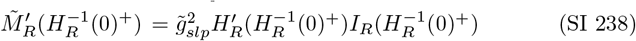

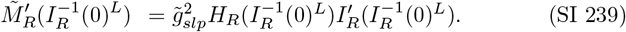 The signs of Eqs. (SI 236) to (SI 239) under [C1] are given as:

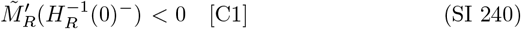

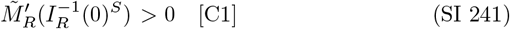

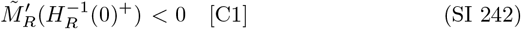

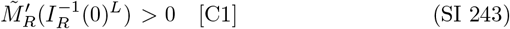 These signs are derived using Eqs. (SI 32) to (SI 34), (SI 79), (SI 80), (SI 155) to (SI 157), (SI 159), (SI 160), and (SI 167) to (SI 169) under [C1].
(b) Roots: 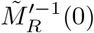 Since 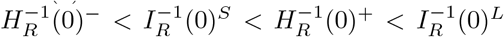 under [C1] (Eqs. (SI 167) to (SI 169)), Eqs. (SI 240) to (SI 243) indicate that there are three roots for 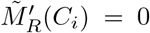 at 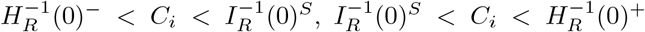,and 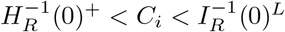.To distinguish them, these are expressed as:

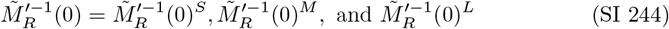

which follow the inequality:

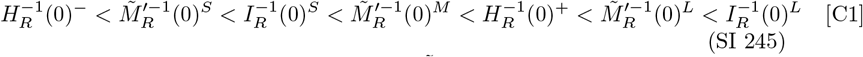 Considering that the coefficient of 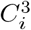 for 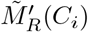 is positive (Eq. (SI 235)), we have:

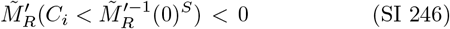

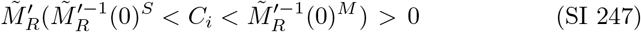

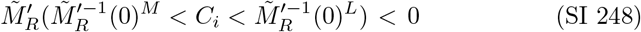

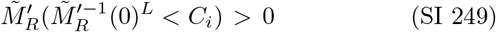
(c) 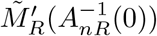

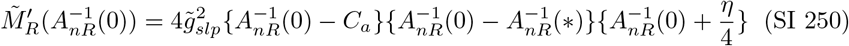 Since 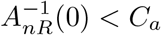 and 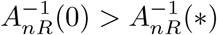 under [C1] (Table SI 1), we have:

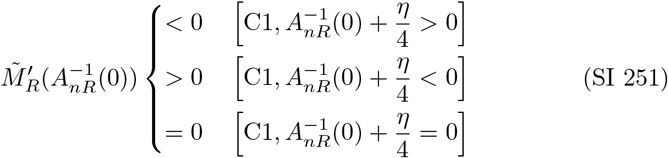 Eq. (SI 251) shows that the signs of 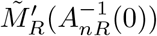 are opposite to those of 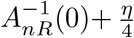 under [C1]. Thus, we have:

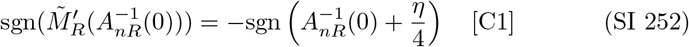
(d) Inequality relation between 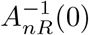 and 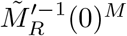 From the signs of 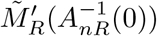,we can derive the inequality relation between 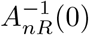 and 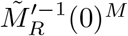,within the range 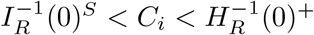 (Eq. (SI 245)). For 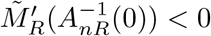, i.e., 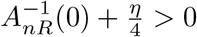 under [C1] (Eq. (SI 252)), we have 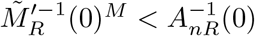 from Eqs. (SI 247) and (SI 248). Similarly, we have:

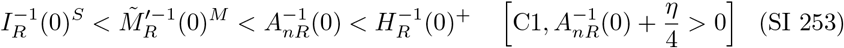

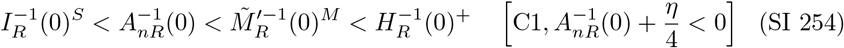

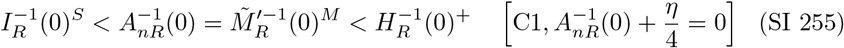
(e) Geometry of 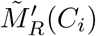 Figures SI 15 to SI 17 show the curves of 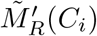,and Tables SI 38 to SI 40 show the sign charts of 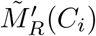 for each inequality pattern from Eqs. (SI 253) to (SI 255) over the interval 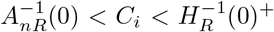,which is the possible range of appropriate solutions under [C1] (Table 2).
(g) Sign of 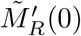(0) From Eq. (SI 235), we have

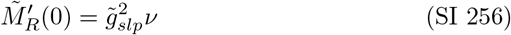 We aim to prove the following proposition regarding the sign of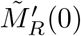.

**Fig. SI 15:**
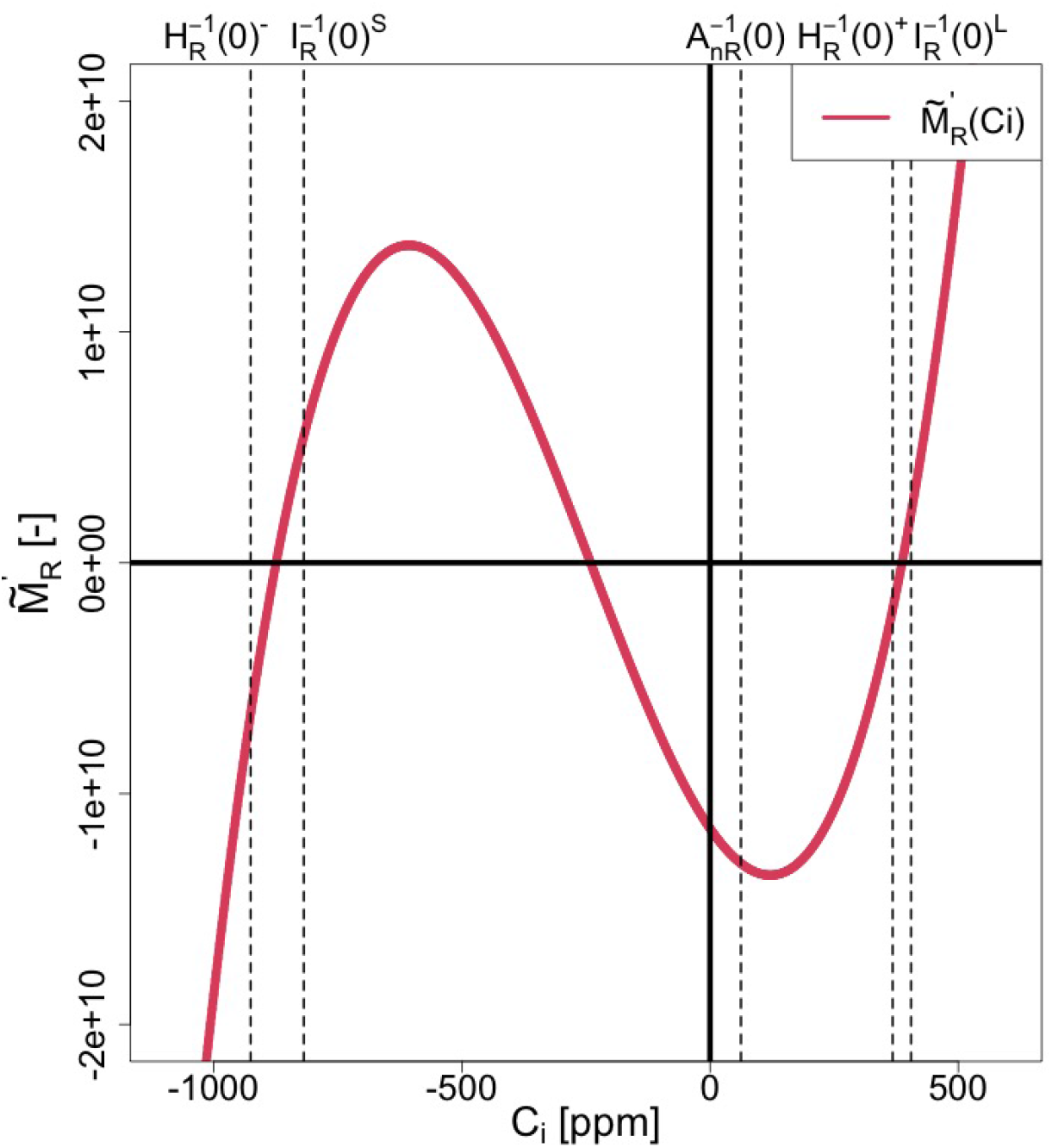
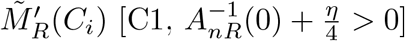

**Fig. SI 16:**
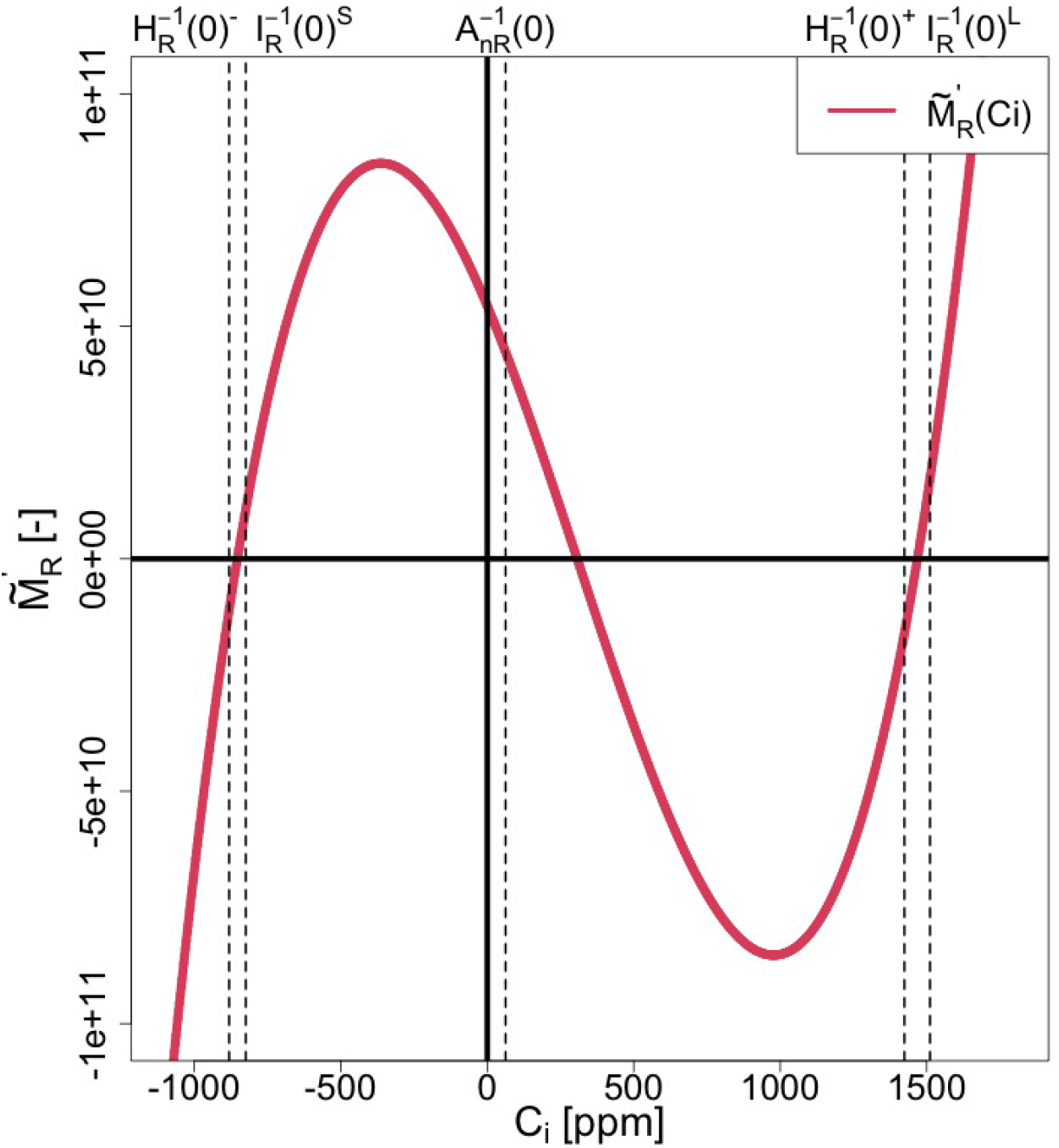
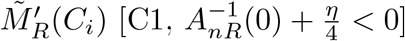

**Fig. SI 17:**
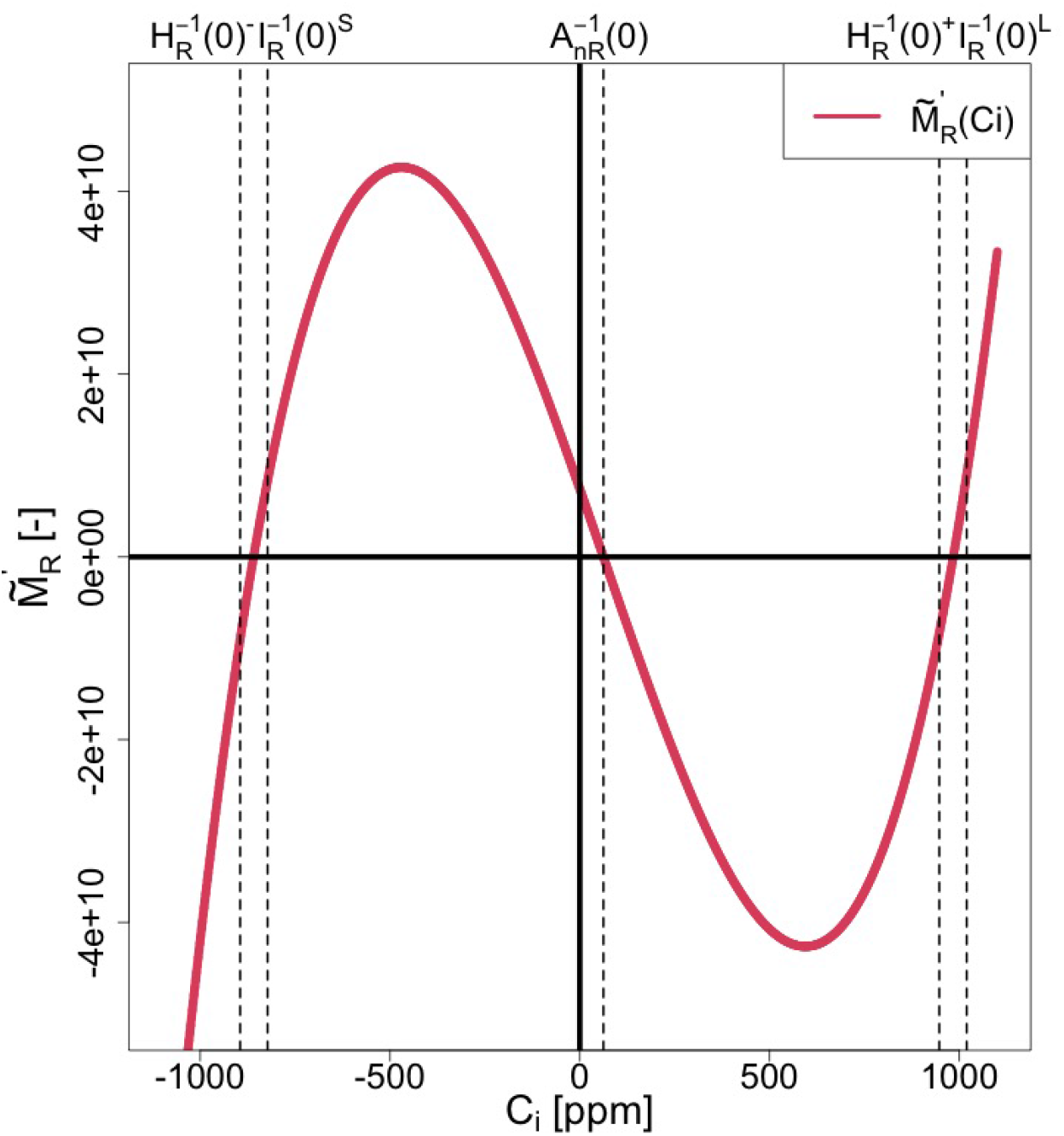
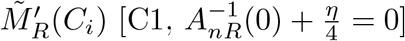

**Table SI 38:**
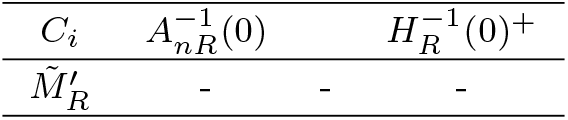
Sign chart of 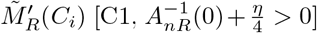.

**Table SI 39:**
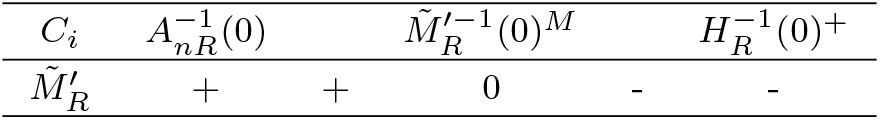
Sign chart of 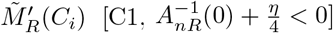.

**Table SI 40:**
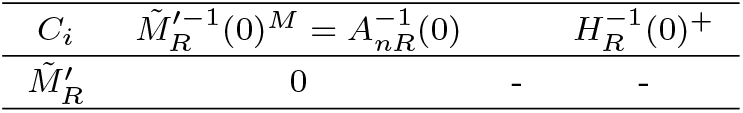
Sign chart of 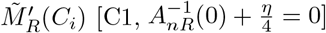.

##### Proposition 1.

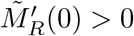*under [C1] and*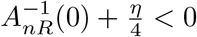. *Proof*. We separate this proof into two cases: 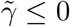and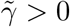.

A. In the case of 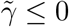 Under 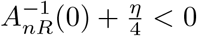and [C1], Eqs. (SI 245) and (SI 254) show

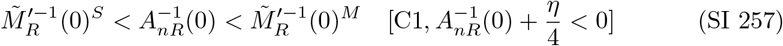 For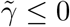, Eq. (SI 232) shows

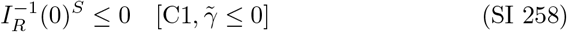 By combining 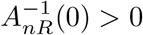 under [C1] (Table SI 1) with (SI 257) and (SI 258), we have

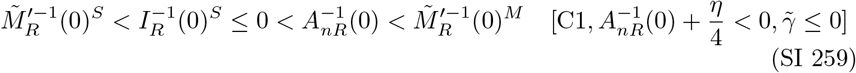 Hence we have

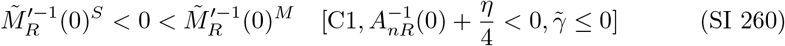 Eqs. (SI 260) and (SI 247) show

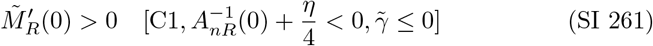
B. In the case of 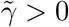 First, we derive an inequality relation for 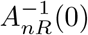 and 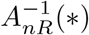 from the condi-tions 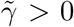 and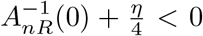. Since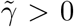, this provides an upper limit for *C*_*a*_, as follows:

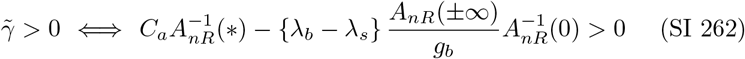

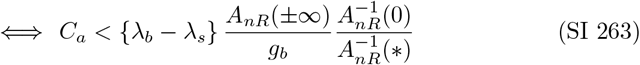 On the other hand, 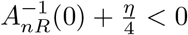 provides a lower limit for *C*_*a*_:

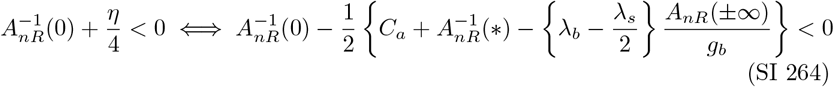

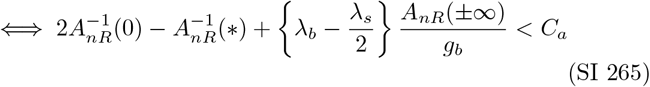 Combining Eqs. (SI 263) and (SI 265), we have

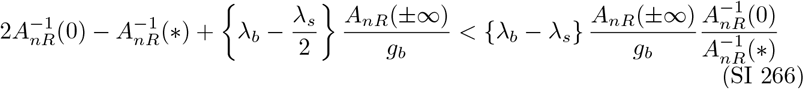

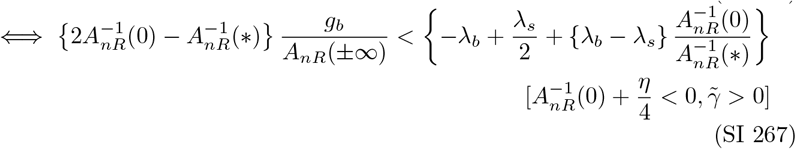

Since the term on the left-hand side of Eq. (SI 267) is positive under [C1] (Table SI 1 and Eq. (SI 8)), the term on the right-hand side of Eq. (SI 267) is also positive, as follows:

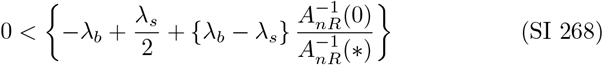

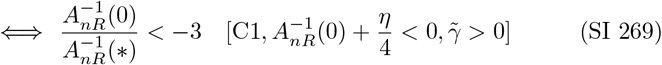

Eq. (SI 269) provides an upper limit for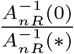.

Next, we examine the sign of ν to determine the sign of 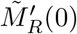(Eq. (SI 256)). From Eq. (SI 227), we can consider ν as a quadratic function with respect to *C*_*a*_, given by

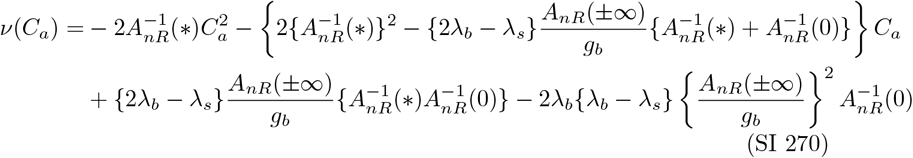

Eq. (SI 270) shows that ν(*C*_*a*_) is a convex quadratic function of *C*_*i*_ since 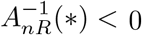(Eq. SI 3).

ν^−1^(min) represents *C*_*a*_, where ν(*C*_*a*_) is minimized, given by

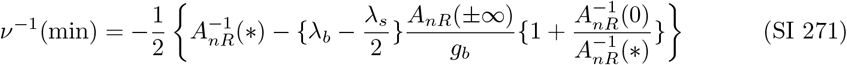

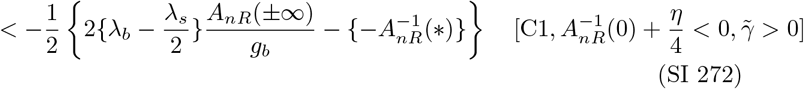

To derive the inequality in Eq. (SI 272), Eq. (SI 269) was used. To explore the sign of ν^−1^(min), we compare the first and second terms in the right-hand side of Eq. (SI 272). From Eq. (SI 263), we have

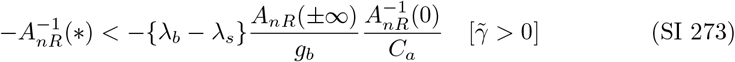

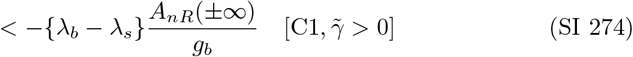

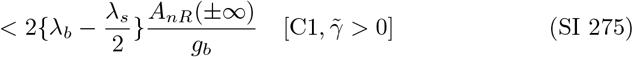

In deriving Eq. (SI 274), we used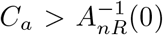, which holds under [C1]. For Eq. (SI 275), we used *λ*_*b*_ = 1.4 and *λ*_*s*_ = 1.6. Eq. (SI 275) shows the inequality relation between the first and second terms on the right-hand side of Eq. (SI 272). Thus we have

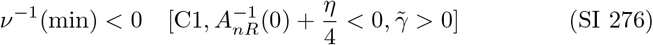

Next, we examine the sign of ν(0). From Eq. (SI 270), ν(0) is given as

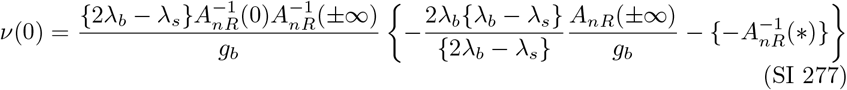

Following similar steps to derive Eq. (SI 275), we have

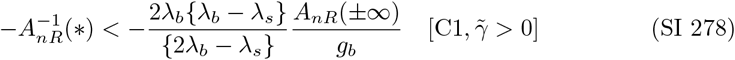

Eq. (SI 278) implies

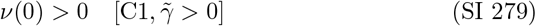

Since ν(*C*_*a*_) is a convex quadratic function of *C*_*a*_ (Eq. (SI 270)), Eqs. (SI 276) and (SI 279) lead to

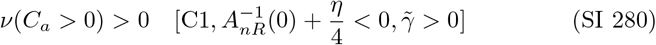

Figure SI 18 shows the curve of ν(*C*_*a*_).

**Fig. SI 18:**
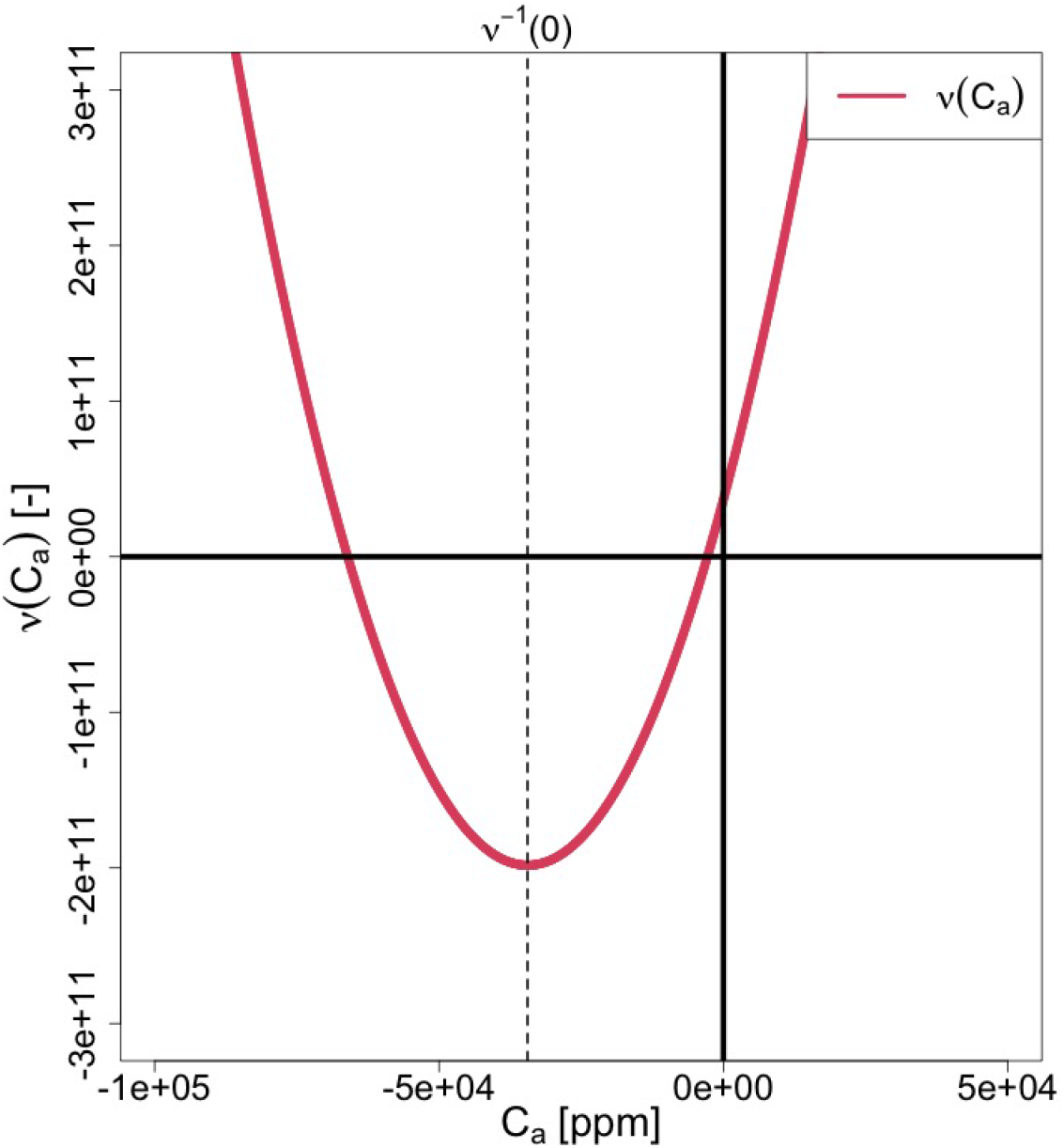
ν (Ca)

Since *C*_*a*_ *>* 0, Eqs. (SI 256) and (SI 280) show

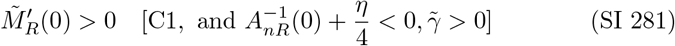

Eqs. (SI 261) and (SI 281) show

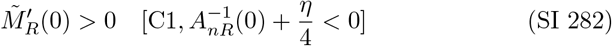

Thus, the proposition is proven.

#### SI 5.4.7 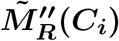

By differentiating Eq. (SI 234) or (SI 235) with respect to *C*_*i*_, we have

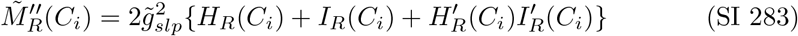

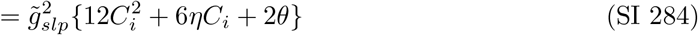

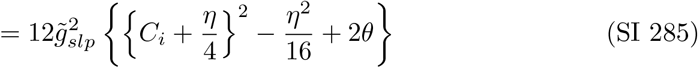

Eq. (SI 285) shows that 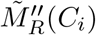 is a convex quadratic function of *C*_*i*_ and has its minimum at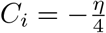, which is expressed as

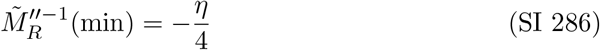

(a) Root: 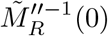 Since there are three roots for 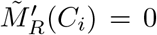(Eq. (SI 244)), there must be two roots for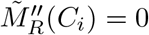. These roots are expressed as

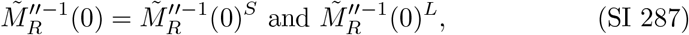

which follow the inequality:

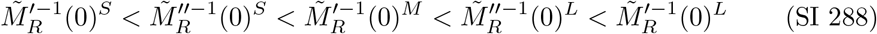 Since 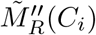 is a convex quadratic function of *C*_*i*_ (Eq. (SI 284)), we have

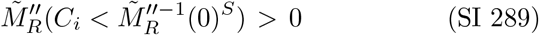

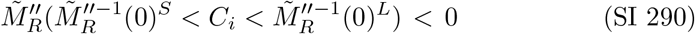

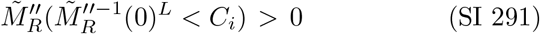 Since 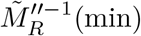 must be in the range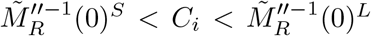, Eq. (SI 286) leads to:

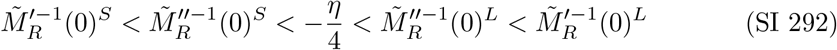
(b) 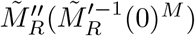 and 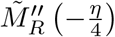 From Eqs. (SI 288), (SI 290), and (SI 292), we have

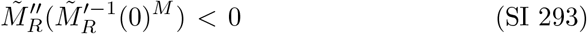

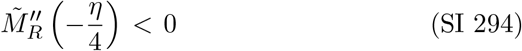
(c) Inequality relationship between 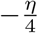 and 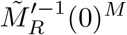 Eqs. (SI 288) and (SI 292) show that 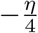 and 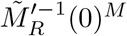 exist within the range 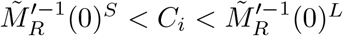, as follows:

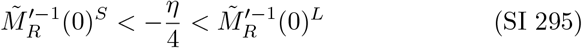

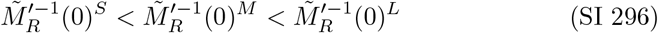 If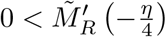, Eq. (SI 247) leads to

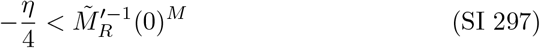 Conversely, if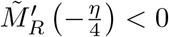, Eq. (SI 248) leads to

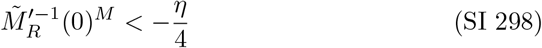 Furthermore, 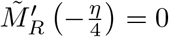 means

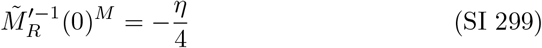 From Eqs. (SI 295) to (SI 299), we have

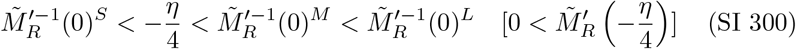

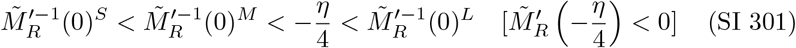

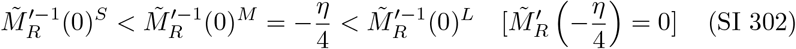

#### SI 5.4.8 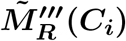

By differentiating Eq. (SI 285) with respect to *C*_*i*_, we have

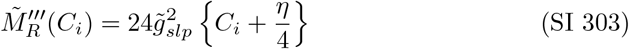

Eq. (SI 303) shows that 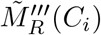 is a linear function of *C*_*i*_ with a positive slope.

(a) Root: 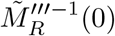 From Eq. (SI 303), 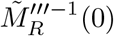is given by

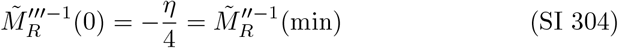 Note that 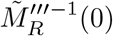 is the inflection point of 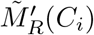and the minimum point of 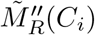. Since the slope of 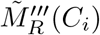 is positive (Eq. (SI 303)), we have

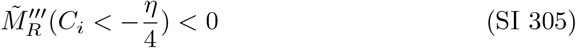

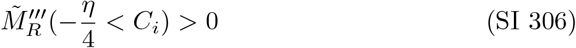

#### SI 5.4.9 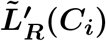 and 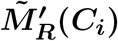

We prove several propositions for 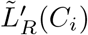 and 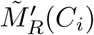.

##### Proposition 2.

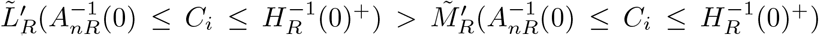 *under [C1] and* 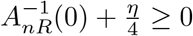.

*Proof*. Since 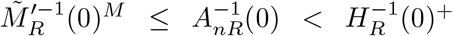 under the conditions [C1] and 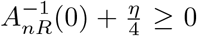 (Eqs. (SI 253) and (SI 255)), and 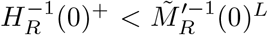 under [C1]

(Eq. (SI 245)), we obtain

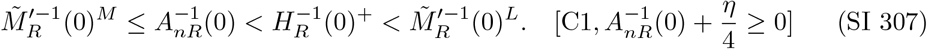

Combining Eq. (SI 307) with 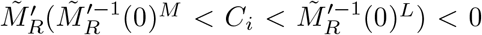 (Eq. (SI 248)) and 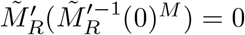 (Eq. (SI 244)), we obtain

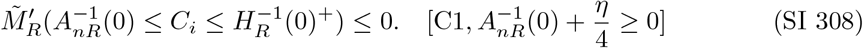

Since 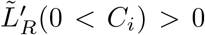 (Eq. (SI 206)) and 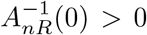 under [C1] (Table SI 1), we obtain

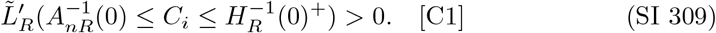

By combining Eqs. (SI 308) and (SI 309), we have

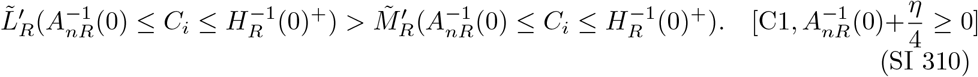

##### Proposition 3.

*There is no intersection between* 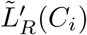 *and* 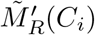 *for* 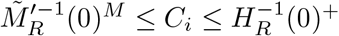 *under [C1] and* 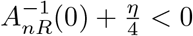.

*Proof*. Eq. (SI 245) leads to

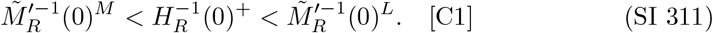

Under [C1] and 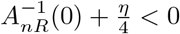, Eqs. (SI 254) and (SI 311) leads to

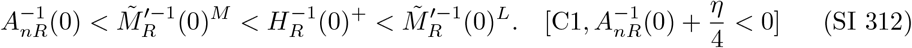

By combining 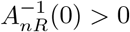 under [C1] (Table SI 1) with Eq. (SI 312), we have

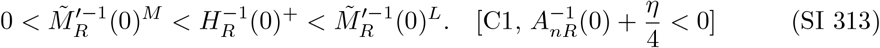

Since 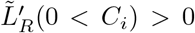 (Eq. (SI 206)) and 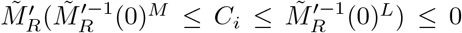 (Eqs. (SI 244) and (SI 248)), considering Eq. (SI 313), we have

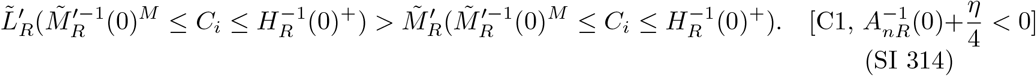

Therefore, there is no intersection between 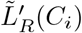 and 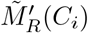 for 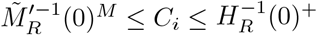 under [C1] and 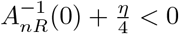.

##### Proposition 4.

*There is only one intersection between* 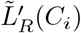 *and* 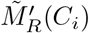 *at* 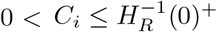 *under C1 and* 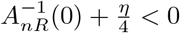.

*Proof*. Proposition 3 shows that there is no intersection between 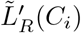 and 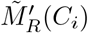 in the range 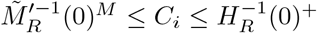 under [C1] and 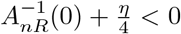. Therefore, we only need to examine the range 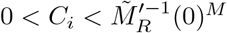. Before that, several inequality relations are derived.

First, 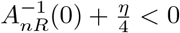, along with 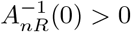 under [C1] (Table SI 1), leads to:

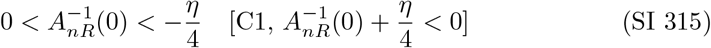

Second, Proposition 1 shows 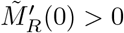 under [C1] and 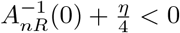. Meanwhile, Eq. (SI 201) shows 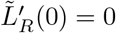. Thus we have:

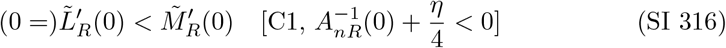

Third, Eq. (SI 219) shows 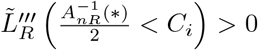. Since 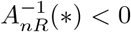 (Eq. (SI 3)), we have

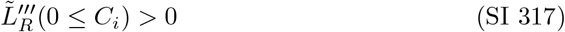

Eqs. (SI 304) and (SI 305) show 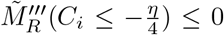. Since 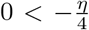 under [C1] and 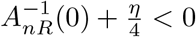 (Eq. (SI 315)), we have

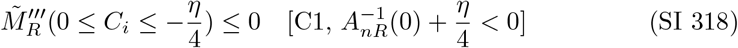

By combining Eqs. (SI 317) and (SI 318), we obtain:

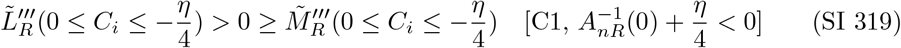

Fourth, since 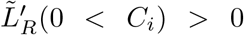 (Eq. (SI 206)) and 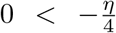 under [C1] and 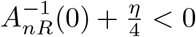 (Eq. (SI 315)), we have:

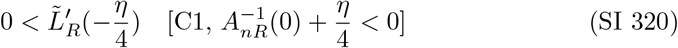

Based on Eq. (SI 320), we split the analysis into three cases: (A): 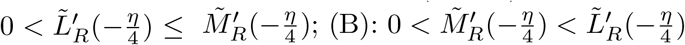 and (C): 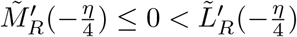.

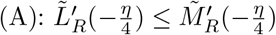

By combining 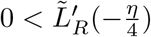 (Eq. (SI 320)) with 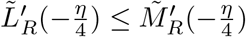, we have

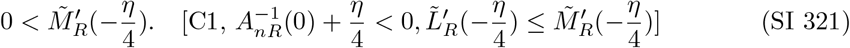

In this case, Eq. (SI 300) shows

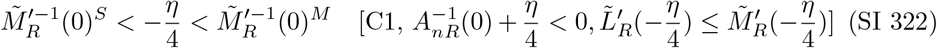

Combining Eq. (SI 322) with 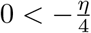 under [C1] and 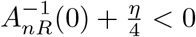 (Eq. (SI 315)), we obtain

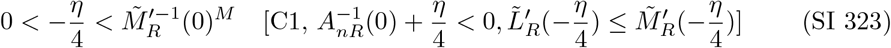

Thus, we divide the range 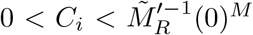 from Eq. (SI 323) into the following two ranges: (i) 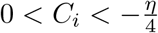 and (ii) 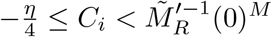.

i. 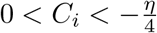 From Eqs. (SI 316), (SI 319), and 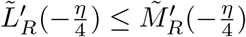, which is the condition of case (A), we have:

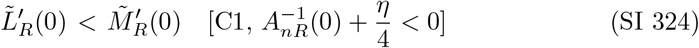

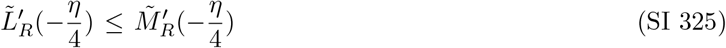

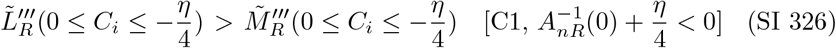 Therefore, there is no intersection between 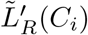 and 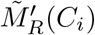 at 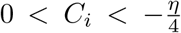 under [C1], 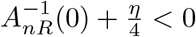, and 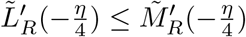.
ii. 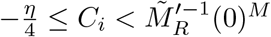 Since 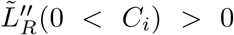 (Eq. (SI 214)) and 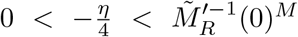 under [C1], 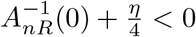, and 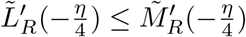 (Eq. (SI 323)), we obtain:

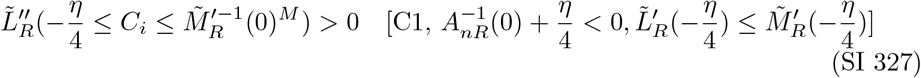 Meanwhile, 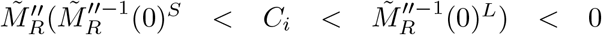 (Eq. (SI 290)) and 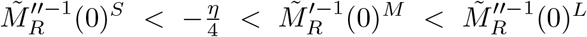 under [C1], 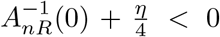, and 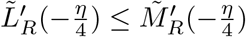 (Eqs. (SI 288), (SI 292), and SI 323) lead to:

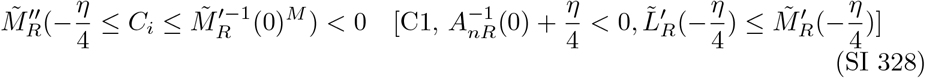

Eqs. (SI 327) and (SI 328) show that

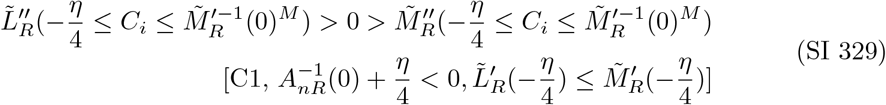

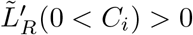 (Eq. (SI 206)) and 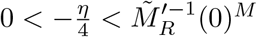 under [C1], 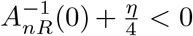, and 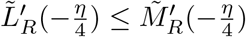 (Eq. (SI 323)) lead to:

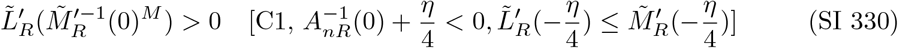

From 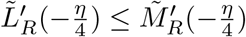, Eqs. (SI 330) and (SI 329), we have:

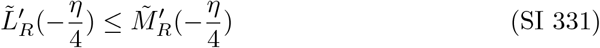

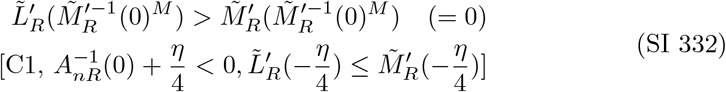

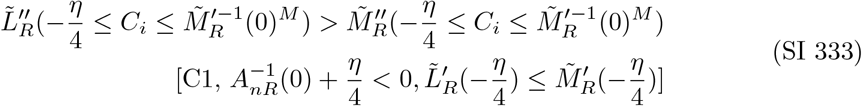

Therefore, there is one intersection between 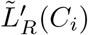 and 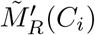 in the range 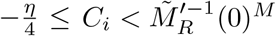 under [C1], 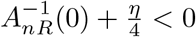, and 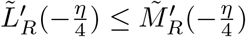.

The results in cases (i) and (ii), along with Proposition 3, show that there are no intersections between 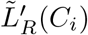 and 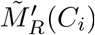 at 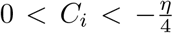, one intersection at 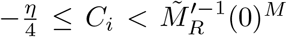, and no intersections at 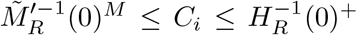 under [C1], 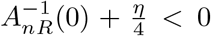, and 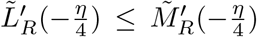. Thus, there is only one intersection between 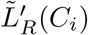 and 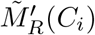 in the range 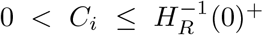 under [C1], 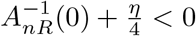, and 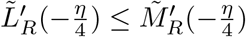.

(B) 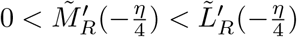 In this case, we have:

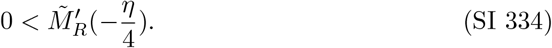 By following the same steps to derive Eq. (SI 323) in case (A), we obtain the same inequality for case (B):

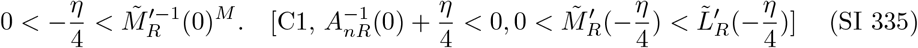 Thus, we separate the range into two parts as follows: (i) 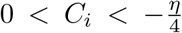 and (ii) 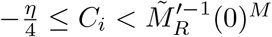
  i. 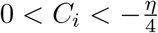 From Eqs. (SI 316) and (SI 319), along with the condition 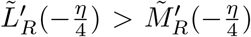, we obtain:

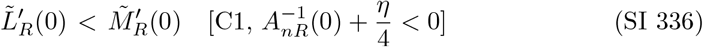

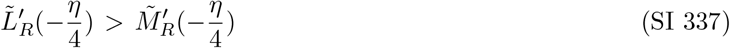

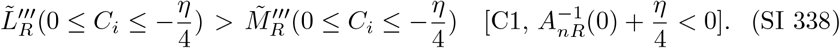 These confirm that there is exactly one intersection between 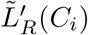 and 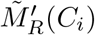 in the range 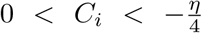 under the conditions [C1], 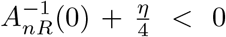, and 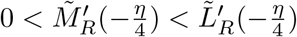.
  ii. 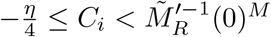 Since 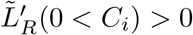 (Eq. (SI 206)) and 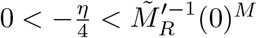 (Eq. (SI 335)), we find:

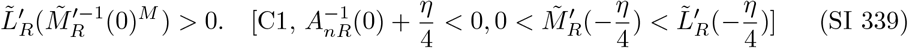 Following the same process as in (A) (Eqs. (SI 327) to (SI 329)), we find:

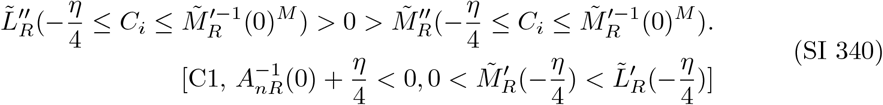 From the conditions of (B), Eqs. (SI 339) and (SI 340), we find:

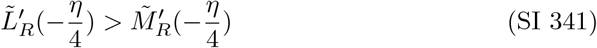

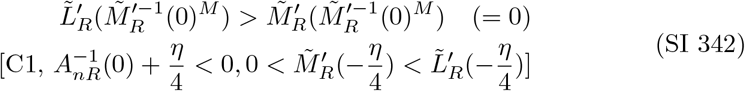

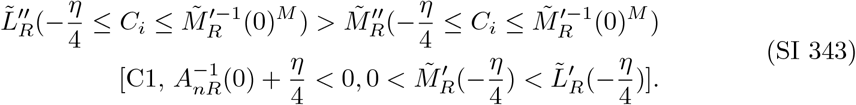 These indicate that there is no intersection between 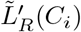 and 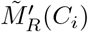 in the range 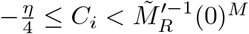 under [C1], 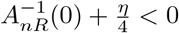, and 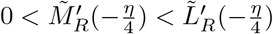. Thus, under the conditions [C1], 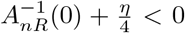, and 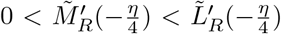, there is only one intersection between 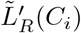 and 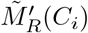 in the range 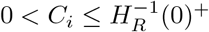.
(C) 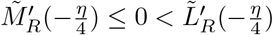 In this case, Eqs. (SI 301) and (SI 302) show that 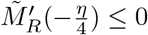 implies:

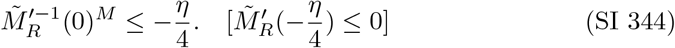 Under 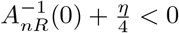, Eq. (SI 254) shows:

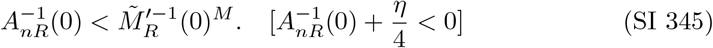

By combining Eqs. (SI 344) and (SI 345) with 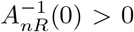 under [C1] (Table SI 1), we obtain:

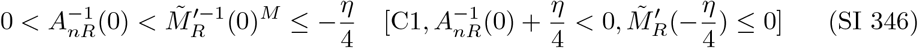

From 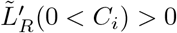 (Eq. (SI 206)) and Eq. (SI 346), we have:

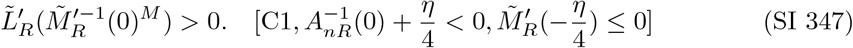

For the range 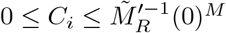, from Eqs. (SI 316), (SI 347), and (SI 319), we find:

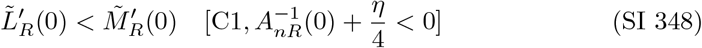

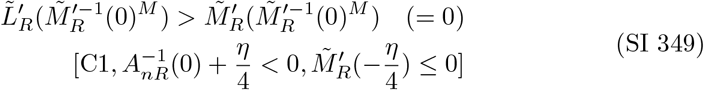

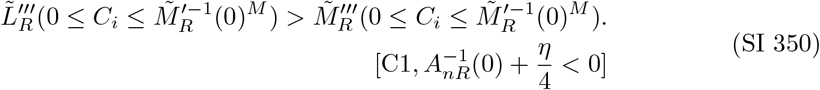

These show that there is only one intersection between 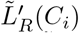 and 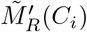 in the range 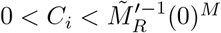 under [C1], 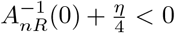, and 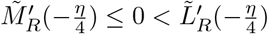.

Proposition 3 further confirms that there are no intersections in the range 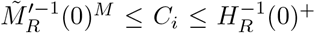 under [C1] and 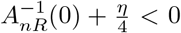.Thus, there is exactly one intersection between 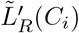 and 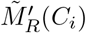 in the range 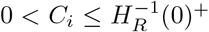 under [C1] and 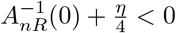,and 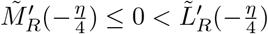.

This completes the proof that there is exactly one intersection between 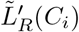 and 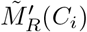 in the range 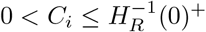 under [C1] and 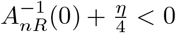.

##### Proposition 5.

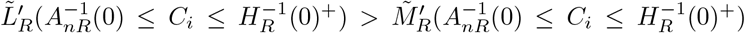 *under* 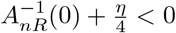,*and* 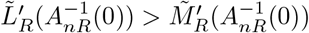.

*Proof*. In the case of [C1] and 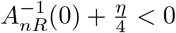,Eq. (SI 254) leads to

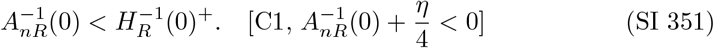

Combining Eq. (SI 351) with 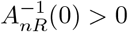 under [C1] (Table SI 1), we have:

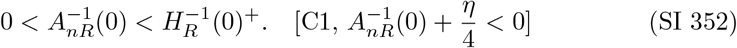

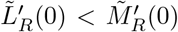 under [C1] and 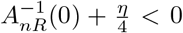 (Eq. (SI 316)) and 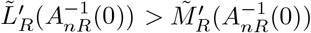, which is a condition of this proposition, lead to:

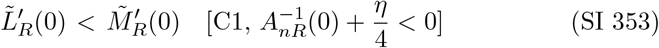

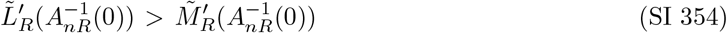

Eqs. (SI 353) and (SI 354) indicate that there is at least one intersection between 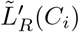 and 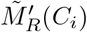 in the range 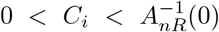.Considering Proposition 4 together with Eqs. (SI 353) and (SI 354), we conclude that there is exactly one intersection between 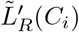 and 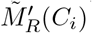 in the range 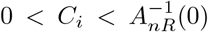,and no intersections in the range 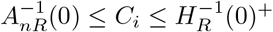.Therefore, we have

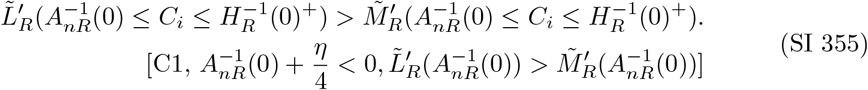

##### Proposition 6.

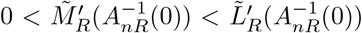 *under [C1]*, 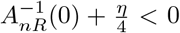 *and* 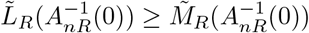

*Proof*. Eq. (SI 252) shows that 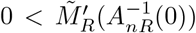 under [C1] and 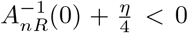.

Therefore, what remains to be proven is that 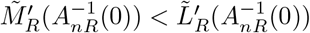.

Using Eqs. (SI 198) and (SI 233), we can express the inequality 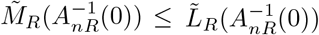, which is a condition of this proposition, as follows:

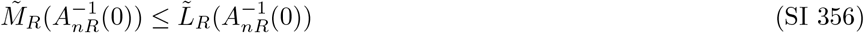

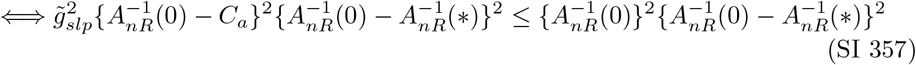

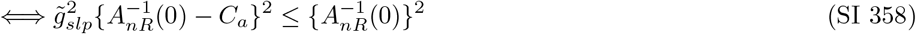

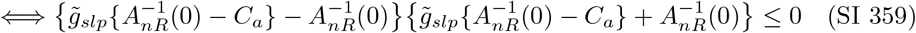

Since 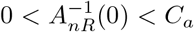 under [C1] (Table SI 1), the first term of the product on the left-hand side of Eq. (SI 359) is negative, as follows:

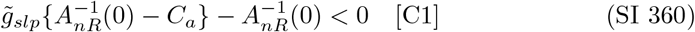

From Eqs. (SI 359) and (SI 360), we have:

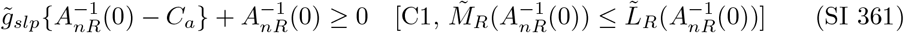

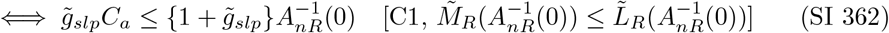

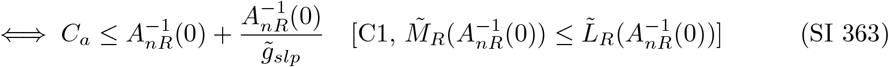

Eq. (SI 363) provides the upper limit of *C*_*a*_ in the case where [C1] and 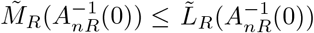.

Next, we derive a similar inequality for *C*_*a*_ from 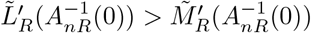.Using Eqs. (SI 207) and (SI 250), we rearrange the inequality 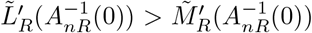, as follows:

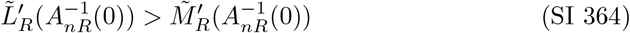

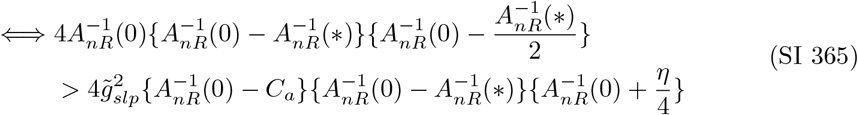

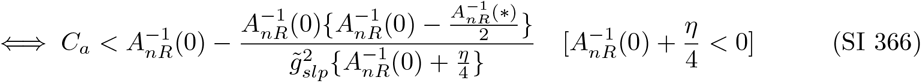

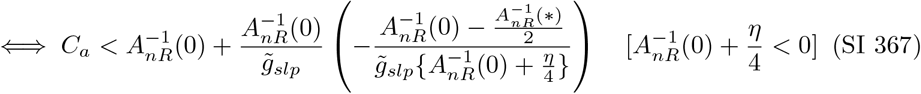

Note that 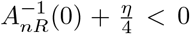 is assumed to obtain Eqs. (SI 366) and (SI 367). Thus, 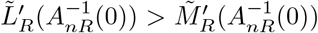 holds under 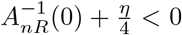 if Eq. (SI 367) holds under 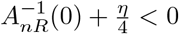. By comparing the terms on the right-hand side of Eqs. (SI 363) and (SI 367), we assume the following inequality:

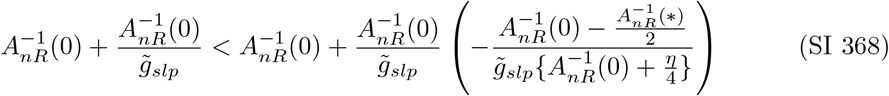

Eq. (SI 368) holds under [C1] where 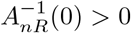 if the following inequality holds:

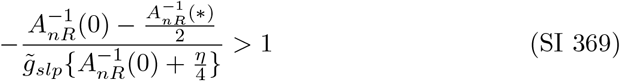

Rearranging Eq. (SI 369), we obtain:

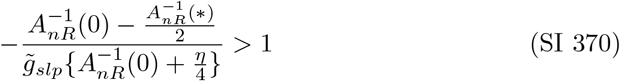

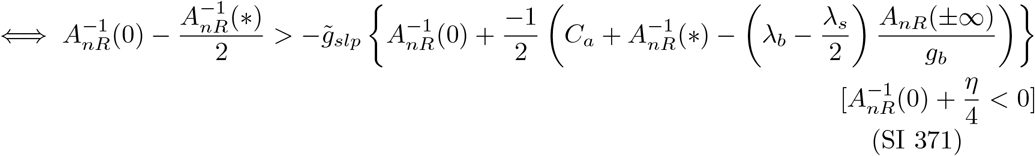

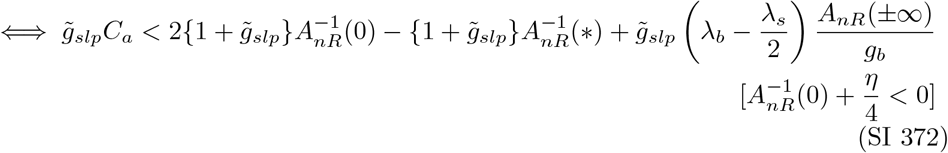

Note that 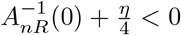 is assumed to derive Eq. (SI 371).

Considering that 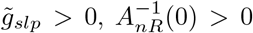,and *A*_*nR*_(*±*∞) *>* 0 under [C1] (Table SI 1), we can derive the following valid inequality:

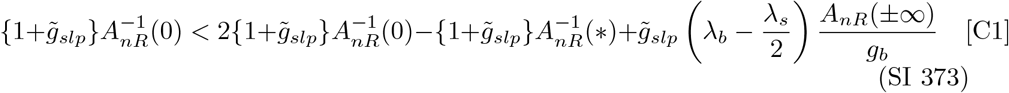

Eq. (SI 373) confirms that Eq. (SI 372) holds if Eq. (SI 362) is valid. Indeed, Eq. (SI 362) is valid as it is derived from 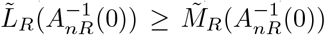,which is one of the conditions for the proposition. Therefore, Eq. (SI 372) is valid under [C1], 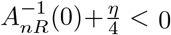, and 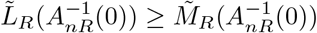.

The validity of Eq. (SI 372) implies that Eq. (SI 369) also holds (Eqs. (SI 369) to (SI 372)). Thus we have:

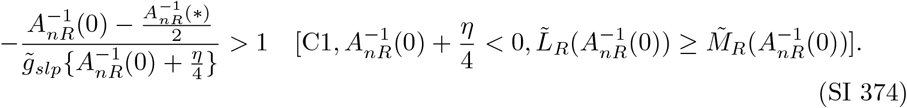

Considering Eq. (SI 374), it follows that Eq. (SI 368) is valid. Combining Eq. (SI 368) with Eq. (SI 363), we obtain:

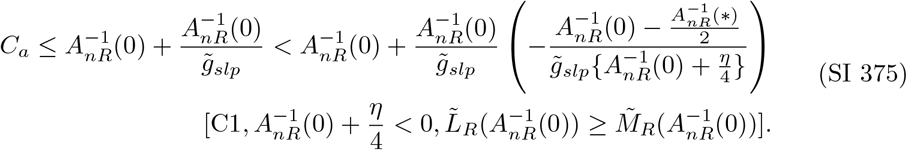

Eq. (SI 375) demonstrates that Eq. (SI 367), which is equivalent to Eq. (SI 364), is valid under 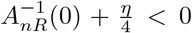, and 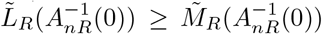.Therefore, Proposition 6 is proven.

#### SI 5.4.10 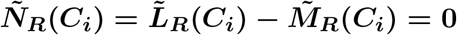

The roots of 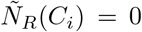,denoted as 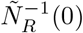,represent the intersections between 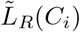 and 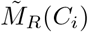 (Eq. (SI 192)). To analyze these intersections, we categorize the geometric patterns of 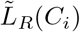 and 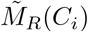 into four cases based on the relationships between 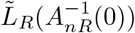 and 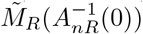,as well as their derivatives. These cases are labeled [P1] through [P4], and Table SI 41 provides the specific conditions for each pattern.

**Table SI 41:** Five geometric patterns for 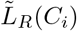 and 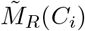.

| Case | $\tilde{L}_R(A_{nR}^{-1}(0))$ and $\tilde{M}_R(A_{nR}^{-1}(0))$ | $\tilde{L}'_R(A_{nR}^{-1}(0))$ and $\tilde{M}'_R(A_{nR}^{-1}(0))$ |
| --- | --- | --- |
| P1 | $\tilde{L}_R(A_{nR}^{-1}(0)) < \tilde{M}_R(A_{nR}^{-1}(0))$ | $\tilde{M}'_R(A_{nR}^{-1}(0)) \leq 0 < \tilde{L}'_R(A_{nR}^{-1}(0))$ |
| P2 | $\tilde{L}_R(A_{nR}^{-1}(0)) \geq \tilde{M}_R(A_{nR}^{-1}(0))$ | $\tilde{M}'_R(A_{nR}^{-1}(0)) \leq 0 < \tilde{L}'_R(A_{nR}^{-1}(0))$ |
| P3 | $\tilde{L}_R(A_{nR}^{-1}(0)) < \tilde{M}_R(A_{nR}^{-1}(0))$ | $0 < \tilde{M}'_R(A_{nR}^{-1}(0))$ |
| P4 | $\tilde{L}_R(A_{nR}^{-1}(0)) \geq \tilde{M}_R(A_{nR}^{-1}(0))$ | $0 < \tilde{M}'_R(A_{nR}^{-1}(0)) < \tilde{L}'_R(A_{nR}^{-1}(0))$ |

It is important to note that Proposition 6 shows that 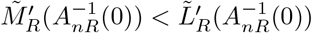 when 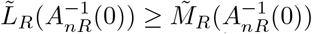 and 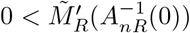 (Eq. (SI 252)). Additionally, we focus on the range 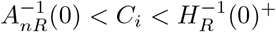,which is the possible range of appropriate solutions under [C1] (Table 2).

(P1) 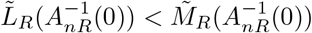 and 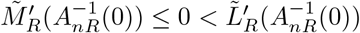

Eq. (SI 252) shows that 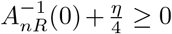 holds when 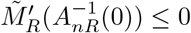 under [C1].

Thus, we assume 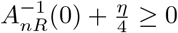 in [P1].

Given 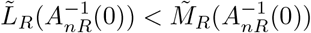,which is one of the conditions in [P1], along with 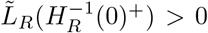 (Eq. (SI 199)), 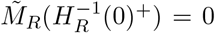 (Eq. (SI 229)), and Proposition 2, we obtain:

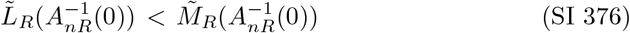

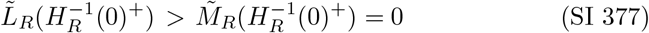

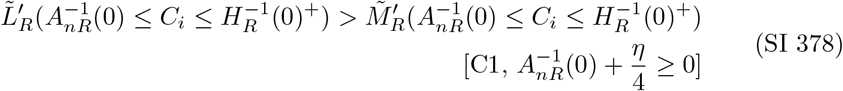

Therefore, there is only one solution between 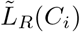 and 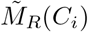 in the range 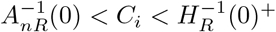.Thus, we have:

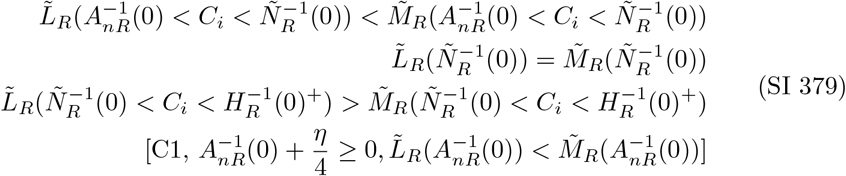

Hence,

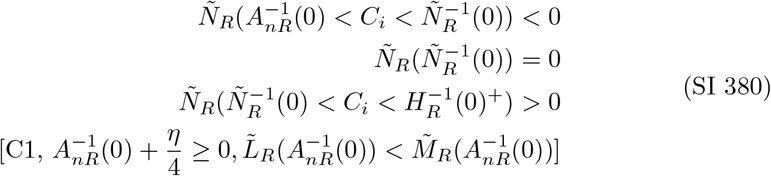

Figures SI 19 and SI 20 illustrate the curves of 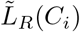 and 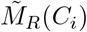,as well as 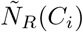 for case [P1].

**Fig. SI 19:**
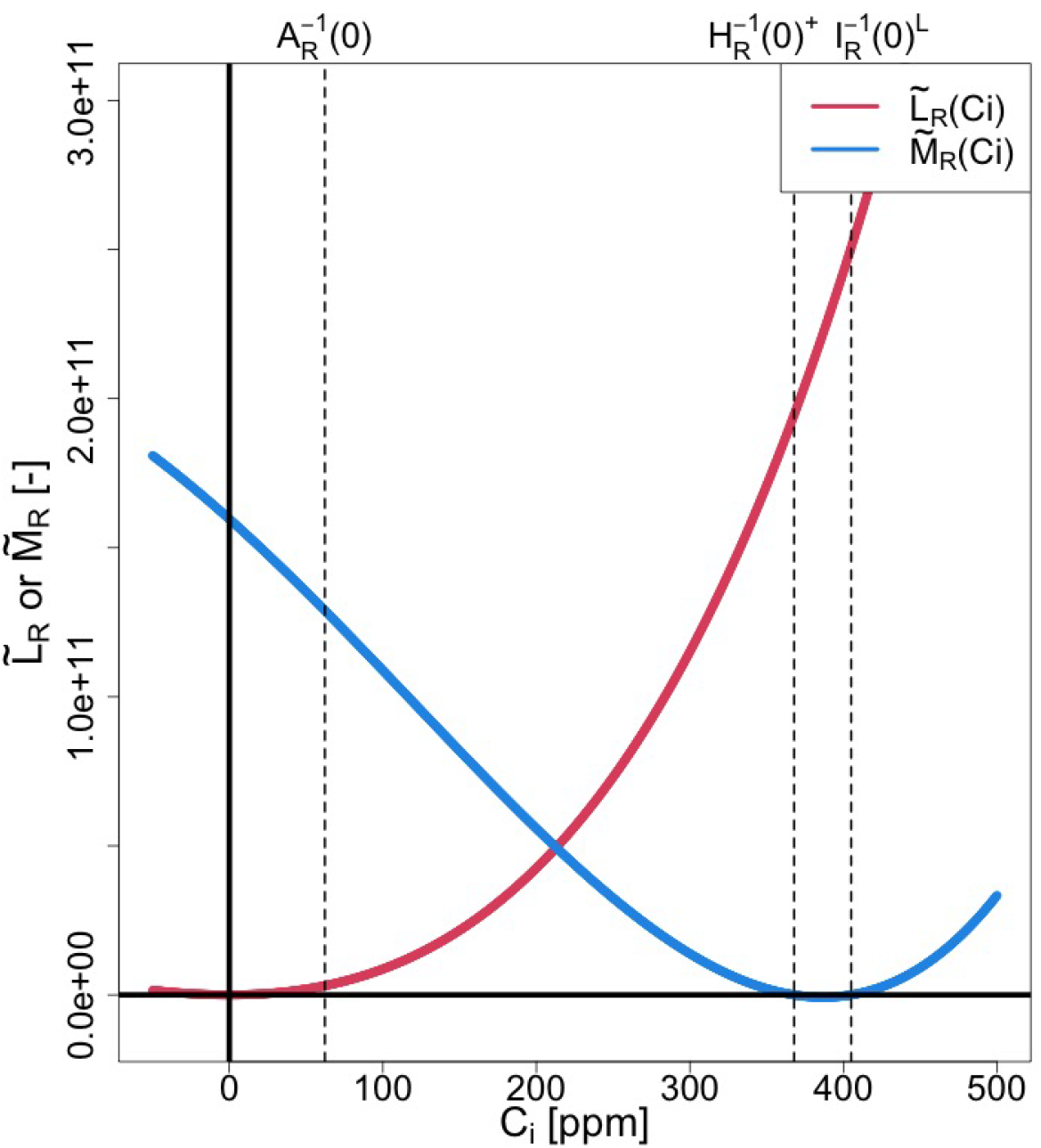
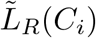 and 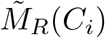 for [P1]

**Fig. SI 20:**
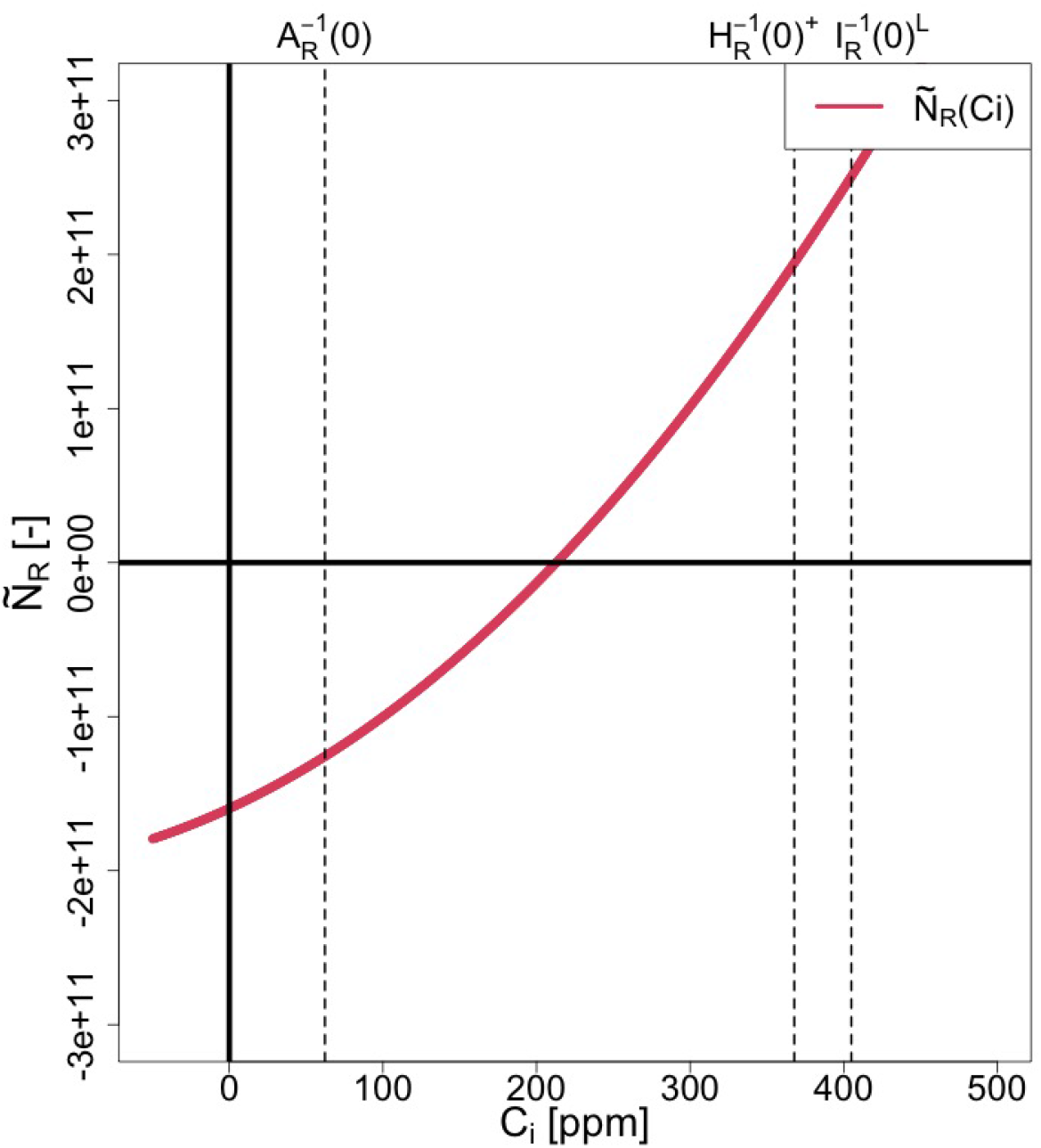
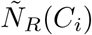 for [P1]

(P2) 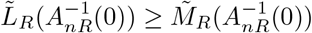 and 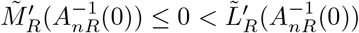

Similar to the case in [P1], we can assume 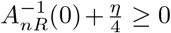 from 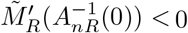 (Eq. (SI 252)).

Given 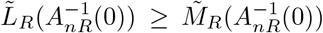,which is one of the conditions in (P2), along with 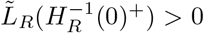 (Eq. (SI 199)), 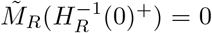 (Eq. (SI 229)), and Proposition 2, we obtain:

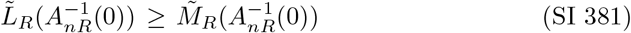

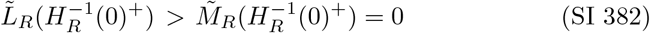

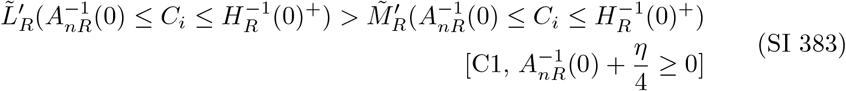

Therefore, there is no solution between 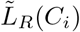 and 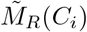 in the range 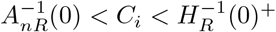. Thus, we have:

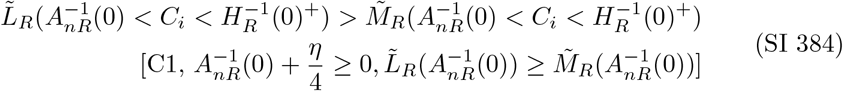

Hence,

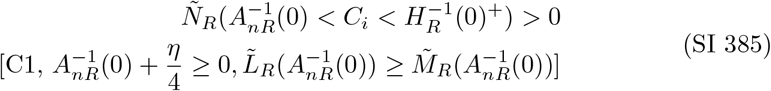

Figures SI 21 and SI 22 illustrate the curves of 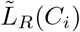 and 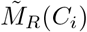,as well as 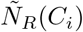 for case [P2].

**Fig. SI 21:**
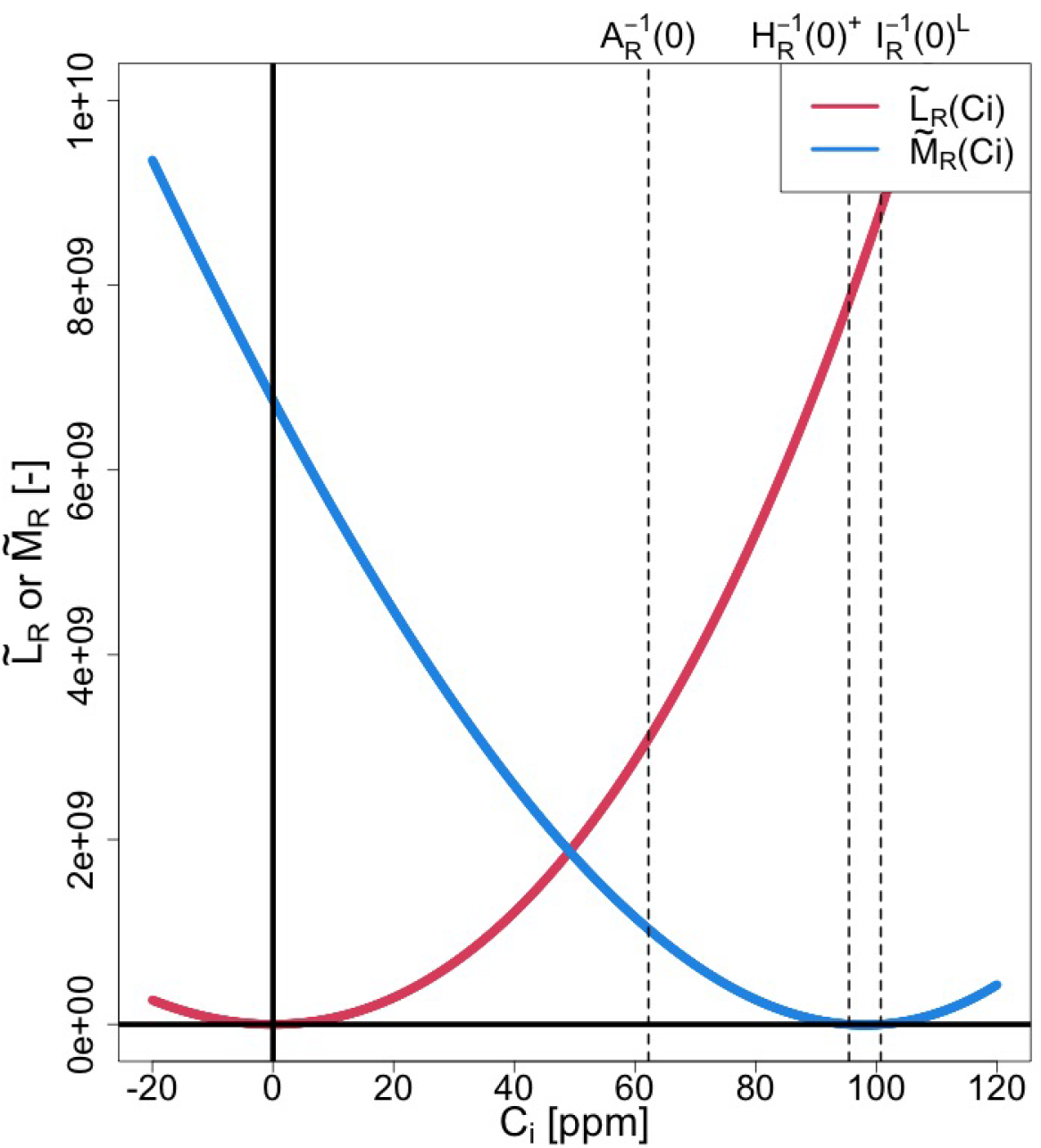
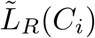 and 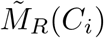 for [P2]

**Fig. SI 22:**
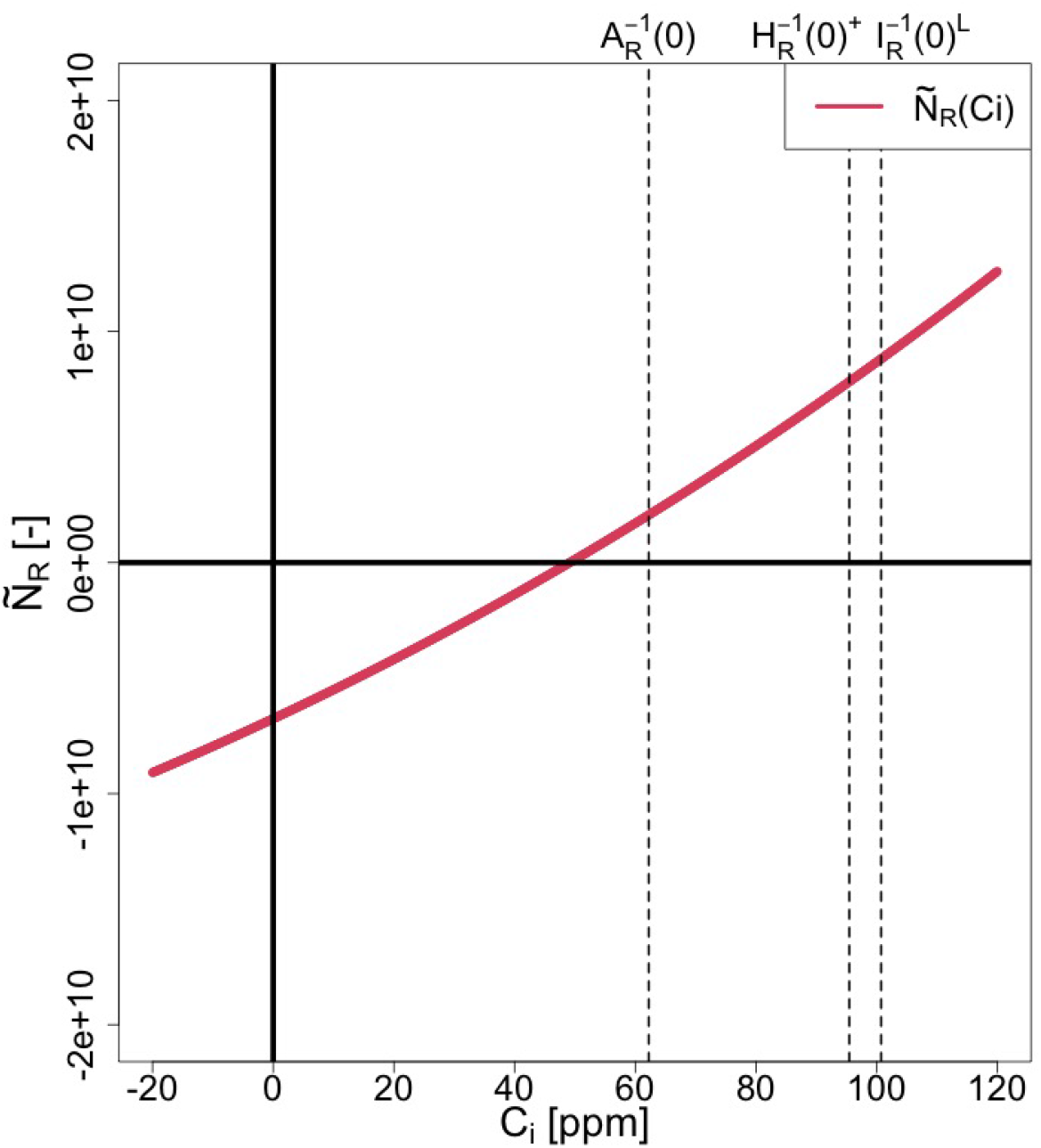
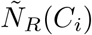 for [P2]

(P3) 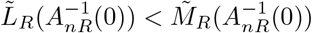 and 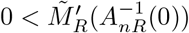

Eq. (SI 252) shows that 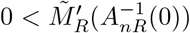 corresponds to 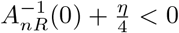 under [C1]. Thus, [C1] and 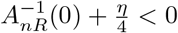 are assumed in case P3.

From 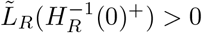 (Eq. (SI 199)) and 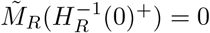 (Eq. (SI 229)), combined with 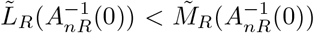,which is the first condition of [P3], we obtain:

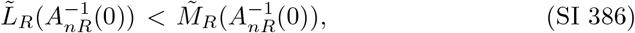

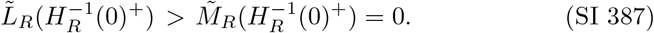

Therefore, there are one or more intersections between 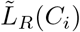 and 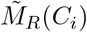 within 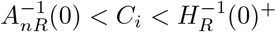.

Proposition 4 demonstrates that there is exactly one intersection between 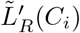 and 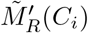 within 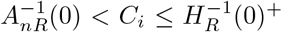 under [C1] and 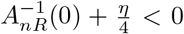. This intersection is denoted as 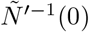.From 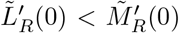 (Eq. (SI 316)) and 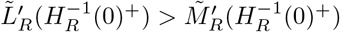 (Eq. (SI 314)) under [C1] and 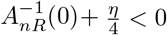, we have:

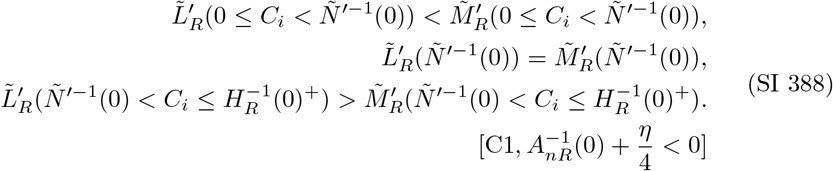

We split the analysis into two cases: (A): 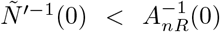 and (B):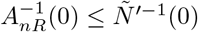

(A) 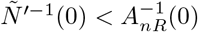 In this case, Eq. (SI 388) implies:

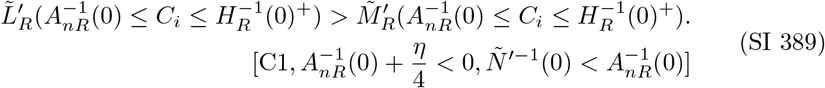 Combining this with Eqs. (SI 386) and (SI 387), we find:

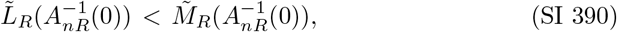

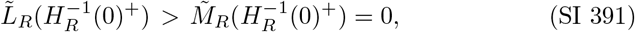

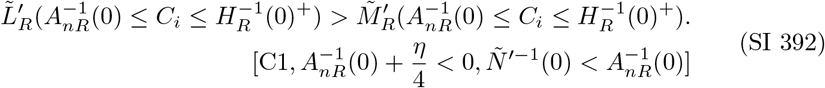 Thus, there is exactly one intersection between 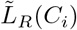 and 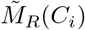 within 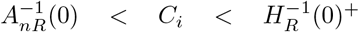 under [C1], 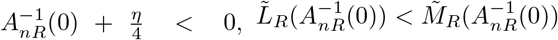 and 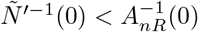.
(B) 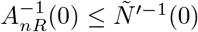 We divide the range 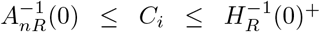 into two sub-ranges: (i) 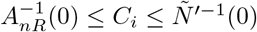,and (ii) 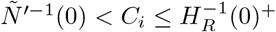.
  i. 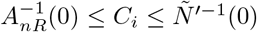 From Eqs. (SI 386), and (SI 388), we have:

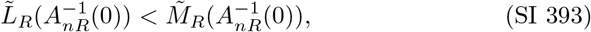

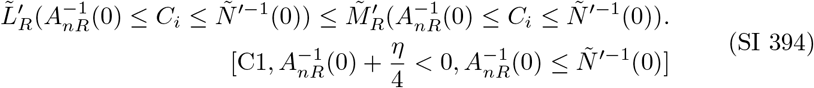 Therefore, there is no intersection between 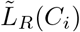 and 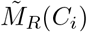 within 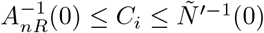 under [C1], 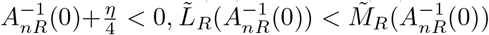, and 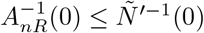.Thus, we find:

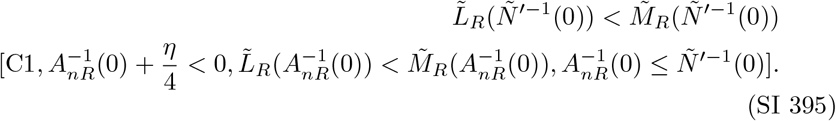
  ii. 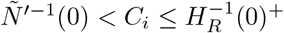 From Eqs. (SI 395), (SI 387), and (SI 388), we have:

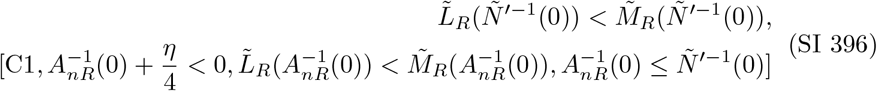

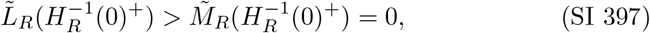

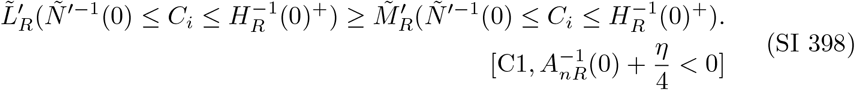 Therefore, there is one intersection between 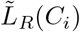 and 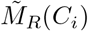 within 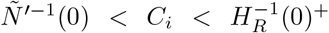 under [C1], 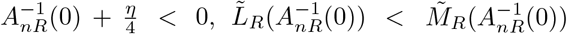 and 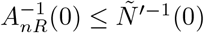. The results of the two sub-ranges (i) and (ii) show that there is one intersection between 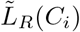 and 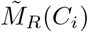 within 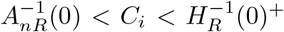 under [C1],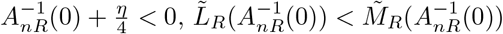 and 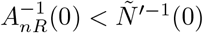. Consequently, the results of cases (A) and (B) show that there is exactly one intersection between 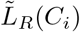 and 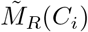 within 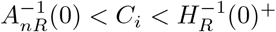 under [C1], 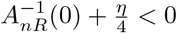,and 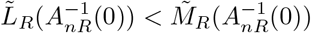.Thus, we find:

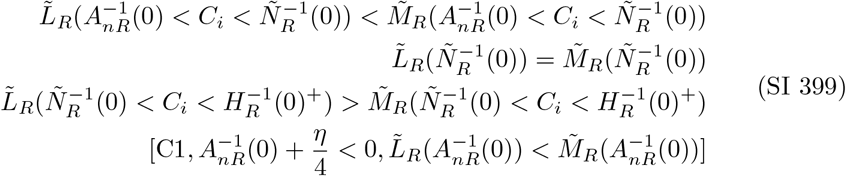

Hence,

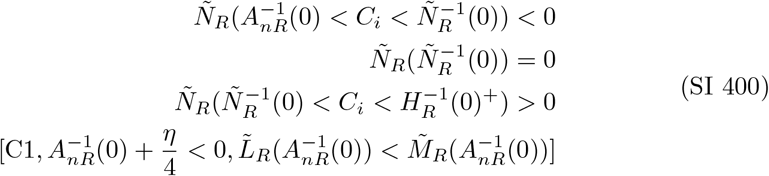

Figures SI 23 and SI 24 show the curves of 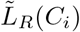 and 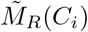,as well as 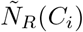 for case [P3].

**Fig. SI 23:**
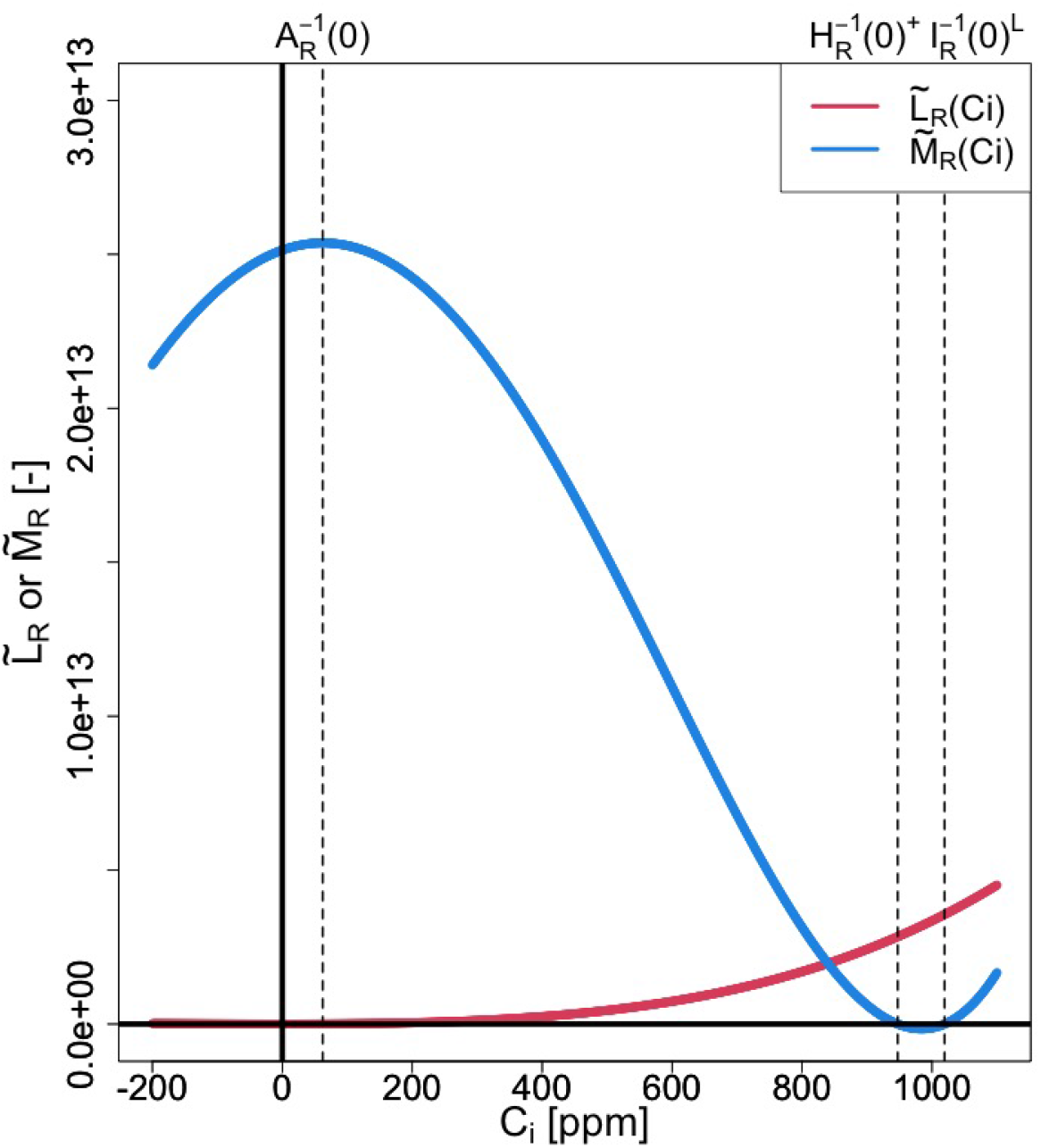
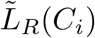 and 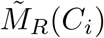 for [P3]

**Fig. SI 24:**
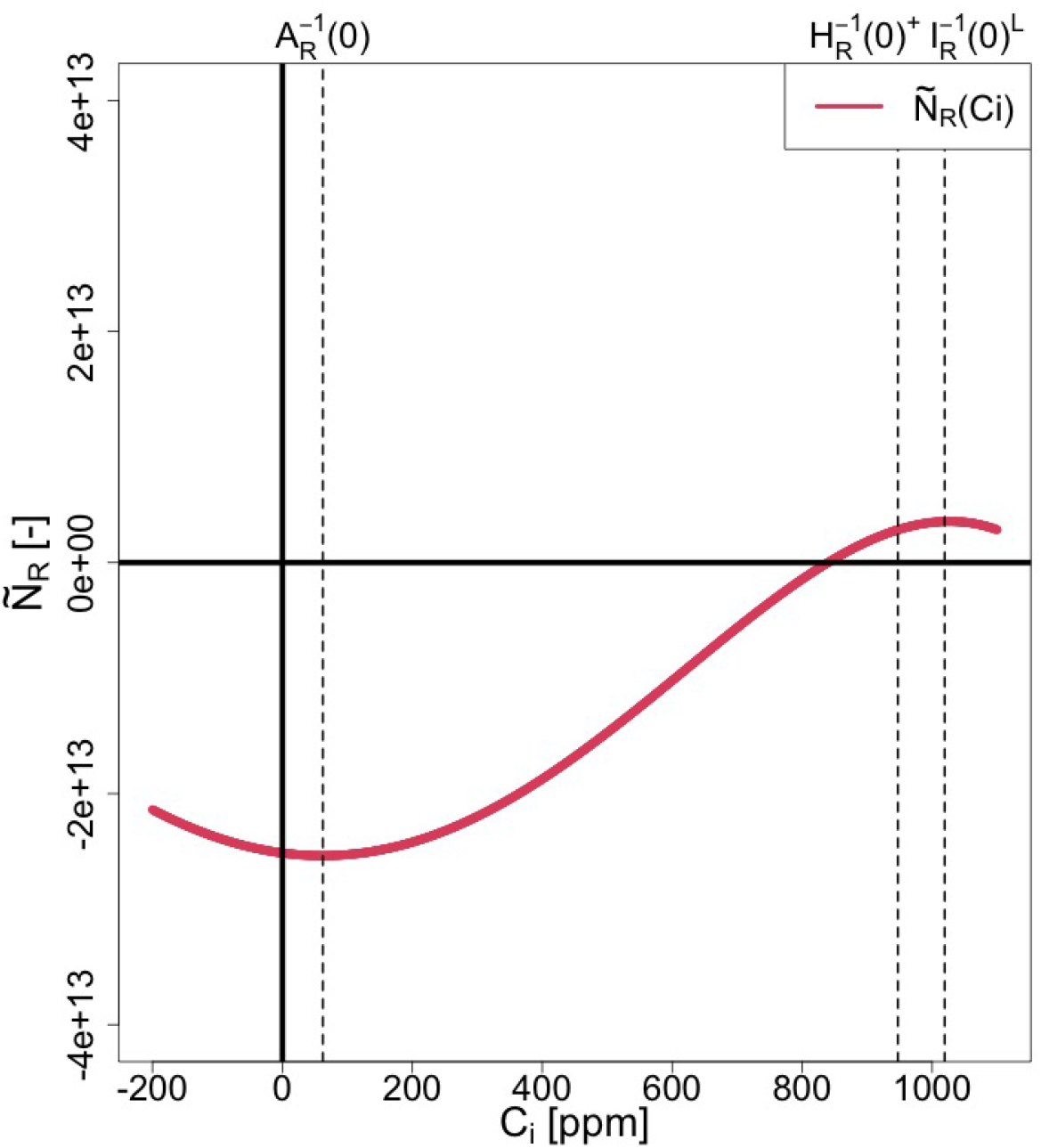
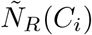 for [P3]

(P4) 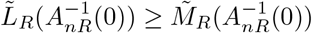 and 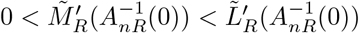 Similar to case [P3], [C1] and 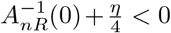 are assumed since 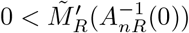 (Eq. (SI 252)). Applying 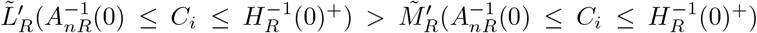 under [C1], 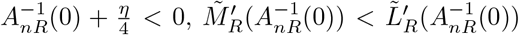 (Proposition 5), and 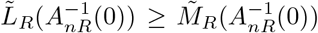,along with 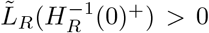 (Eq. (SI 199)) and 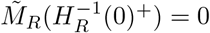 (Eq. (SI 229)), we have:

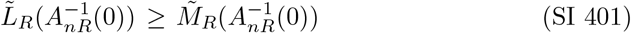

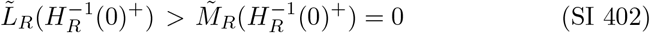

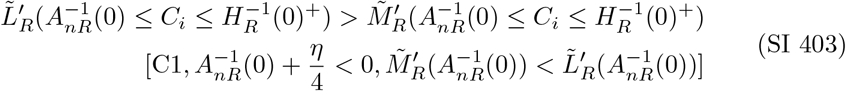

Therefore, there is no intersection between 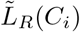 and 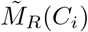 within the range 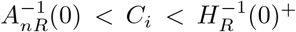 under [C1], 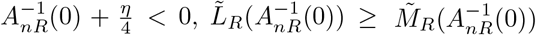 and 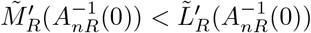.Thus, we have:

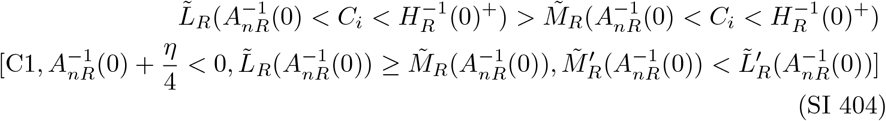

Hence,

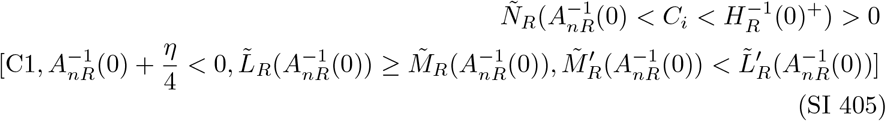

Figures SI 25 to SI 27 show the curves of 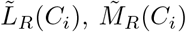,and 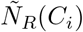 for case [P4]

**Fig. SI 25:**
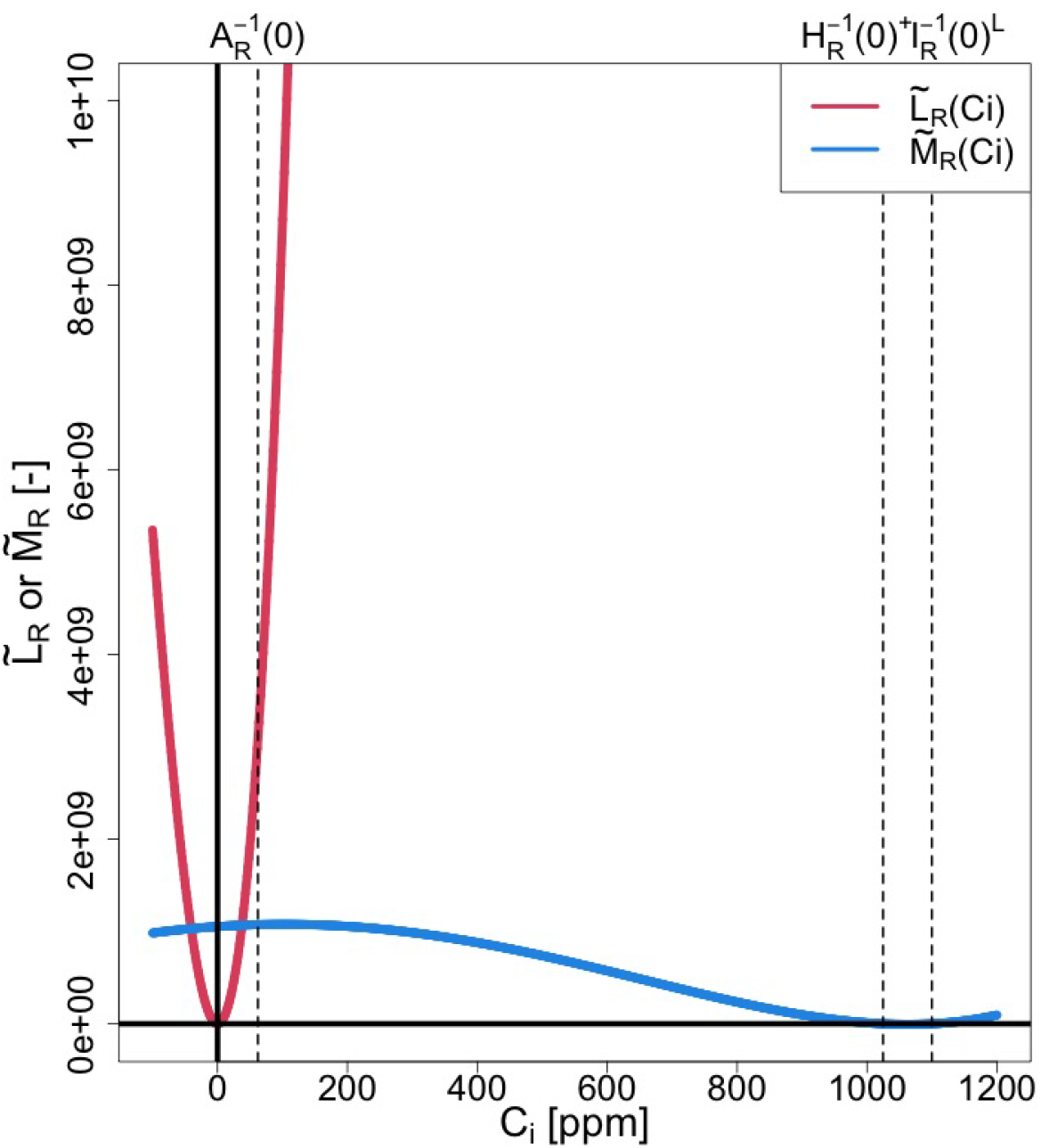
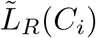 and 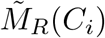 for [P4]

**Fig. SI 26:**
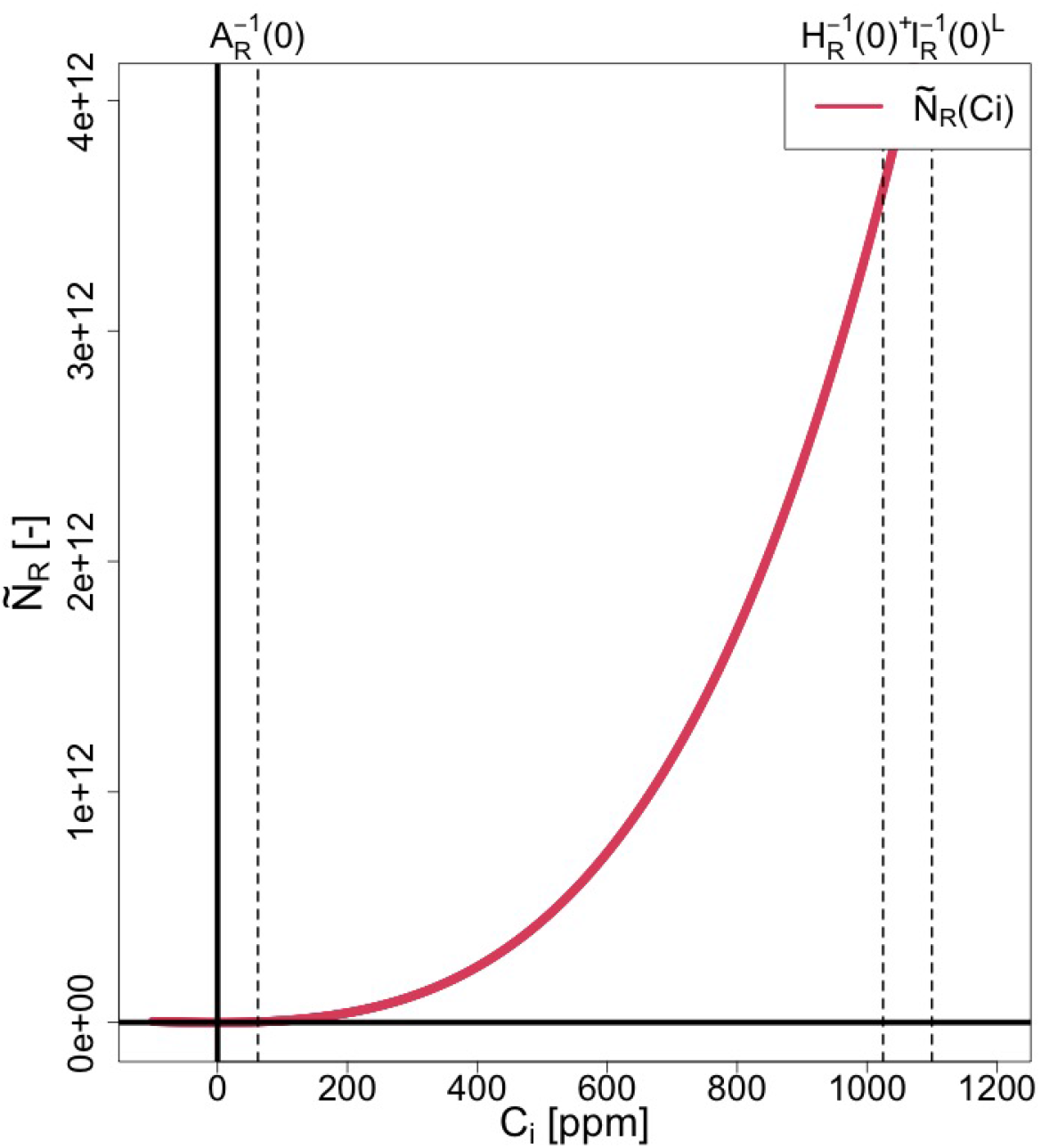
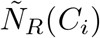 for [P4]

**Fig. SI 27:**
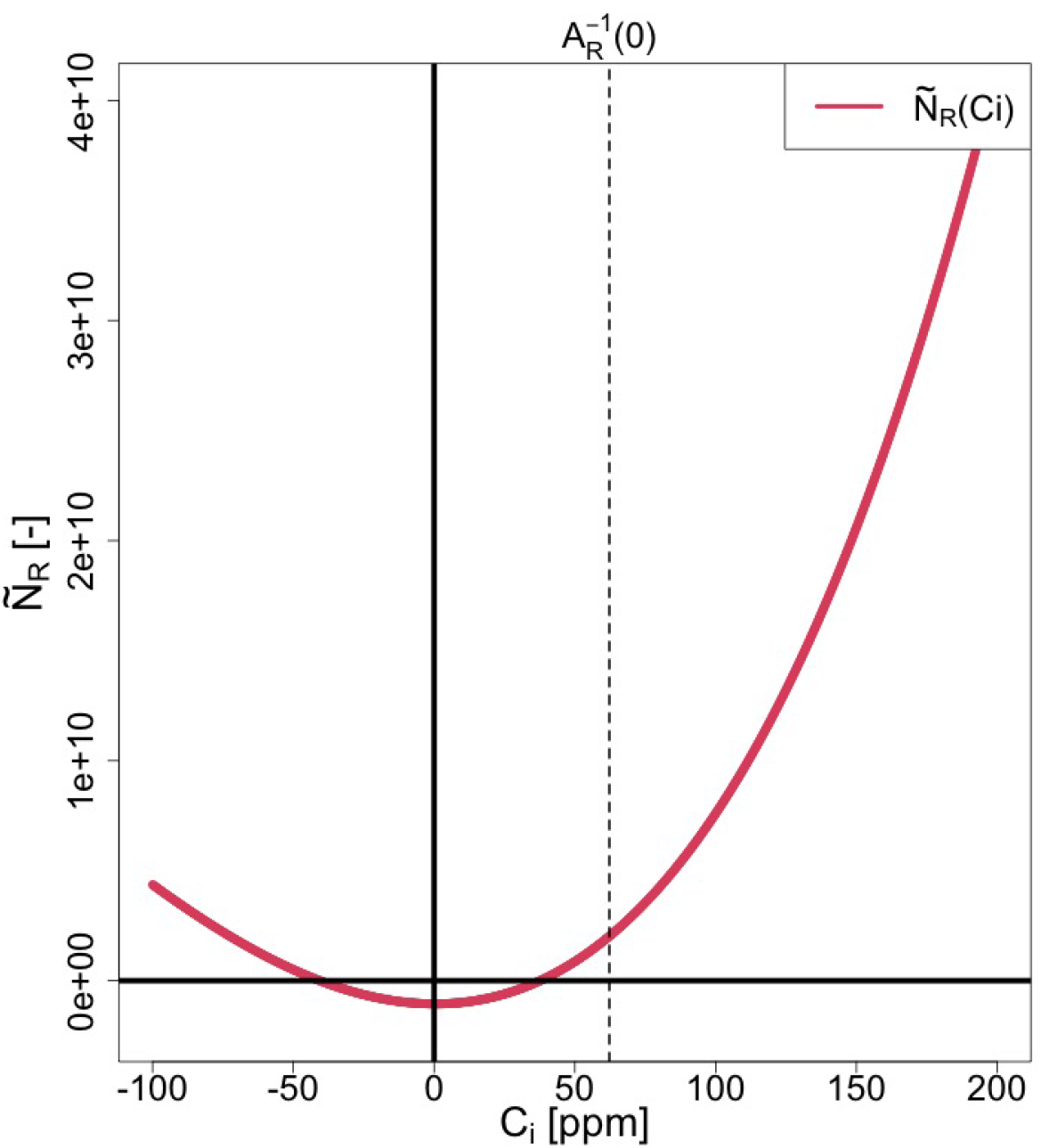
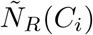 at – 100 < *C*_*i*_ < 200 for [P4]

Table SI 42 summarizes the number of intersections between 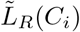 and 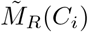 within the range 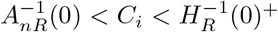 for all four geometric patterns from P1 to P4.

**Table SI 42:** The number of intersections between 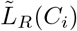 and 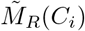 at 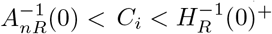.

| Cases | # of Intersections | Type (ZI or ZII) |
| --- | --- | --- |
| P1 | 1 | ZI |
| P2 | 0 | ZII |
| P3 | 1 | ZI |
| P4 | 0 | ZII |

Table SI 42 reveals two distinct geometric patterns: one pattern has one intersection, while the other has none within the range 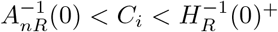.We denote these patterns as [ZI] for the case with one intersection and [ZII] for the case with none. Table SI 42 also categorizes each case as either [ZI] or [ZII]. We can summarize the sign behavior of 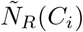 within 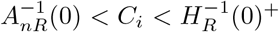 for cases [ZI] and [ZII], as follows:

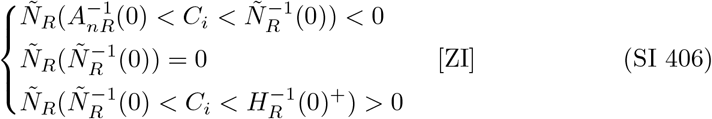

and,

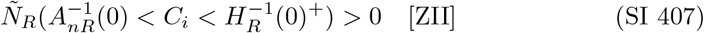

Since 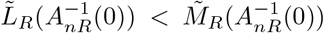 under [ZI] and 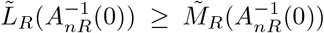 under [ZII] (Table SI 41), we obtain the following inequalities:

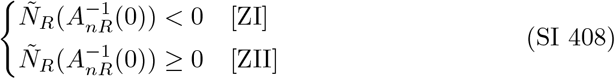

In case [C1], Eqs. (SI 356) to (SI 363) lead to:

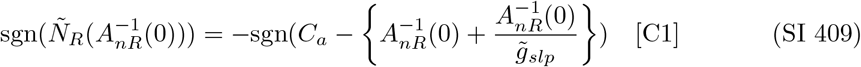

Therefore, Eqs. (SI 408) and (SI 409) give conditions for cases [ZI] and [ZII] as:

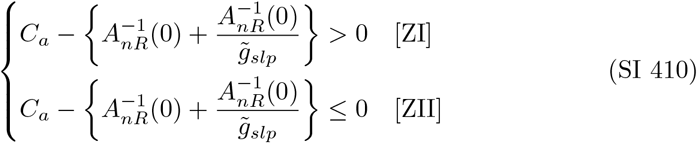

Finally, given that 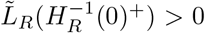 (Eq. (SI 199)) and 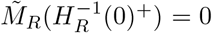 (Eq. (SI 229)), we conclude:

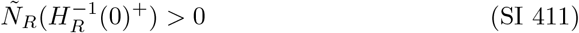

### SI 5.5 *N*_*R*_(*C*_*i*_)

*N*_*R*_(*C*_*i*_) is given from Eq. (SI 138) as follows:

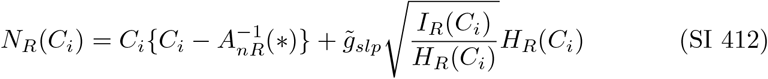

Since the roots and signs of *N*_*R*_(*C*_*i*_) are the same as those of 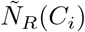 at 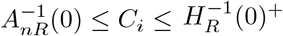 under [C1] (Eqs. (SI 195) and (SI 191)), Eqs. (SI 406) and (SI 407) lead to:

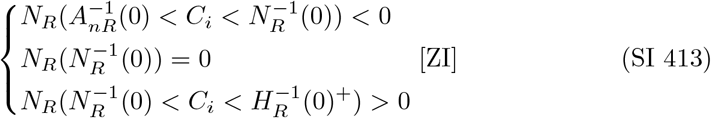

and,

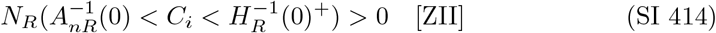

Since 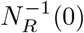 exist at 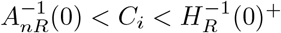 under [ZI], we have

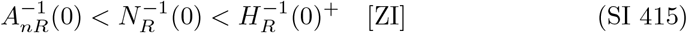

In addition, Eqs. (SI 408) and (SI 411) lead to:

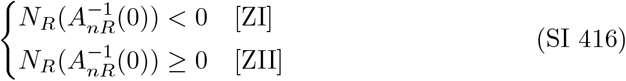

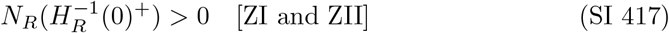

Figures SI 28 and SI 29 show the curves of *N*_*R*_(*C*_*i*_) for [ZI] and [ZII]. Tables SI 43 and SI 44 show the sign charts of *N*_*R*_(*C*_*i*_) for [ZI] and [ZII].

**Fig. SI 28:**
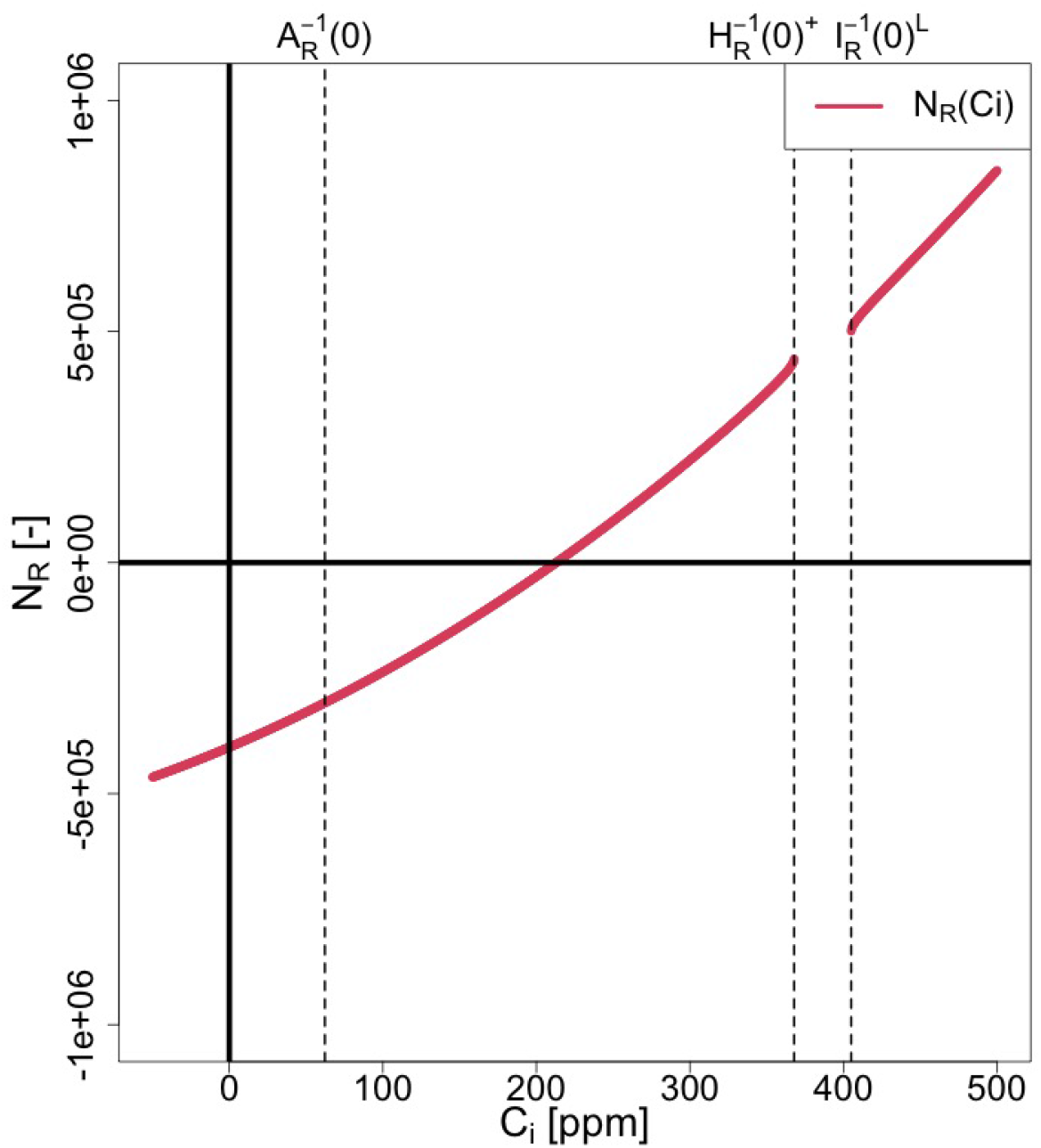
*N*_*R*_(*C*_*i*_) for [ZI]

**Fig. SI 29:**
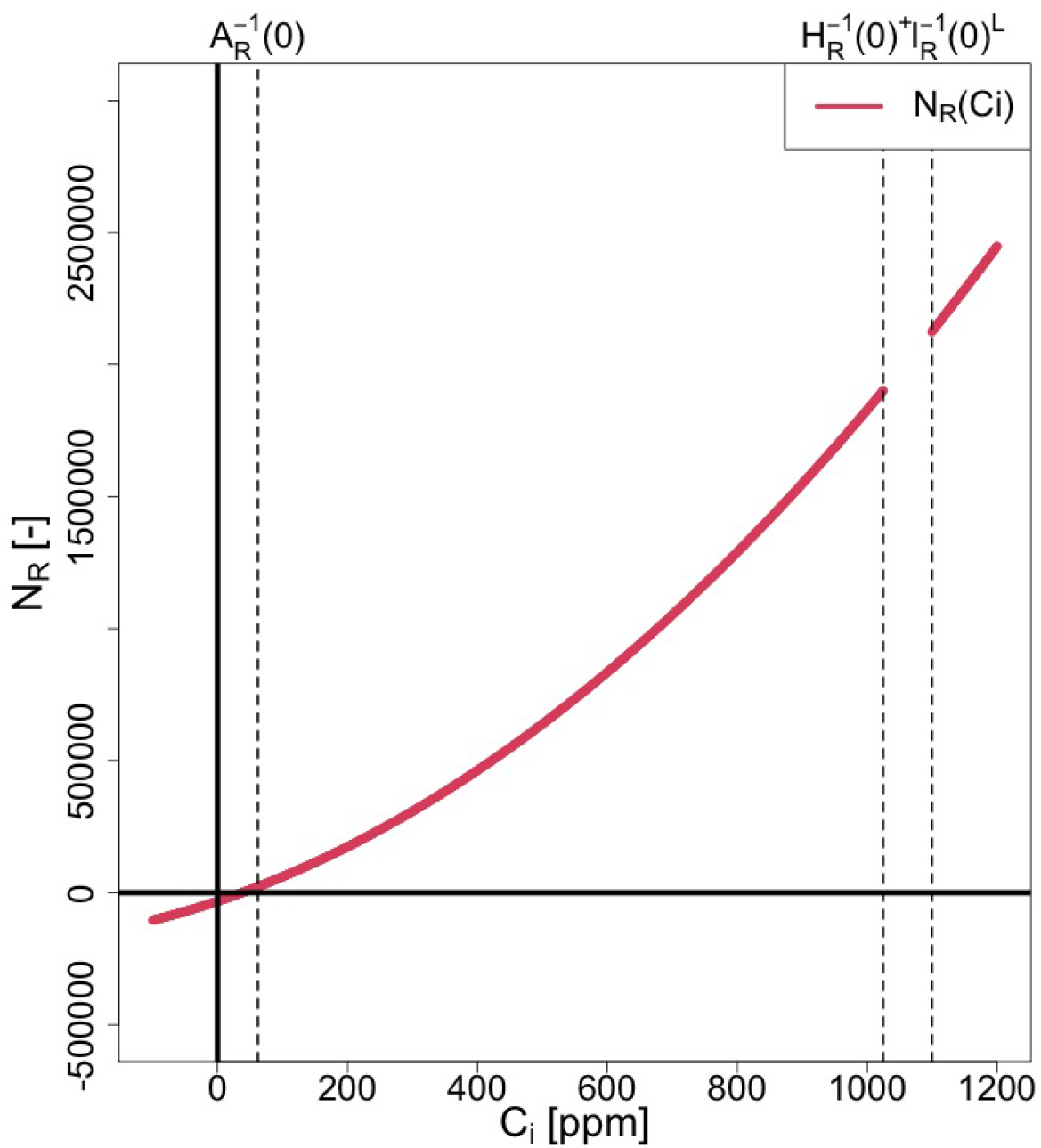
*N*_*R*_(*C*_*i*_) for [ZII]

**Table SI 43:**
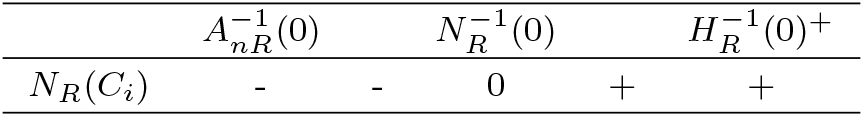
Sign chart of *N*_*R*_(*C*_*i*_) for ZI.

| | $A_{nR}^{-1}(0)$ | | $N_R^{-1}(0)$ | | $H_R^{-1}(0)^+$ |
| --- | --- | --- | --- | --- | --- |
| $N_R(C_i)$ | - | - | 0 | + | + |

**Table SI 44:** Sign chart of *N*_*R*_(*C*_*i*_) for ZII.

| | $A_{nR}^{-1}(0)$ | | $H_R^{-1}(0)^+$ |
| --- | --- | --- | --- |
| $N_R(C_i)$ | 0 or + | + | + |

### SI 5.6 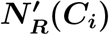 and 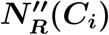

In this section, we focus on the region 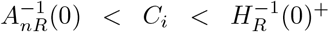,which is the possible range of solutions under [C1] (Table 2). Within this range, using 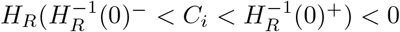 (Eq. (SI 33)) and 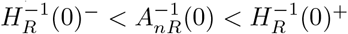 under [C1] (Eq. (SI 46)), we derive the following inequality:

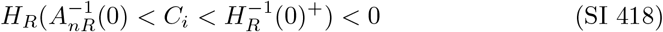

Additionally, from 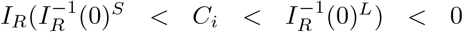 (Eq. (SI 156)) and 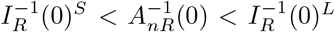 under [C1] (Eqs. (SI 167) to (SI 169)), we derive the following inequality:

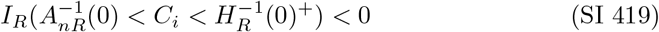

By differentiating Eq. (SI 138) with respect to *C*_*i*_, we obtain

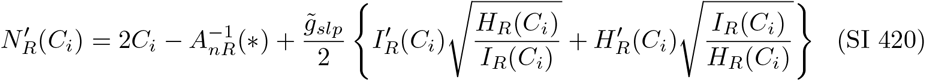

In deriving Eq. (SI 420), Eq. (SI 141) is used.

Differentiating Eq. (SI 420) and taking into account Eqs. (SI 418) and (SI 419), we obtain:

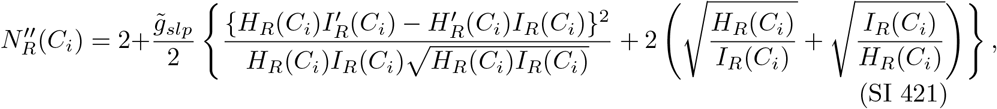

(a) Geometry of 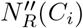 From Eq. (SI 421), we have

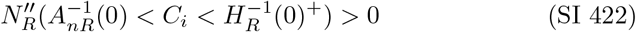 Figures SI 30 and SI 31 illustrate the curves of 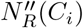 for [ZI] and [ZII].
(b) Singularity: 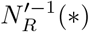 From Eq. (SI 420), 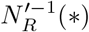 is given by:

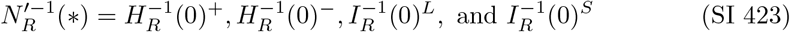
(c) 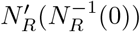 Since the sign of 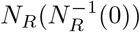 changes from negative to positive at 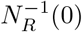 under [ZI] (Table SI 43), along with 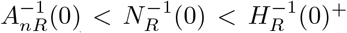 under [ZI] (Eq. (SI 415)) and 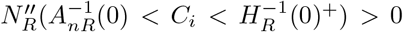 (Eq. (SI 422)), 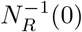 must be positive. Thus, we obtain:

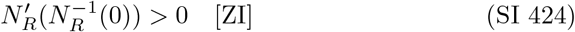
(d) 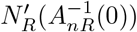

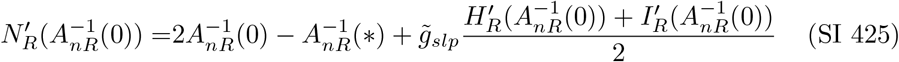

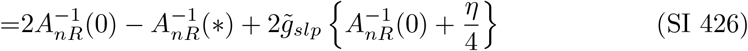 Since 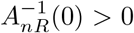 under [C1] (Table SI 1) and 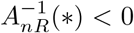 (Eq. SI 3), Eq. (SI 426) under [C1] and 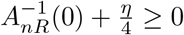 implies:

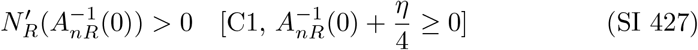 In the case where [C1], 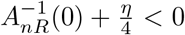 and 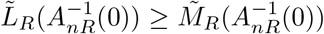,Eq. (SI 374) is valid. Using Eq. (SI 374), Eq. (SI 426) leads to:

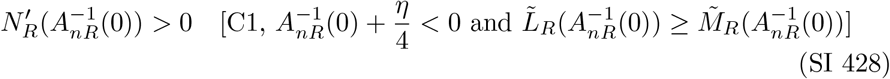

**Fig. SI 30:**
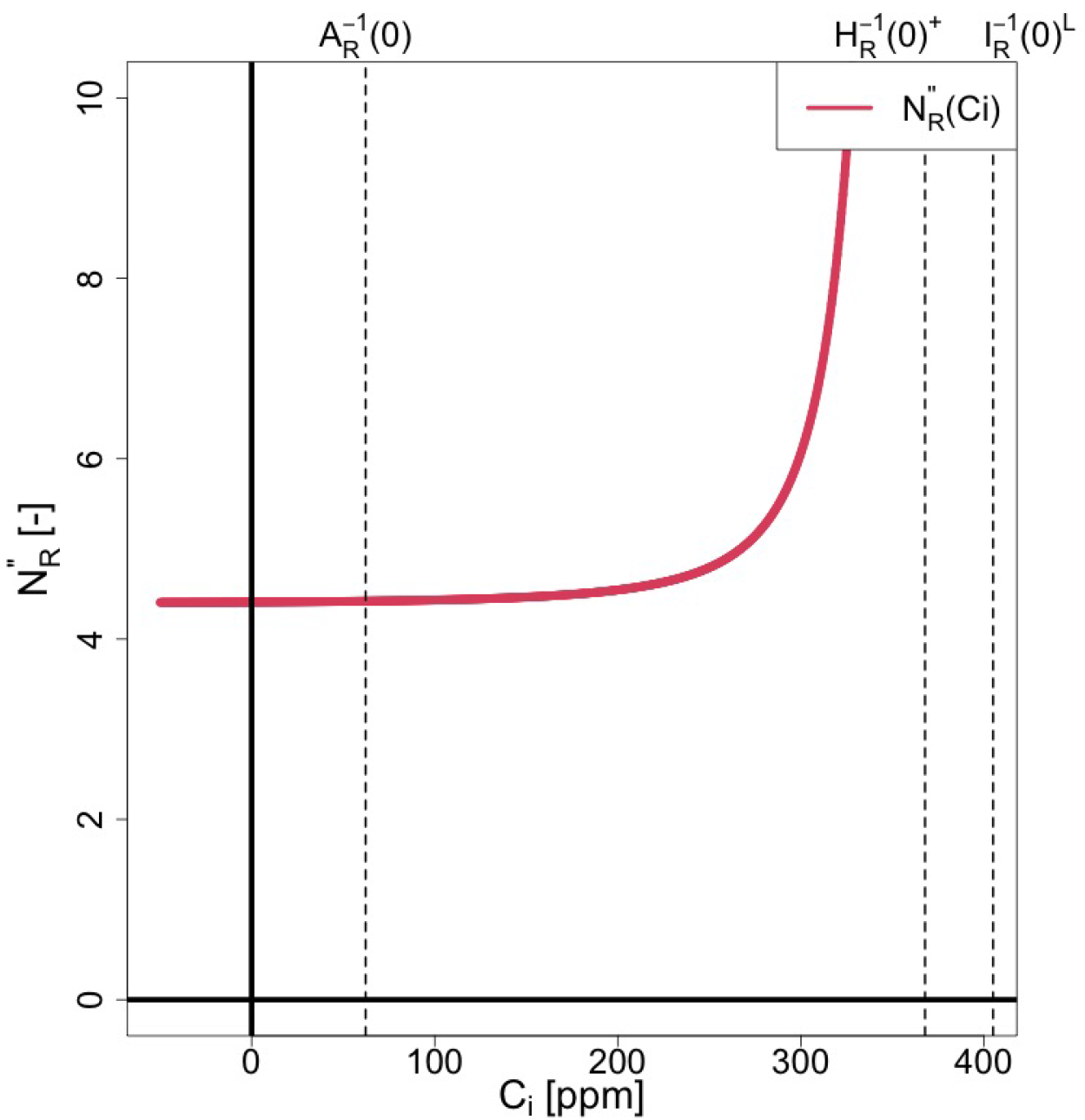
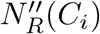 for [ZI]

**Fig. SI 31:**
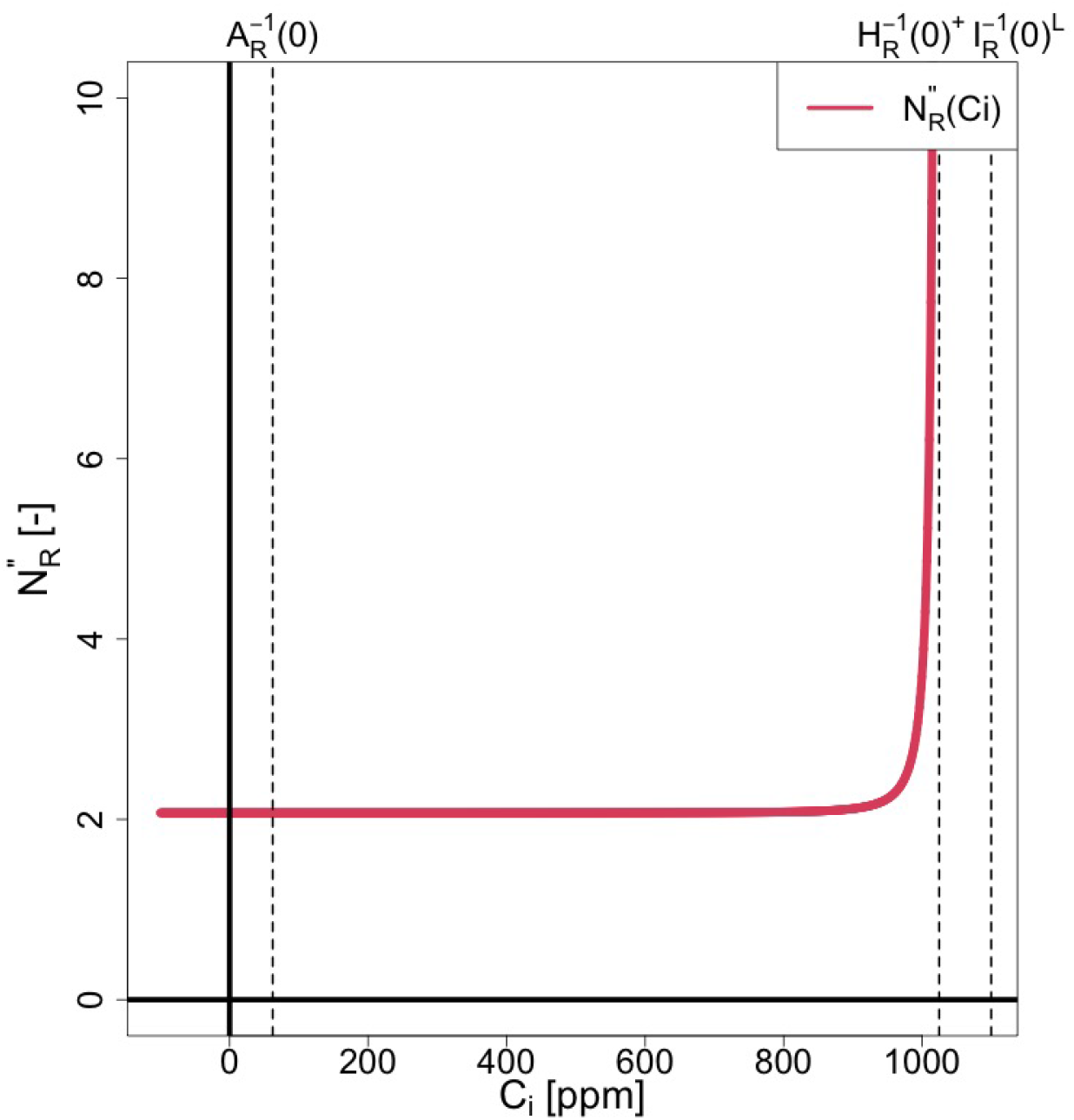
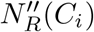 for [ZII]

The conditions for Eq. (SI 427) correspond to [P1] and [P2], while those for Eq. (SI 428) correspond to [P4] (Table SI 41 and Eq. (SI 252)). Thus, we have:

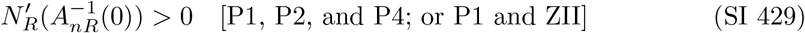

Then, we prove the following proposition:

#### Proposition 7.

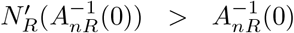 *under [C1] and* 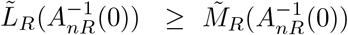.

*Proof*. Similar to Eq. (SI 373), we can derive the following inequality under [C1]:

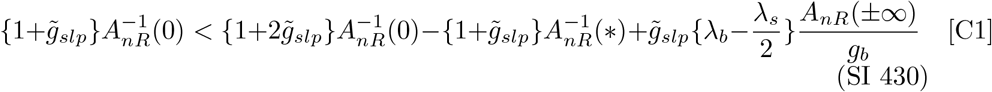

In the case where 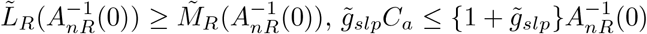 holds under [C1] (Eq. (SI 362)). Therefore, we obtain:

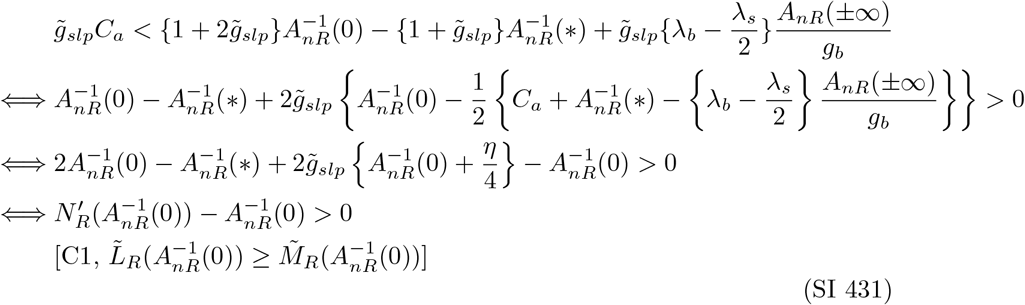

(e) Geometry of 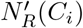 Figures SI 32 and SI 33 illustrate the curves of 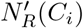 for [ZI] and [ZII].

**Fig. SI 32:**
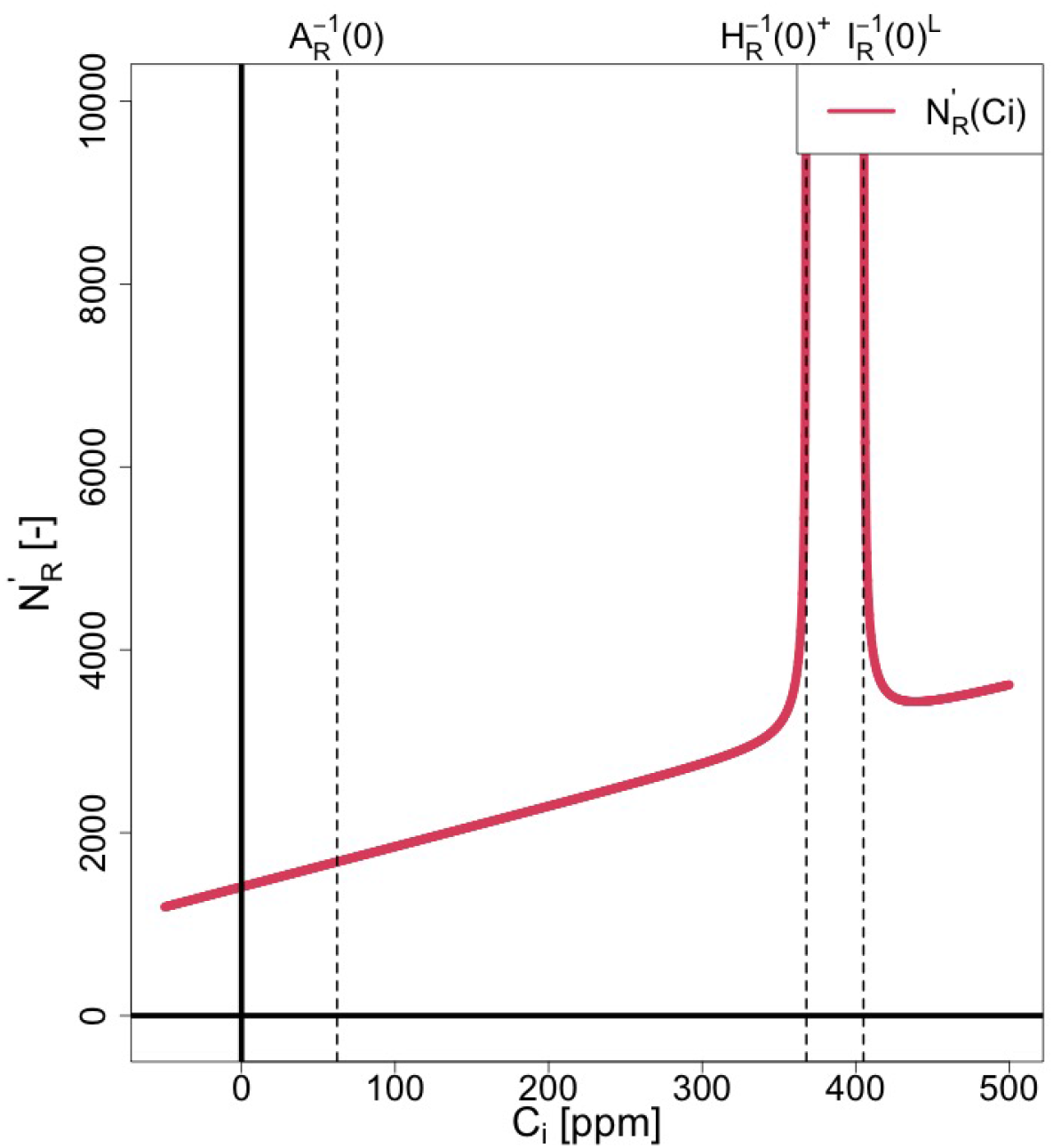
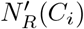 for [ZII]

**Fig. SI 33:**
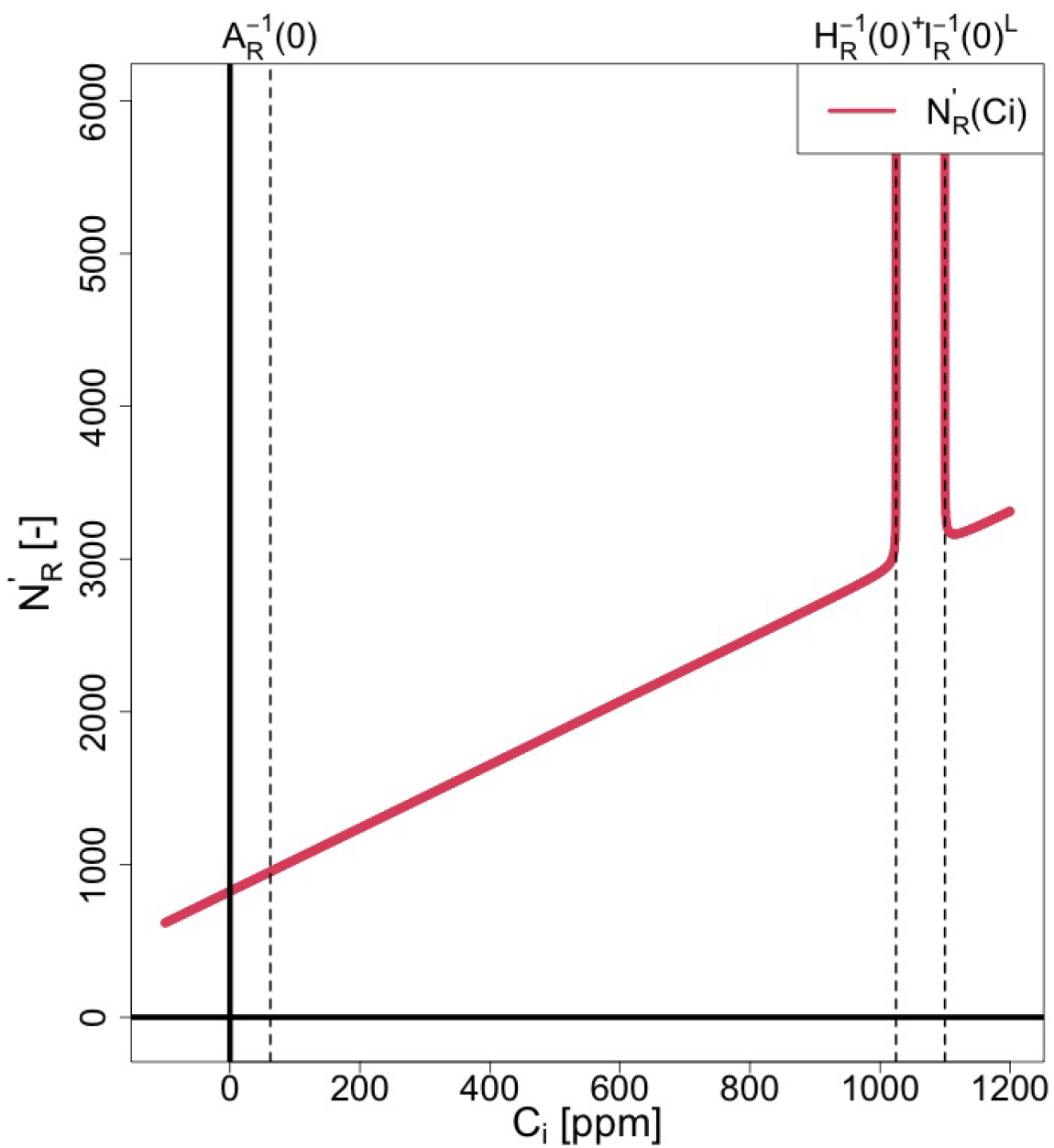
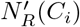 for [ZII]

#### SI 5.7 *Z*_*R*_(*C*_*i*_)

*Z*_*R*_(*C*_*i*_) is rewritten from Eq. (SI 137) as follows:

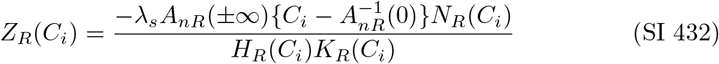

(a) Singularity: 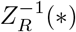 From Eq. (SI 432), we have 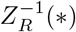 as follows:

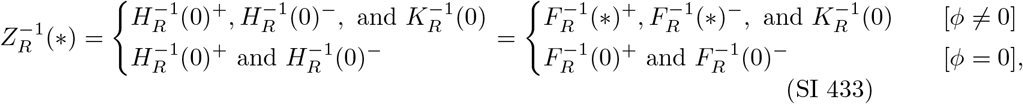

where 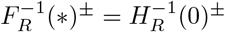 (Eqs. (SI 83) and (SI 84)) is used. Given that *A*_*nR*_(*±*∞) *>* 0 under [C1] (Table SI 1), 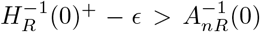 under [C1] (Eq. (SI 46)), 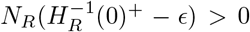 (Eqs. (SI 406) and (SI 407)), 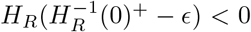 (Eq. (SI 33)), and 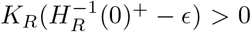 (Eq. (SI 182)), we obtain:

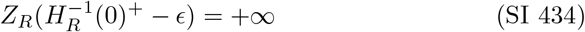
(c) Root: 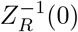 From Eq. (SI 432), we find two roots of *Z*_*R*_(*C*_*i*_) under [ZI] and one root under [ZII], as follows:

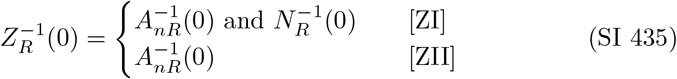 To distinguish between the two roots, we denote the roots corresponding to 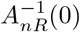 and 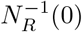 as follows:

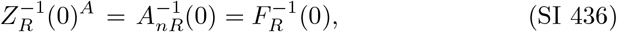

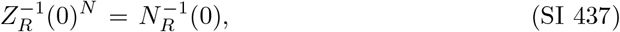

where 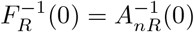 (Eq. (SI 89)) is used.
(d) Geometry: 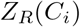 Figures 3a and 3b show the curves of *Z*_*RI*_ (*C*_*i*_) and *Z*_*RII*_ (*C*_*i*_), and Tables SI 45 and SI 46 show their sign charts.

**Table SI 45:**
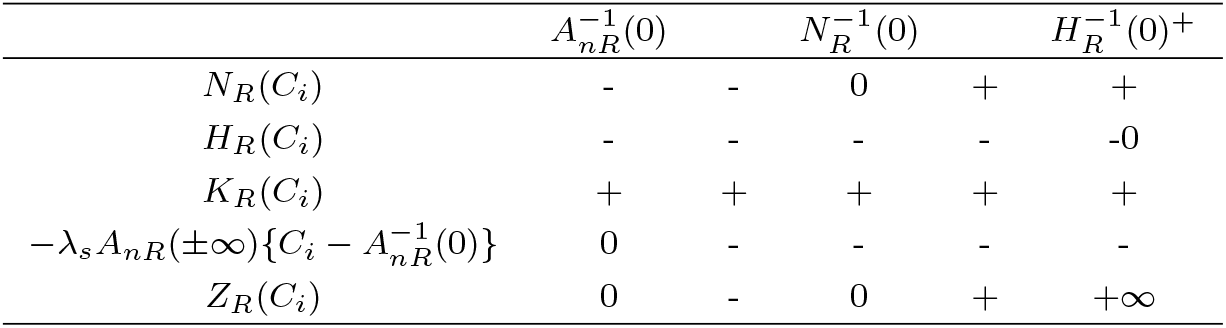
Sign chart of *Z*_*R*_(*C*_*i*_) for [ZI].

**Table SI 46:**
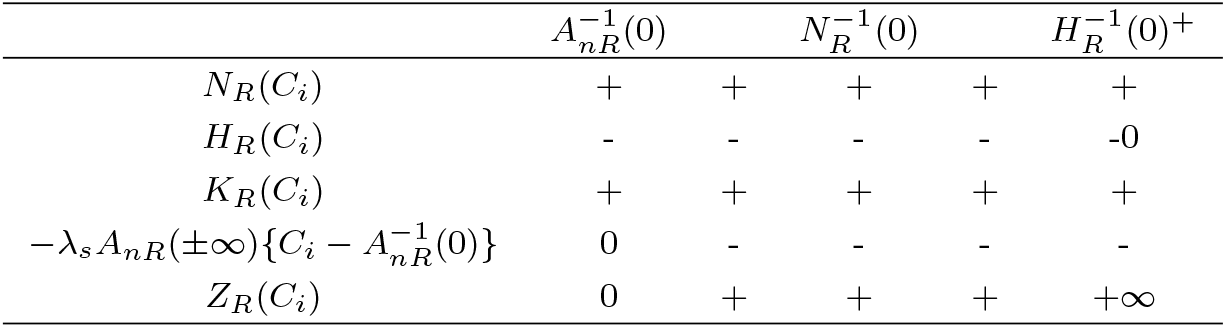
Sign chart of *Z*_*R*_(*C*_*i*_) for [ZII].

#### SI 5.8 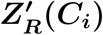

By differentiating Eq. (SI 432) with respect to *C*_*i*_, we have

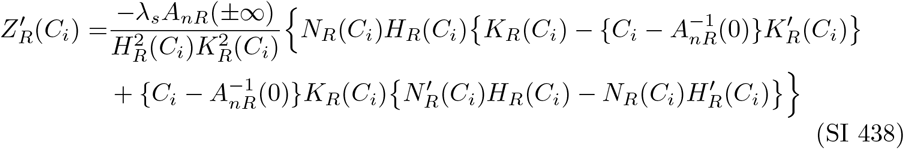

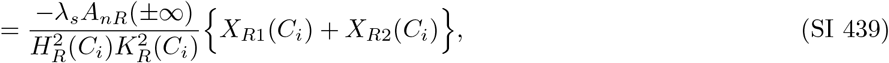

where *X*_*R*1_(*C*_*i*_) and *X*_*R*2_(*C*_*i*_) are given by

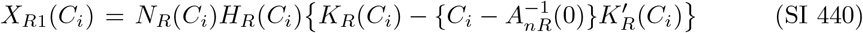

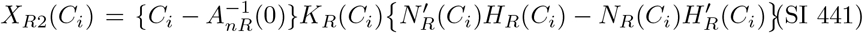

##### SI 5.8.1 *X*_*R*1_(*C*_*i*_)

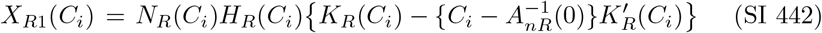

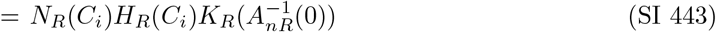

Given that 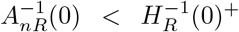 under [C1] (Eq. SI 46), 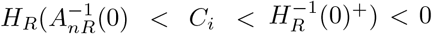 (Eq. (SI 33)), 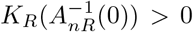 (Eq. (SI 185)), 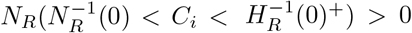 under [ZI] (Eq. (SI 413)), and 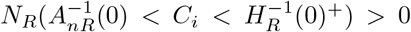 under [ZII] (Eq. (SI 414)), we find that:

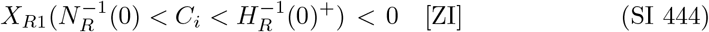

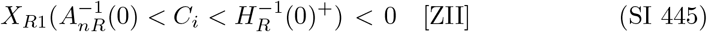

##### SI 5.8.2 *X*_*R*2_(*C*_*i*_)

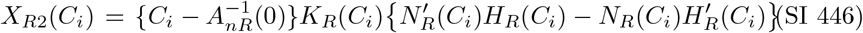

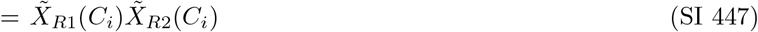

where 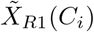 and 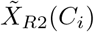 are given by

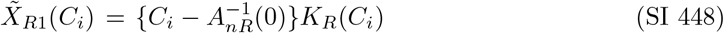

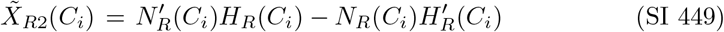

##### SI 5.8.3 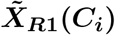

Since 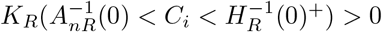 under [C1] (Eq. (SI 182)), Eq. (SI 448) leads to:

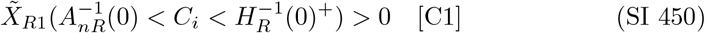

##### SI 5.8.4 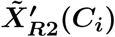

By differentiating 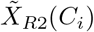 (Eq. (SI 449)) with respect to *C*_*i*_, we obtain:

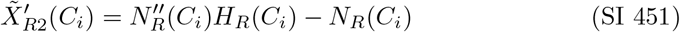

In case of [ZI], since 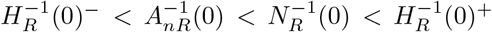 (Eqs. (SI 46) and (SI 415)), 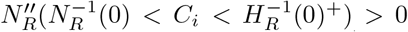 (Eq. (SI 422)), 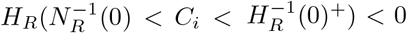 (Eq. (SI 33)), and 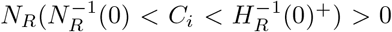 (Eq. (SI 413)), Eq. (SI 451) leads to:

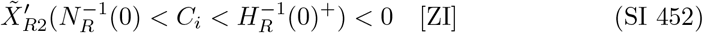

In case of [ZII], since 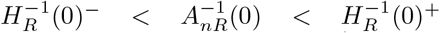 (Eq. (SI 46)),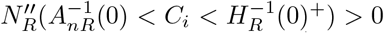 (Eq. (SI 422)), 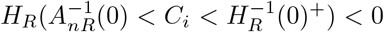 (Eq. (SI 33)), and 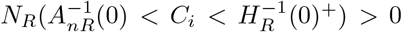 (Eq. (SI 414)), Eq. (SI 451) leads to:

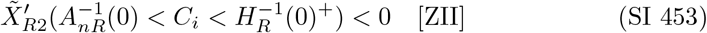

##### SI 5.8.5 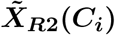

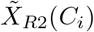 is given from Eq. (SI 449) as:

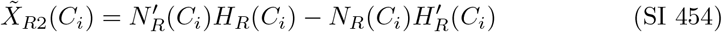

(a) 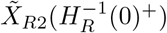

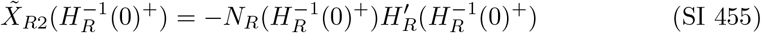 Given that 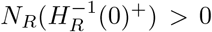 (Eq. (SI 417)) and 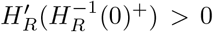 (Eq. (SI 79)), we obtain:

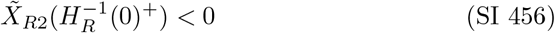
(b) 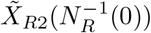 [ZI] is assumed to calculate 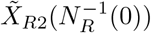,since 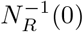 exists under [ZI]. From Eq. (SI 454), 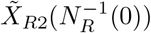 is given by:

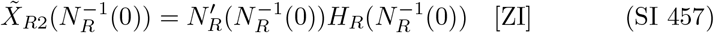 Since 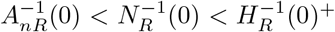 under [ZI] (Eq. (SI 415)), 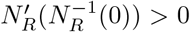 (Eq. (SI 424)) and 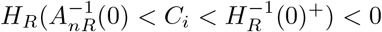 (Eq. (SI 33)), we obtain:

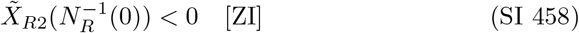
(c) 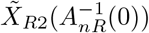 under [ZII]

In this section, [ZII] is assumed to calculate 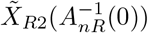.From Eq. (SI 454), 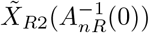 is given by

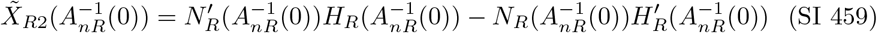

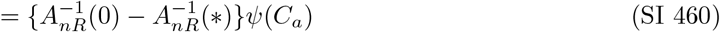

In Eq. (SI 460), we introduce *ψ*(*C*_*a*_), a linear function of *C*_*a*_ given by

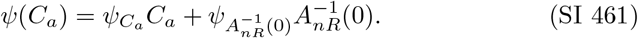

where

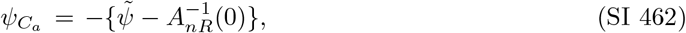

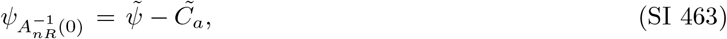

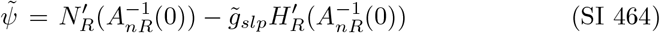

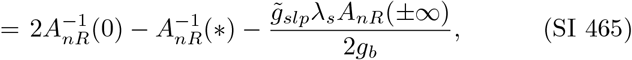

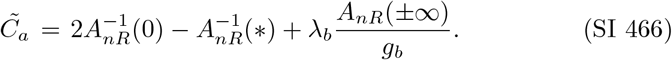

Since 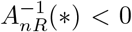 (Eq. (SI 3)) and *A*_*nR*_() *>* 0 under [C1] (Table SI 1), Eq. (SI 466) leads to:

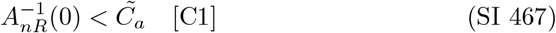

We consider two cases for 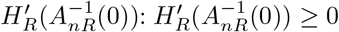 and 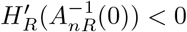.

i. 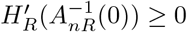 Since 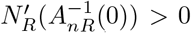 under [ZII] (Eq. (SI 429)), 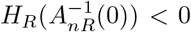 (Eq. (SI 33)), and 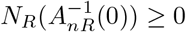 under [ZII] (Eq. (SI 416)), Eq. (SI 459) leads to:

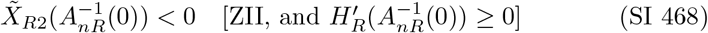
ii. 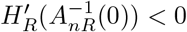

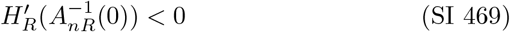

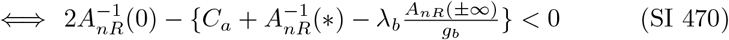

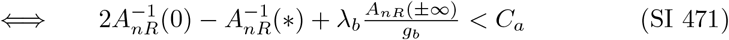

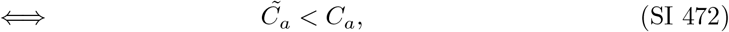

Eq. (SI 472) shows that 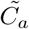 represents the lower limit of *C*_*a*_ for the case 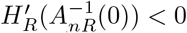.

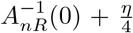 can be represented as a linear function of *C*_*a*_ with a negative slope, as follows:

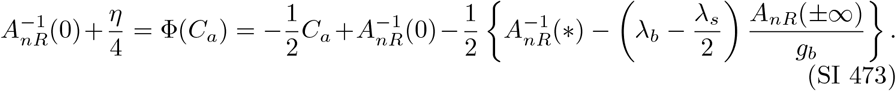

Here Φ(*C*_*a*_) was introduced. 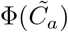 is the maximum under 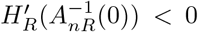, given by:

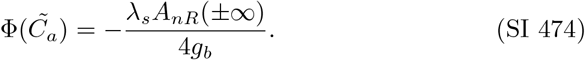

Since *A*_*nR*_(*±*∞) *>* 0 under [C1] (Table SI 1), we have

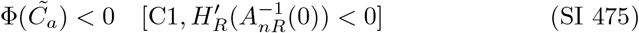

Thus we have:

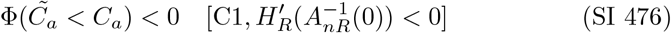

Combining Eqs. (SI 472), (SI 473), and (SI 476), we obtain:

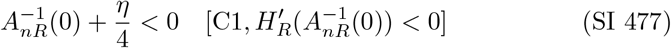

Since sgn 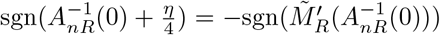 under [C1] (Eq. (SI 252)), Eq. (SI 477) leads to

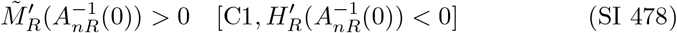

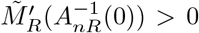 represents the case [P4] under [ZII] (Table SI 41). Thus only [P4] is considered in the case 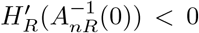 .Since 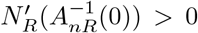 under [P4] (Eq. (SI 429)), Eq. (SI 464) shows:

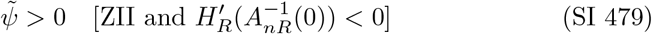

Using Eqs. (SI 462) and (SI 464), we have:

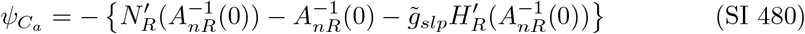

From 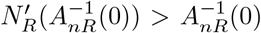 under [ZII] (Proposition 7) and 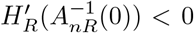, we obtain:

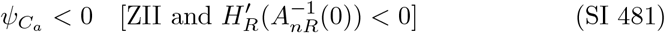

Therefore, the slope of *ψ*(*C*_*a*_) is negative. Hence, 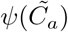 is maximized under [ZII] and 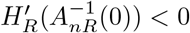 as follows:

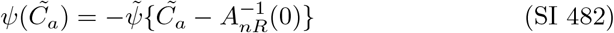

Since 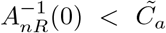 under [C1] (Eq. (SI 467)) and 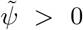 under [ZII] and 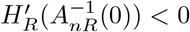 (Eq. (SI 479)), we have:

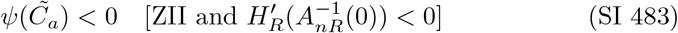

Thus, we obtain:

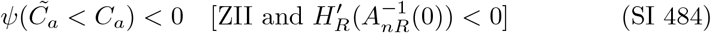

Since 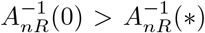 under [C1] (Table SI 1) and 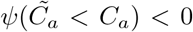 (Eq. (SI 484)), Eq. (SI 460) yields:

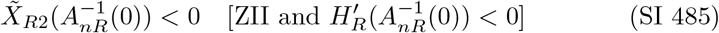

(d) 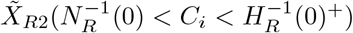 and 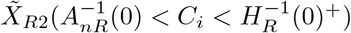 From 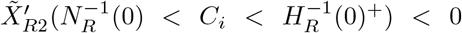 under [ZI] (Eq. (SI 452)) and 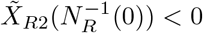 under [ZI] (Eq. (SI 458)), we obtain:

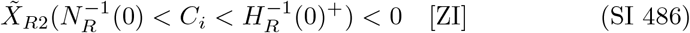 From 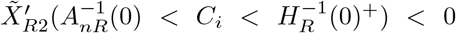 under [ZII] (Eq. (SI 453)) and 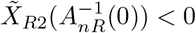 under [ZII] (Eqs. (SI 468) and (SI 485)), we obtain:

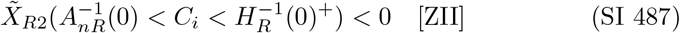

##### SI 5.8.6 Sign of 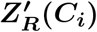

From 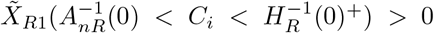 (Eq. (SI 450)), 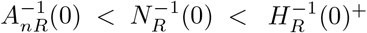 under [Z1] (Eq. (SI 415)), 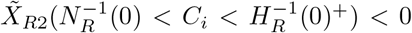 under [ZI] (Eq. (SI 486)), and 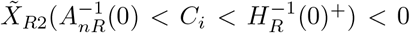 under [ZII] (Eq. (SI 487)), Eq. (SI 447) gives:

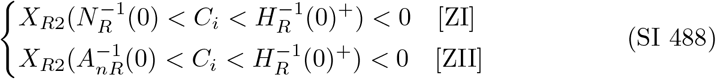

From *A*_*nR*_(*±*∞) *>* 0 (Table SI 1); 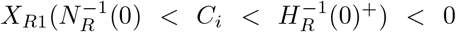 and 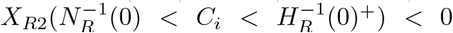 under [ZI] (Eqs. (SI 444) and (SI 488)); 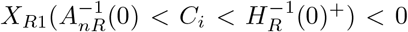 and 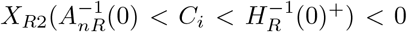 under [ZII] (Eqs. (SI 445) and (SI 488)), Eq. (SI 439) gives:

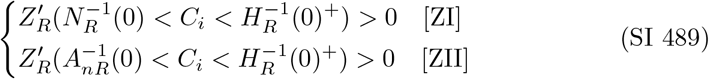

Therefore, *Z*_*R*_(*C*_*i*_) increases monotonically for 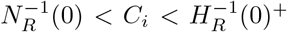 under [ZI], and for 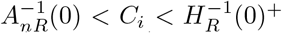 under [ZII]. By using 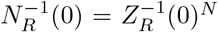 (Eq. (SI 437)), 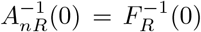 (Eq. (SI 89)), and 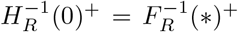 (Eq. (SI 83)), we have:

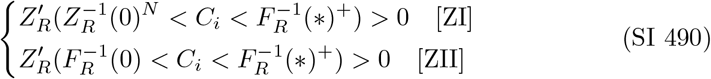

### SI 6 *Z*_*P*_ (*C*_*i*_)

The possible range of appropriate solutions for 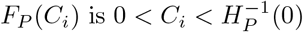 (Table 2).

Therefore, in this chapter, we focus on the range 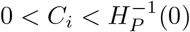.

Using Eq. (21) and Eq. (SI 121), *Z*_*P*_ (*C*_*i*_), defined by *F*_*P*_ (*C*_*i*_) − *G*_*P*_ (*C*_*i*_) (Table 3), can be expressed as:

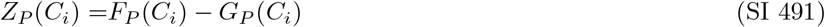

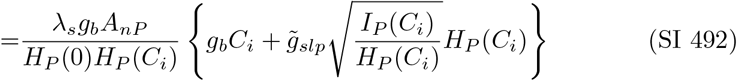

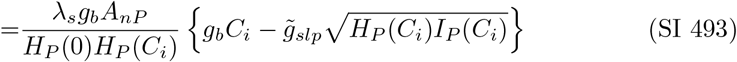

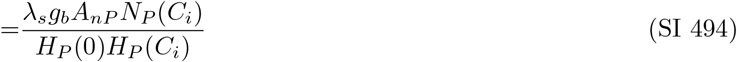

where *I*_*P*_ (*C*_*i*_) and *N*_*P*_ (*C*_*i*_) are given by

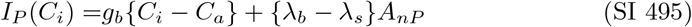

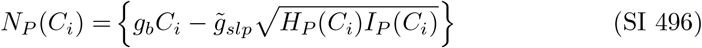

Note that 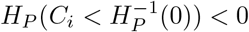 (Eq. (SI 124)) is used to obtain Eq. (SI 493).

#### SI 6.1 *I*_*P*_ (*C*_*i*_)

(a) 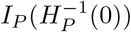

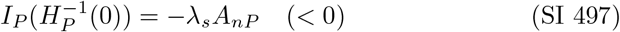 Since *I*_*P*_ (*C*_*i*_) is a linear function of *C*_*i*_ with a positive slope, Eq. (SI 497) leads to:

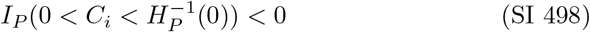

#### SI 6.2 *N*_*P*_ (*C*_*i*_)

(a) *N*_*P*_ (0) and 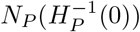

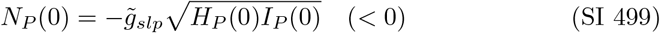

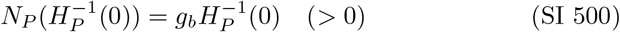 We used 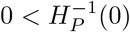 (Eq. (SI 128)) for the inequality in Eq. (SI 500).
(b) 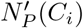 The derivative of *N*_*P*_ (*C*_*i*_) is given by

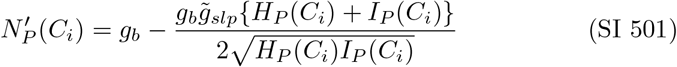 Since 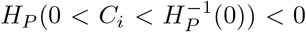 (Eq. (SI 124)) and 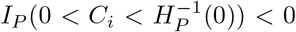 (Eq. (SI 498)), Eq. (SI 501) leads to:

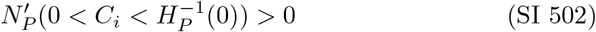
(c) Root: 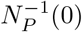

##### Proposition 8.

*Only one root for N*_*P*_ (*C*_*i*_) = 0 *exists at* 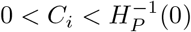.*Proof*. From *N*_*P*_ (0) *<* 0 (Eq. (SI 499)), 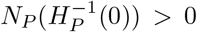 (Eq. (SI 500)), and 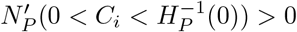 (Eq. (SI 502)), there is only one root for *N*_*P*_ (*C*_*i*_) = 0 at 0 *< C*_*i*_ *< H*^−1^(0). This proves the proposition.

Proposition 8 shows that the root, 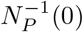,satisfies the following inequality:

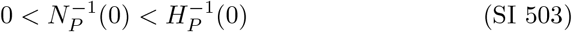

In addition, from *N*_*P*_ (0) *<* 0 (Eq. (SI 499)) and 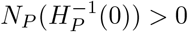 (Eq. (SI 500)), we have

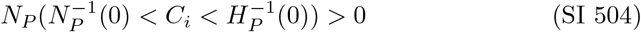

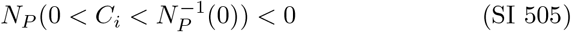

#### SI 6.3 *Z*_*P*_ (*C*_*i*_)

*Z*_*P*_ (*C*_*i*_) can be rewritten from Eq. (SI 494) as follows:

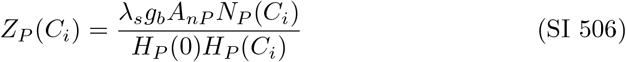

(a) Singularity: 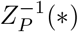 Eq. (SI 506) shows that the singularity of *Z*_*P*_ (*C*_*i*_) is given by

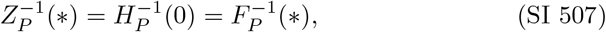

where 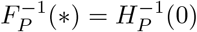 (Eq. (SI 133)) is used.
(b) Root: 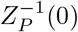 Eq. (SI 506) shows that the roots of *Z*_*P*_ (*C*_*i*_) are the same as those of *N*_*P*_ (*C*_*i*_), as follows:

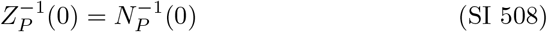 Proposition 8 shows that only one 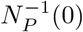 exists at 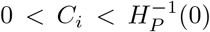.Since 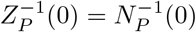 (Eq. (SI 508)), there is only one 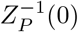 in the range 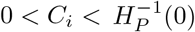.

Given that *A*_*nP*_ *>* 0 (Eq. (15)), *H*_*P*_ (0) *<* 0 (Eq. (SI 130)), 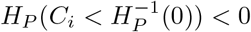 (Eq. (SI 124)), and 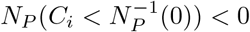 (Eq. (SI 505)), Eq. (SI 506) implies:

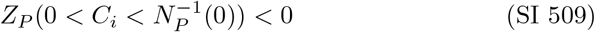

### SI 6.4 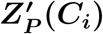

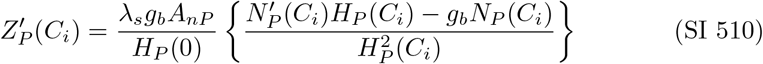

From *A*_*nP*_ *>* 0 (Eq. (15)), *H*_*P*_ (0) *<* 0 (Eq. (SI 130)), 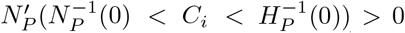 (Eq. (SI 502)), 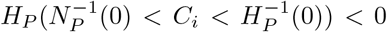 (Eq. (SI 124)), and 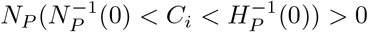 (Eq. (SI 504)), Eq. (SI 510) implies:

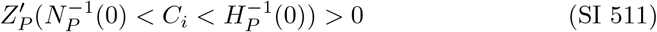

By using 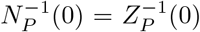 (Eq. (SI 508)) and 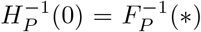 (Eq. (SI 507)), it follows from Eq. (SI 511):

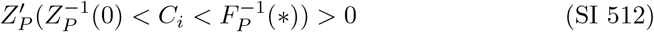

